# Dynamics of transposable element accumulation in the non-recombining regions of mating-type chromosomes in anther-smut fungi

**DOI:** 10.1101/2022.08.03.502670

**Authors:** Marine Duhamel, Michael E. Hood, Ricardo C. Rodríguez de la Vega, Tatiana Giraud

## Abstract

**Background:** Transposable element (TE) activity is generally deleterious to the host fitness, such that TE copies are often purged by selection, which is facilitated when meiotic recombination reshuffles variation among loci. In the absence of recombination, the number of TE insertions usually increases, but the dynamics of such TE accumulations is unknown.

**Results:** In this study, we investigated the temporal dynamics of TE accumulation in the non-recombining genomic regions of 15 *Microbotryum* species, leveraging on a unique dataset of 21 independent evolutionary strata of recombination cessation of different ages. We show that TEs rapidly accumulated in regions lacking recombination, but that the TE content reached a plateau at ca. 50% of occupied base pairs by 1.5 MY following recombination suppression. The same TE superfamilies have repeatedly expanded in independently evolved non-recombining regions, in particular rolling-circle replication elements (*Helitrons*), despite being scarce before recombination suppression. The most abundant elements, long-terminal repeat (LTR) retrotransposons of the *Copia* and *Ty3* superfamilies, expanded through transposition bursts affecting both the non-recombining regions of mating-type chromosomes and autosomes, thus suggesting that non-recombining regions constitute a reservoir of TEs that transpose to recombining regions. Based on LTR sequence divergence within and among copies, we could distinguish bursts of transposition from gene conversion.

**Conclusion:** Our study supports the TE reservoir hypothesis, by showing that TE accumulation in non-recombining has a genome-wide impact. TEs accumulated through bursts, and following a non-linear, decelerating dynamics, thus improving our knowledge on genome evolution, particularly in association with sex or mating-type chromosomes.

## Background

Transposable elements (TEs) are autonomously replicating DNA sequences ubiquitous in eukaryotic genomes (Hua-Van *et al*., 2005; Wicker *et al*., 2007; Bourque *et al*., 2018). TEs have occasionally been recruited by their host genomes, *e.g.,* as regulatory sequences, for centromere maintenance or other functions (Kaessmann *et al*., 2009; Martin *et al*., 2009; Hof *et al*., 2016; Dallery *et al*., 2017; Lanciano & Mirouze, 2018). However, TE replication and insertion are generally deleterious to their host genome (Payer & Burns, 2019; Andrenacci *et al*., 2020; Burns, 2020). Their insertions can indeed disrupt gene coding or regulatory sequences (Hancks & Kazazian, 2016), induce epigenetic changes (Castanera *et al*., 2016; Choi & Lee, 2020) or promote chromosomal rearrangements via ectopic recombination (Lim & Simmons, 1994; Han *et al*., 2008; Ade *et al*., 2013; Bennetzen & Wang, 2014; Balachandran *et al*., 2022). TEs can also incur a substantial replication cost as they can represent a large proportion of the genome (Schnable *et al*., 2009). TE proliferation is therefore often controlled by the host genome, by inactivating TE copies through epigenetic mechanisms such as RNAi silencing (Saito & Siomi, 2010) and DNA methylation (Zemach *et al*., 2010; Jones, 2012; Bewick *et al*., 2019; Deniz *et al*., 2019).

Deleterious TE insertions can also be purged *a posteriori* by recombination between homologous chromosomes carrying different TE insertions. In the absence of recombination, the average number of TE insertions in the population increases with time (Dolgin & Charlesworth, 2008). TEs therefore tend to accumulate in regions with lower recombination rates (Kent *et al*., 2017; Rifkin *et al*., 2022), particularly in non-recombining regions such as in centromeres (Plohl *et al*., 2014), pericentromeric regions (Wright *et al*., 2003; Peng & Karpen, 2008; Charlesworth & Campos, 2014), and the non-recombining regions of sex chromosomes in plants and animals (Bachtrog, 2003, 2005; Marais *et al*., 2008; Ahmed *et al*., 2014) and of mating-type chromosomes in fungi (Lengeler *et al*., 2002; Bakkeren *et al*., 2006; Badouin *et al*., 2015; Branco *et al*., 2018). The TE copies accumulating in the non-recombining regions of sex and mating-type chromosomes may form a reservoir for the rest of the genome, with TE copies transposing from these regions onto autosomes when epigenetic defenses are insufficient (Hood *et al*., 2004; Wei *et al*., 2020; Peona *et al*., 2021).

The accumulation of TEs has been extensively documented, but the dynamics of their proliferation in non-recombining regions, and whether their accumulation in non-recombining regions has genome-wide impacts as a source of TEs, are unknown (Charlesworth, 2021). Comparing TE proportions in non-recombining regions of different ages in closely related species would allow assessing the dynamics of TE accumulation, *i.e.*, the rate at which TEs increase in copy number in the absence of recombination. The cessation of recombination leads to degeneration in a variety of ways, which can have non-linear, decelerating dynamics as suggested for gene losses on theoretical grounds (Bachtrog, 2013; Charlesworth, 2021) and for optimal codon usage as shown by genomic data (Carpentier *et al*., 2022). Alternatively, other components of degeneration can have a linear dynamics of accumulation, as shown for changes in protein sequences (Carpentier *et al*., 2022). As TEs are a major degeneration factor, it is important to assess whether TEs accumulate rapidly in young non-recombining regions, whether the accumulation slows down over time, for example because of the evolution of more efficient control mechanisms, whether the number of TE copies grows exponentially out of control, versus linearly or by occasional bursts of proliferation. The curve shape for TE accumulation could thus inform our understanding of how their proliferation is controlled in the absence of recombination. In addition, comparing the genealogies of TE copies within the genomes using sequence identity and genomic localisation would allow testing whether TE insertion events following recombination suppression are restricted to the non-recombining regions or whether TE copies from non-recombining regions can transpose to the rest of the genome, thus having a genome-wide impact.

Chromosomes carrying the genes for reproductive compatibility are particularly interesting and useful models to address the temporal dynamics of TE accumulation in non-recombining regions. Closely related species can have independently acquired sex chromosomes with non-recombining regions of different sizes and ages, which have often extended in a stepwise manner, providing independent events of recombination suppression within and across species (Lahn & Page, 1999; Bergero & Charlesworth, 2009; Charlesworth, 2017). Such stepwise extensions of recombination suppression produce evolutionary strata of different levels of genetic differentiation between sex chromosomes, as first observed on the human sex chromosomes (Lahn & Page, 1999), and later on various sex chromosomes in other animals and plants (Handley *et al*., 2004; Bergero & Charlesworth, 2009).

The accumulation of TEs has even been suggested to play a role in the extension of recombination cessation along sex chromosomes (Li *et al*., 2016; Kent *et al*., 2017). TE-induced loss-of-function mutations constitute recessive deleterious mutations in the recombining regions adjacent to the non-recombining zone, called pseudo-autosomal regions. The segregation of deleterious mutations near non-recombining regions can foster the expansion of recombination suppression. For example, inversions carrying fewer deleterious mutations than average are selected when rare but suffer from their exposed genetic load when frequent enough to occur as homozygotes; if such less-loaded inversions occur near a permanently heterozygous allele, such as near mating-type loci or Y-like sex-determining alleles, they remain heterozygous and can reach maximal frequencies, thereby expanding the non-recombining region (Jay *et al*., 2022; Tezenas *et al*., 2022). TEs can themselves promote inversions at the margin of non-recombining regions (Ironside, 2010; Zhang *et al*., 2011). Additionally, epigenetic modifications that silence TEs, such as DNA methylation and heterochromatin formation, can also in themselves block recombination (Maloisel & Rossignol, 1998; Ben-Aroya *et al*., 2004; Yelina *et al*., 2012, 2015; Bachtrog, 2013; Li *et al*., 2016). If TEs accumulate in the margin of non-recombining regions, they may thus drive the extension of recombination suppression, either as ultimate or proximate causes, or both (Li *et al*., 2016; Kent *et al*., 2017). Similarly to sex chromosomes, fungal mating-type chromosomes can display large non-recombining regions, with evolutionary strata of different ages as well as footprints of degeneration (Menkis *et al*., 2008; Whittle *et al*., 2011; Fontanillas *et al*., 2015; Idnurm *et al*., 2015; Branco *et al*., 2018; Hartmann *et al*., 2021; Carpentier *et al*., 2022). Fungal mating-type chromosomes are particularly useful models for studying sex-related chromosome evolution as they often display young non-recombining regions, with many independent events of recombination suppression, which allows studying early stages of degeneration (Whittle *et al*., 2011; Fontanillas *et al*., 2015; Carpentier *et al*., 2022). Fungi with evolutionary strata are most often automictic, i.e. undergoing intra-tetrad selfing (Hartmann *et al*., 2020; Vittorelli *et al*., 2022). Theoretical models have suggested that this may be because automictic fungi are dikaryotic for most of their lifecycle, even in ascomycetes, thus allowing the sheltering of deleterious alleles (Jay *et al*., 2022), and because automixis can prevent diploid selfing to purge deleterious mutations near mating-type loci (Antonovics & Abrams, 2004; Tezenas *et al*., 2022). When non-recombining regions occur on fungal mating-type chromosomes, they are relatively young and some genera display multiple independent events of recombination cessation across closely related species, allowing to study the early stages of genomic degenerescence with independent data points (Carpentier *et al*., 2022). In addition, fungi generally have small genomes and can be cultivated at the haploid phase, facilitating genomic assemblies and comparisons. The reduced TE load in fungal genomes compared to plants and animals (Amselem *et al*., 2015) further facilitates assemblies of their chromosomes, including the TE-enriched non-recombining regions (Badouin *et al*., 2015; Branco *et al*., 2018). Fungal defenses against TE activity include several forms of RNAi-mediated post-transcriptional inactivation and heterochromatic silencing (Gladyshev, 2017), in particular DNA methylation (Castanera *et al*., 2016; Bewick *et al*., 2019) and targeted C-to-T mutation in repetitive sequences, such as repeat-induced point mutation (RIP), targeting specific di- or tri-nucleotides (Galagan & Selker, 2004; Hood *et al*., 2005; Amselem *et al*., 2015).

*Microbotryum* fungi form a complex of closely related plant-castrating species, mostly infecting Caryophyllaceae plants, and multiple species have independently evolved non-recombining mating-type chromosomes (Branco *et al*., 2017, 2018; Carpentier *et al*., 2019; Duhamel *et al*., 2022). In *Microbotryum* fungi, as in most other basidiomycetes, mating only occurs between haploid cells carrying alternative alleles at two mating-type loci: the pheromone-receptor (PR) locus controlling gamete fusion, encompassing genes coding for a pheromone-receptor and pheromones, and the homeodomain (HD) locus controlling dikaryotic hyphal growth, encompassing two homeodomain genes (Kronstad & Staben, 1997). Most *Microbotryum* fungi display only two mating types, called a_1_ and a_2_ (Day, 1979), which is due to multiple independent events of recombination suppression having linked the two mating-type loci across the *Microbotryum* genus (Badouin *et al*., 2015; Branco *et al*., 2017, 2018; Duhamel *et al*., 2022; Carpentier *et al*., 2022) (Figure 1); a few of the studied *Microbotryum* species still have unliked mating-type loci, but often with linkage of each mating-type locus to its centromere (Carpentier *et al*., 2019) (Figure 1). These two types of genomic linkage, *i.e.,* of PR and HD mating-type loci to each other or to their centromeres, are beneficial under intra-tetrad mating (automixis), as they increase the odds of compatibility among gametes of a single meiosis during automixis (Carpentier *et al*., 2019), the common mating system in *Microbotryum* fungi (Hood & Antonovics, 2000; Giraud *et al*., 2008).

**Figure 1:**
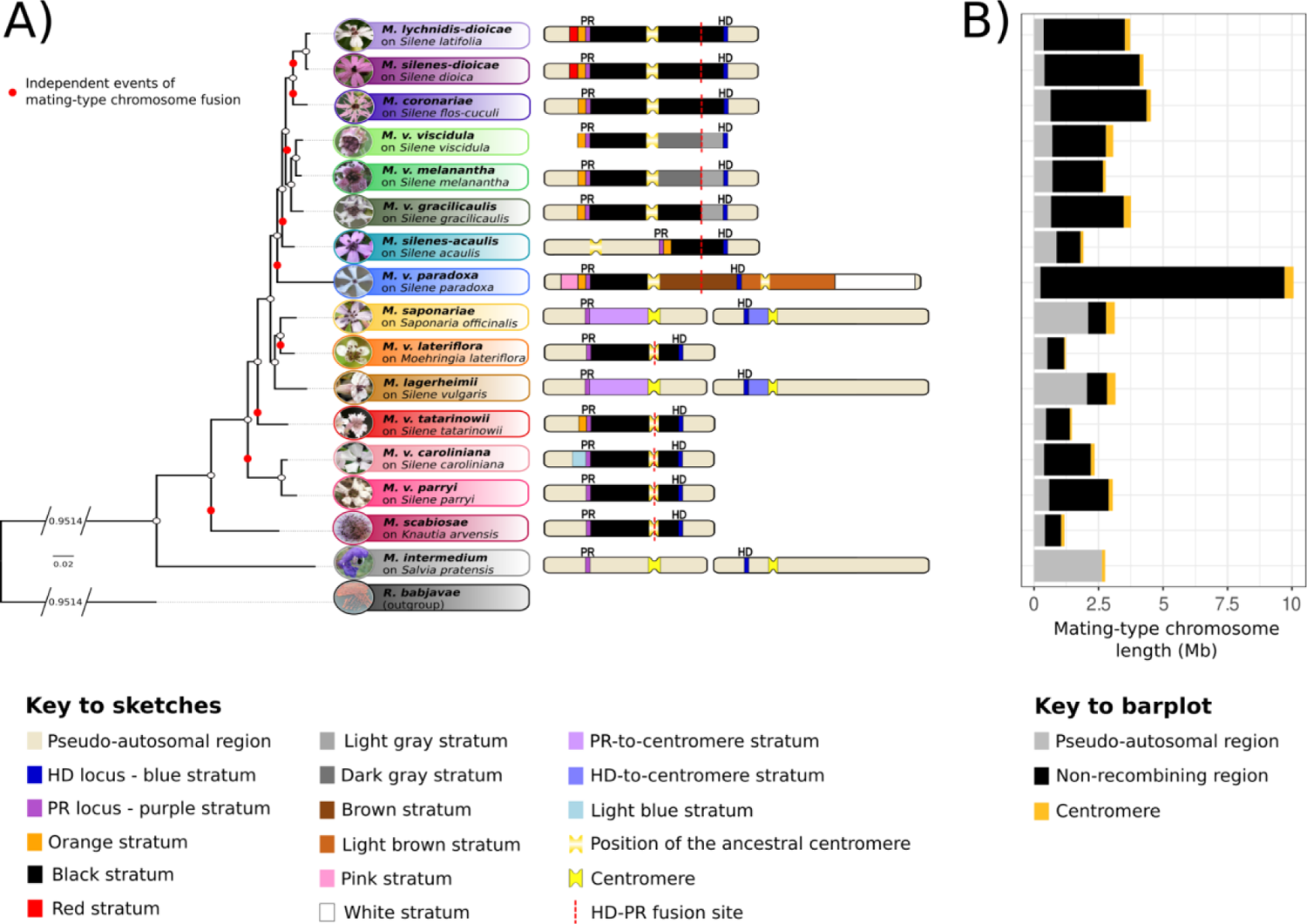
Multiple evolutionary strata formed by independent events of mating-type locus linkage and stepwise extension of recombination suppression in *Microbotryum* fungi. A) The mating-type chromosomes of *Microbotryum* species with linked mating-type loci have had many rearrangements but are depicted here according to the gene order at the moment of the fusion between the homeodomain (HD) and pheromone-receptor (PR) chromosomes, the fusion site being indicated by a dashed red line. The ancestral gene order was inferred from *Microbotryum lagerheimii* and *M. intermedium* which have unlinked mating-type loci and collinear mating-type chromosomes. The old blue and purple strata have evolved before diversification of the *Microbotryum* clade. In *M. saponariae* and *M. lagerheimii*, recombination cessation linked each mating-type locus to their respective centromeres, forming the HD- and PR-to-centromere strata. The black strata correspond to the events of recombination suppression having linked the two mating-type loci, which occurred independently at least nine times in *Microbotryum*, indicated by red dots on the phylogeny. In some species (*M. violaceum viscidula*, *M. v. melanantha*, *M. v. gracilicaulis)*, recombination between the mating-type loci was gradually suppressed following the chromosomal fusion event, and formed several strata called black, light gray and dark gray strata. In *M. v. paradoxa*, recombination was partially restored by introgression before being suppressed again, forming the brown stratum. In many species, recombination cessation further extended beyond the mating-type loci, capturing different sets of genes and forming strata that were named as indicated in the figure key. **B)** Average (a_1_-a_2_) size of the mating-type chromosomes in *Microbotryum* fungi. The average (a_1_-a_2_) size of the pseudo-autosomal region, non-recombining region and centromere is represented in gray, black and yellow, respectively.

Furthermore, patterns of decreasing divergence between the alleles on alternative mating-type chromosomes when farther from the mating-type loci in the ancestral gene order have revealed that recombination suppression extended in a stepwise manner beyond mating-type loci in several *Microbotryum* fungi, forming evolutionary strata of different ages (Branco *et al*., 2017, 2018; Duhamel *et al*., 2022; Carpentier *et al*., 2022) (Figure 1 and Supplementary Table 1). In contrast, the recombining regions of the mating-type chromosomes, called pseudo-autosomal regions, are collinear and highly homozygous in *Microbotryum* fungi (Branco *et al*., 2017, 2018; Duhamel *et al*., 2022). The non-recombining regions of the mating-type chromosomes have accumulated TEs compared to the pseudo-autosomal regions and the autosomes (Fontanillas *et al*., 2015; Branco *et al*., 2018). However, whether specific TE superfamilies have accumulated and what is the temporal dynamics of TE accumulation across these genomic compartments remain unresolved questions.

At the whole-genome scale, *Microbotryum lychnidis-dioicae* carries long-terminal repeat (LTR) retroelements from the *Copia* and *Ty3* (also known as *Gypsy*, but see Wei *et al*., 2022) superfamilies (Hood *et al*., 2005; Amselem *et al*., 2015) as well as, to a lesser extent, *Helitron* DNA transposons, *i.e.*, rolling-circle replicating TEs (Kapitonov & Jurka, 2001). In the *Microbotryum* species parasitizing *Dianthus carthusianorum*, a burst of *Ty3* retroelements has caused a massive accumulation of this element in a short period of time, in spite of RIP-like activity (Horns *et al*., 2017). In *Microbotryum* genomes, a RIP-like activity targets TCG trinucleotides (CGA on the reverse complement strand) rather than di-nucleotides as in ascomycetes (Hood *et al*., 2005; Johnson *et al*., 2010; Horns *et al*., 2012).

In this study, we investigated the temporal dynamics of TE accumulation in the non-recombining regions of 15 *Microbotryum* species, leveraging on the existence of 21 independent evolutionary strata of different ages, within and across species (Carpentier *et al*., 2022) (Figure 1). We show that TEs rapidly accumulate in young non-recombining regions but that their abundance reached a plateau by 1.5 MY following recombination suppression at ca. 50% occupied base pairs. *Helitron* DNA transposons repeatedly expanded in non-recombining regions despite being rare in species with recombining mating-type chromosomes. The most abundant TE superfamilies, *i.e.*, *Copia* and *Ty3*, were also over-represented in non-recombining regions and accumulated through multiple bursts that impacted both the non-recombining regions of the mating-type chromosomes and the autosomes at the same time, supporting the TE reservoir hypothesis. TEs, however, did not preferentially accumulate at the margin of non-recombining regions.

## Results

### Transposable elements in *Microbotryum* genomes

We identified transposable elements (TEs) in the a_1_ and a_2_ haploid genomes deriving from a single diploid individual of each of 15 *Microbotryum* species and of a genome of the red yeast *Rhodothorula babjavae*, an outgroup with the ancestral condition of the PR and HD mating-type loci on separate chromosomes. We detected no significant difference in TE content between the two haploid genomes within species (Wilcoxon paired test, V=5284, *p-value*=0.208, Supplementary Figure S1). In the following analyses, we therefore calculated the mean from the a_1_ and a_2_ genomes for each species. *Microbotryum* genomes, where there was an average of 18.3% base pairs occupied by TEs, displayed a higher density of TEs than in *R. babjavae* (Supplementary Figure S1). *Microbotryum intermedium*, with fully recombining mating-type chromosomes, had the lowest TE content (6.57%), closely followed by *M. v. lateriflora* (7.02%) with a very young non-recombining region (0.304 My, Duhamel *et al*., 2022), while *M. violaceum paradoxa* with the largest and among the oldest non-recombining regions had the highest TE load (30.33%) (Figure 2a and Supplementary Figure S2 and Supplementary Table 2; detailed comparison below).

**Figure 2:**
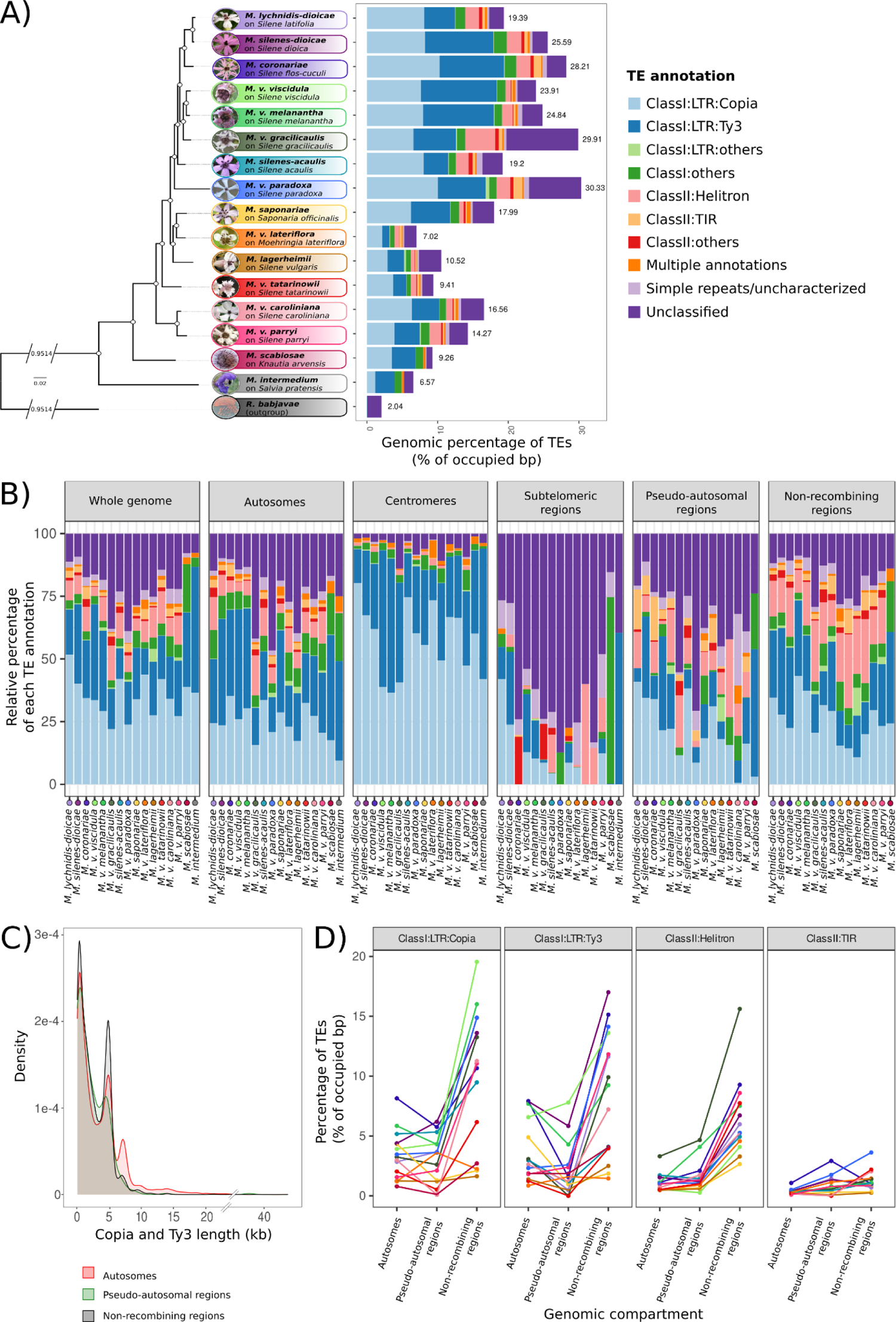
Transposable element (TE) composition of *Microbotryum* genomes. **A)** Whole-genome TE load (as percentage of occupied base pairs) across the *Microbotryum* phylogeny, each color representing a TE category. Numeric values correspond to the a_1_-a_2_ average. **B)** Relative proportions of the various TE categories in the different genomic compartments of *Microbotryum* genomes. Each color represents a TE category, as in A). **C)** Distribution of the length of *Copia* and *Ty3* retroelements in the autosomes, the pseudo-autosomal regions and the non-recombining regions of *Microbotryum* fungi, in red, green and black, respectively. **D)** TE load per genomic compartment, *i.e.*, autosomes (without the non-recombining centromeres), pseudo-autosomal regions and non-recombining regions for the main TE categories (*Copia, Ty3, Helitron,* terminal inverted repeats - TIR). The lines connect the values in the same species, to facilitate comparisons. Each color represents a *Microbotryum* species.

All *Microbotryum* genomes contained a substantial fraction of long-terminal repeat (LTR) retrotransposons from the *Copia* and *Ty3* superfamilies (6.04% and 4.97% of occupied base pairs on average for *Copia* and *Ty3* retroelements, respectively, Supplementary Table 4). These retroelements were particularly abundant in the *Microbotryum* genomes with non-recombining mating-type chromosomes, *Copia* TEs being for example ten times as abundant in *M. coronariae* (10.27% of occupied base pairs) as in *M. intermedium* (1.08% of occupied base pairs). Taken together, *Copia* and *Ty3* superfamilies represented 76.93 to 94.01% of the centromeric TEs across species (Figure 2B). The trimodal distributions of *Copia* and *Ty3* retroelement length (Figure 2C and Supplementary Figure S3a) show that, while most copies are shorter than 1 kb and are likely degenerated and fragmented, a high proportion of these retroelements, being 5 kb and 7 kb length on average, are likely intact TE copies (González & Deyholos, 2012). TE copies with a length greater than 10 kb likely represent nested TE copies and were particularly abundant in the young stratum of *M. v. paradoxa,* called the white stratum (Supplementary Figure S3a).

Because retrotransposons such as LTRs can be common in plants (Piegu *et al*., 2006; González & Deyholos, 2012), we investigated whether horizontal transfers of TEs could have occurred between the host plant and *Microbotryum* genomes. We compared the sequences from the plant Repbase database to the annotated TE sequences in *Microbotryum* genomes by blast. We obtained several BLAST hits with high identity and low e-value scores, but none exceeded 6% of the annotated TE sequence lengths (Supplementary Table 3). Therefore, no putative inter-kingdom horizontal TE transfer was identified.

DNA transposons were mostly represented by *Helitrons* and terminal inverted repeats (TIR) elements in most *Microbotryum* genomes (1.44 and 0.42% of occupied base pairs on average, respectively), while these were nearly absent in *M. intermedium* (0.04% of base pairs occupied by *Helitrons* and no TIR elements) and in *M. scabiosae* (Figure 2a). The proportion of TEs annotated as “*unclassified*” was generally low (0.83 to 3.25% of occupied base pairs), being highest in genomes with high TE content, such as *M. v. paradoxa* and *M. v. gracilicaulis* (7.4 and 10.18% of occupied base pairs, respectively). The low average size of “*unclassified*” TE copies (263.13 bp, Supplementary Figure S3b) suggests that they represent highly fragmented TE relics. Such “*unclassified*” elements were predominant in subtelomeric regions (Figure 2B).

### Higher content of transposable element and of RIP-like footprints in non-recombining regions

TEs represented significantly higher proportions of base pairs in non-recombining regions compared to pseudo-autosomal regions and autosomes (ANOVA post-hoc Tukey test, adj. *p-value* < 0.01), being on average 41.68% and 40.31% higher in the non-recombining regions compared to pseudo-autosomal regions and autosomes, respectively (Supplementary Table 4, Supplementary Figure S4). No significant difference was observed between pseudo-autosomal regions and autosomes (ANOVA post-hoc Tukey test, adj. *p-value* = 0.922, Supplementary Table 4), despite tendencies in some species for lower content of some elements in pseudo-autosomal regions (Figure 2C and Supplementary Figure S4). The centromeres of all species displayed a high density of TEs (73% of base pairs occupied by TEs on average, Supplementary Table 5). Plotting the density of TEs along mating-type chromosomes further illustrated the higher values in the non-recombining regions than in the pseudo-autosomal regions, and that the TE-rich regions corresponded to gene-poor regions (see Figure 3 for *M. lychnidis-dioicae* and Supplementary Figure S5A to S5O for the other *Microbotryum* species).

**Figure 3:**
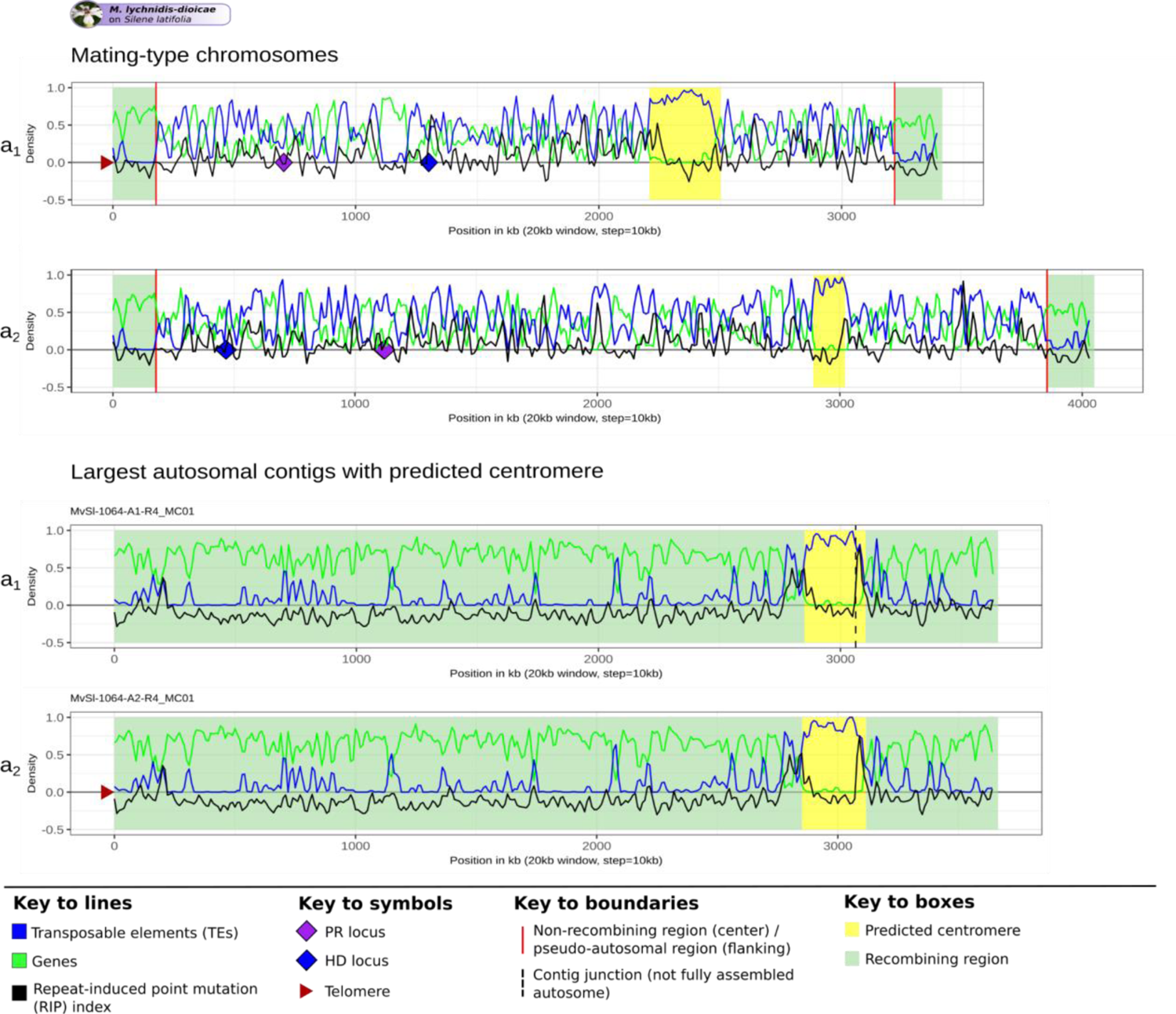
Density of transposable elements (TEs), genes and repeat-induced point mutation (RIP) index along the mating-type chromosome and the autosomes of *Microbotryum lychnidis-dioicae*. Density of TEs, genes and RIP-like index were calculated in 20 kb non-overlapping windows and plotted along the mating-type chromosomes and the largest autosomes, using the same X scale. A RIP-like index greater than zero indicates that the region is affected by a mechanism leading to C to T transitions resembling RIP. Predicted centromeres are indicated in yellow. Recombining regions of the mating-type chromosomes (pseudo-autosomal regions) and an autosome (the largest autosomal contig in the genome harboring a predicted centromere) are highlighted in pale green. The non-recombining region is delimited by red lines. Pheromone-receptor (PR) and homeodomain (HD) loci are indicated by purple and blue diamonds, respectively. Identified telomeres are indicated by brown triangles. The plots for the other species are shown in Figure S5.

TE density was generally higher in older evolutionary strata such as the stratum around the PR locus, called the purple stratum (Supplementary Figure S5H, S5J and S5O), and lower in young strata such as the pink and white strata of *M. v. paradoxa* (Supplementary Figure S5G). In part because of this variation in TE content related to age of the non-recombining region, the size of the mating-type chromosomes varied greatly across *Microbotryum* species (Figure 1B), from 1,177,022 bp in *M. scabiosae* to 10,060,889 bp in *M. v. paradoxa* (3,076,521 bp on average); the size of the mating-type chromosomes also varies because different chromosomal rearrangements were involved in the origin of HD-PR locus linkage (Duhamel *et al*., 2022). For the same reasons, the size of the non-recombining regions was variable, ranging from 630,774 bp in *M. scabiosae* to 9,461,162 bp in *M. v. paradoxa*, thus spanning almost the full length of the mating-type chromosome in this later species (Figure 1B), while being absent in *M. intermedium*.

We found no evidence that TEs have particularly accumulated in the flanking regions of the non-recombining regions when comparing the proportion of base pairs occupied by TEs within the 100 kb of the pseudo-autosomal regions flanking the non-recombining regions to the distribution of the TE proportions in regions of the same size in fully recombining autosomes (Supplementary Figure S6). Indeed, the probability of having a greater TE load in a 100 kb window on autosomes than at the margin of the non-recombining region was ≤ 0.05 in a single species (Supplementary Figure S6). We compared the TE content in the flanking region of the non-recombining region to that in autosomes because the pseudo-autosomal regions were too small for building a distribution.

The non-recombining regions had more RIP-like signatures in TEs than autosomes and pseudo-autosomal regions in most species (Figure 3 and Supplementary Figure S5). The youngest strata, such as the pink and white strata of *M. v. paradoxa* and the young light blue stratum in *M. v. caroliniana,* were in contrast less affected by RIP in TEs than the older non-recombining regions (Supplementary Figure S5G and S5L, respectively).

### Particular TE family expansions in non-recombining regions

Some TE families have particularly expanded in non-recombining regions. *Helitron* elements, for example, have expanded in non-recombining regions, while they are relatively rare in the recombining mating-type chromosomes of *M. intermedium* and in autosomes in all species (ANOVA post-hoc Tukey test, adj. *p-value* < 0.01, see Supplementary Table 4 and Figure 2B and 2C). TEs of the *Copia* and *Ty3* superfamilies displayed higher proportions in the non-recombining regions compared to autosomes and pseudo-autosomal regions in almost all species (ANOVA post-hoc Tukey test, adj. *p-value* < 0.01, see Figure 2C and Supplementary Table 4). The content of *Helitrons*, *Copia* and *Ty3* elements increased on average by 5.42, 6.56, and 4.77% in non-recombining regions compared to autosomes, while the content in other TEs increased by 1.48% in non-recombining regions on average across species. The proportion of *TIR* DNA transposons was higher in the non-recombining regions compared to pseudo-autosomal regions but not compared to autosomes (ANOVA post-hoc Tukey test, adj. *p-value* < 0.01, Supplementary Table 4 and Figure 2C).

### Genome-wide bursts of transposable elements following recombination suppression

We found a significant and strongly positive correlation between the proportions of base pairs occupied by TEs on autosomes and non-recombining regions (r = 0.67; *p-value* = 0.006; Supplementary Figure S7), which is consistent with the reservoir hypothesis, *i.e.*, that the TE copies accumulating in non-recombining regions have a genome-wide impact by transposing to autosomes.

We inferred TE genealogies of the *Copia* and *Ty3* superfamilies independently, using the alignment of their 5’-LTR sequences. Because the genealogies rely on the identification of the 5’-LTR sequences of the retrotransposons, only a subset of the *Copia* and *Ty3* retroelements were used to reconstruct the genealogies (on average 12 and 9.65% of the annotated *Copia* and *Ty3* superfamily retroelements, respectively, Supplementary Figure S8) and genealogies were built for the species having sufficiently large sets of retained *Copia* and *Ty3* sequences.

The *Copia* and *Ty3* genealogies revealed the existence of many copies clustering within large clades having very low divergence, suggesting multiple bursts of transposition of these retroelements that impacted all genomic compartments (Figure 4 and additional species shown in Supplementary Figures S9 and S10), including in the species with old non-recombining regions (Figure 4B and Supplementary Figures S9G-H and S10B). Clusters of similar TE copies could however not only be recent bursts of transpositions (that we define here as a sudden accumulation of copies, resulting from either an increase in intrinsic TE activity or from less efficient purge of new insertions) but also gene conversion events among old copies. We tested these alternative hypotheses by considering that conversion events among old copies would generate multiple similar copies with an old age while recent bursts would generate multiple similar copies with a similar age of insertion as their time of divergence. The age of a copy can be estimated from the divergence between 5’-LTR and 3’-LTR sequences as they are identical at the time of insertion and then diverge gradually with time. Similarly, the 5’-LTR sequences of a parental copy and its progeny copy are identical at the time of transposition and then diverge gradually (Figure 5A). Therefore, copies resulting from a burst of transposition events should have accumulated roughly the same number of substitutions between their 5’- and 3’-LTR sequences as between their 5’-LTR sequence and the 5’-LTR sequence of their parental copy, *i.e.,* their most similar TE copy. We therefore expect that, in case of transposition bursts without conversion events, plotting the number of substitutions between the 5’-LTR sequences of the most similar copies against the number of substitutions between the 5’-LTR - 3’-LTR sequences within copies would result in a cloud of points around the first bisector, with some stochasticity. We estimated the variance around the first bisector expected without gene conversion by simulating a cloud of points under a Poisson distribution. The points outside the expected cloud, at the bottom right, likely represent conversion events (low number of substitutions between 5’-LTRs from different copies but elevated number of substitutions between LTRs within copies).

**Figure 4:**
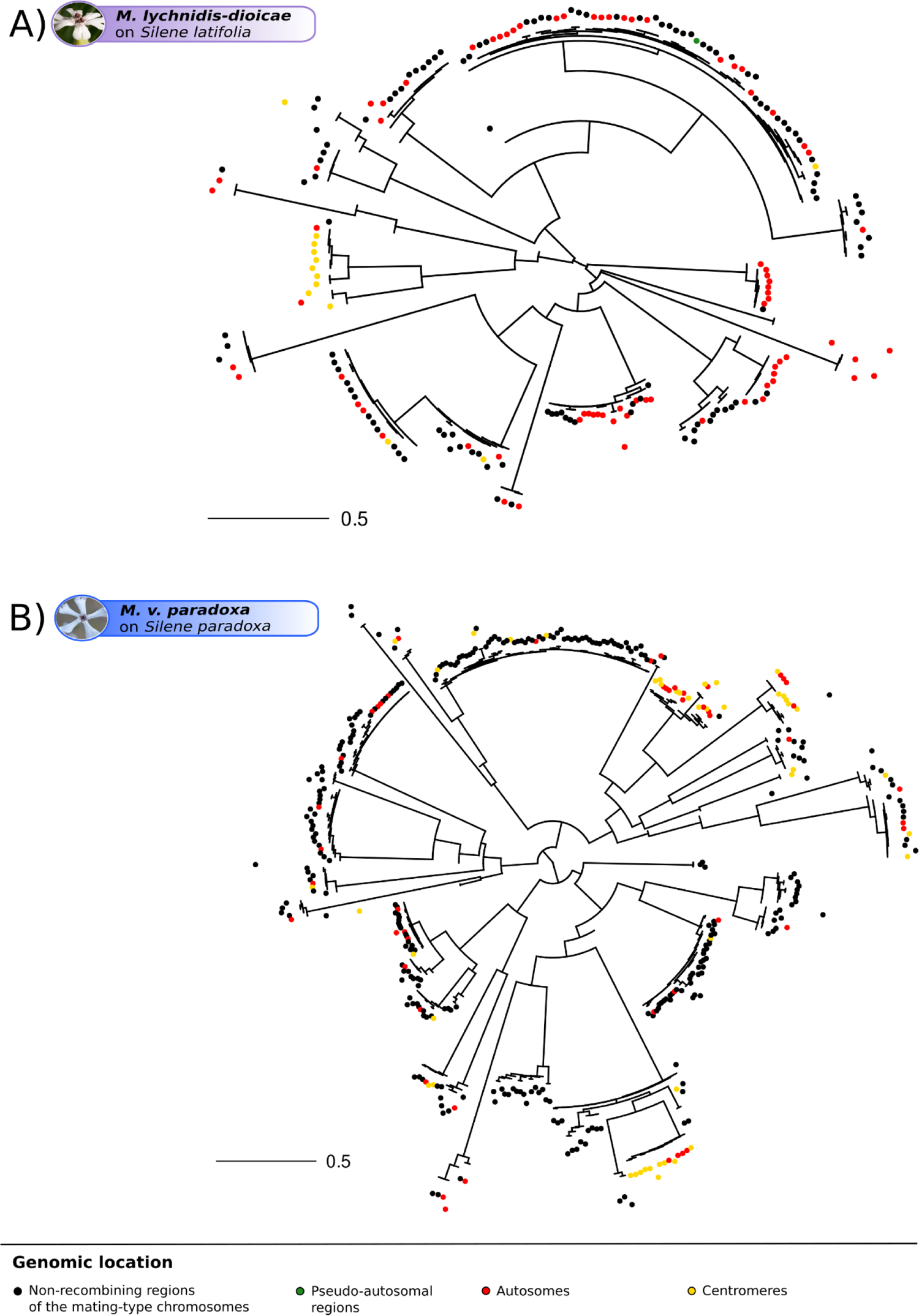
Single-species genealogies of *Copia* retroelement copies in *Microbotryum* genomes. Single-species genealogies of *Copia* copies in *Microbotryum lychnidis-dioicae* **(A)** and *Microbotryum violaceum paradoxa* **(B)**. The color of the dots at the tip of the branches corresponds to the genomic location of the transposable element (TE) copies. Other single-species *Copia* and *Ty3* genealogies are shown in Supplementary Figures S10 and S11, respectively.

**Figure 5:**
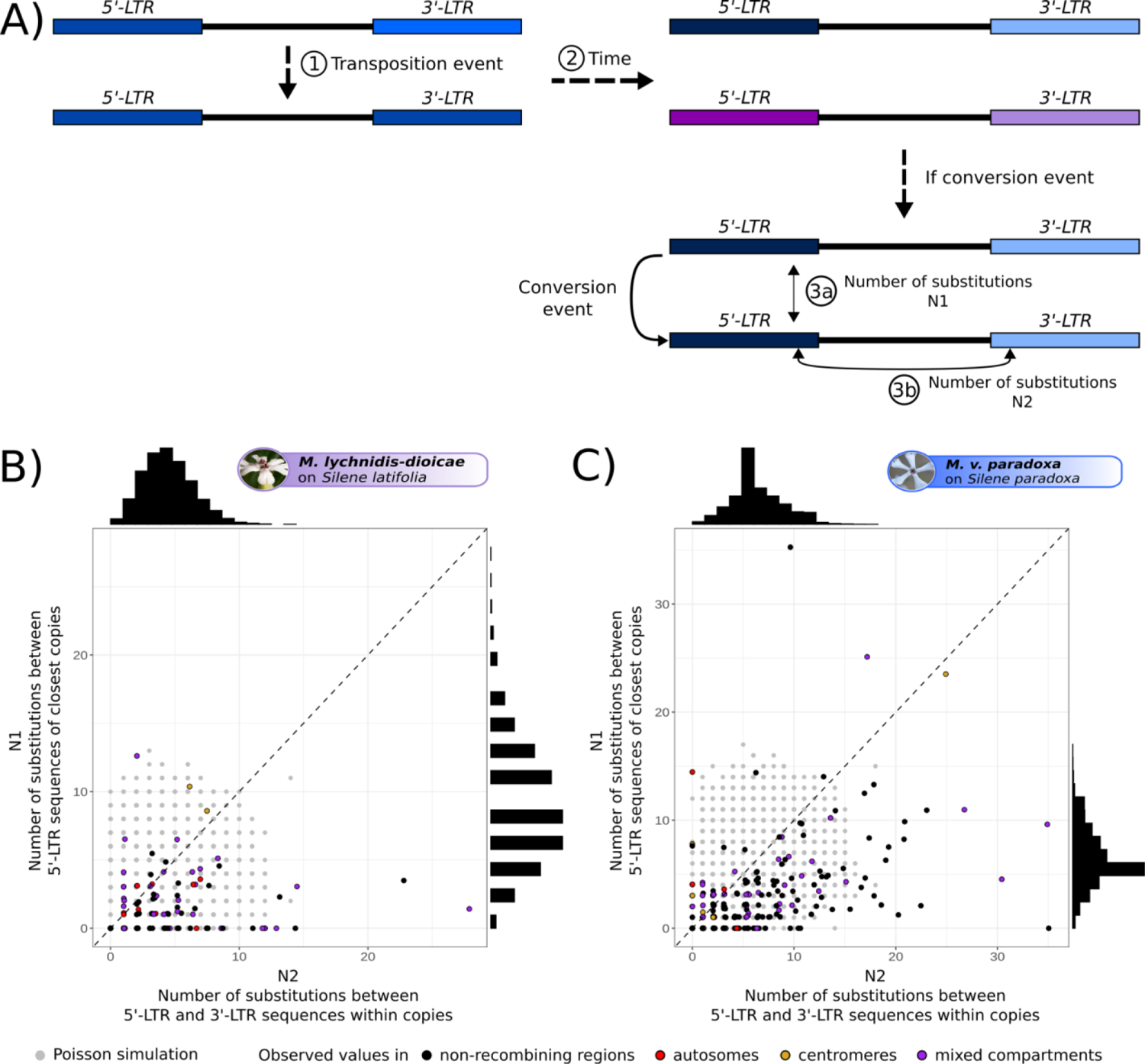
Investigation of the possible occurrence of conversion events within clusters of highly similar LTR-retrotransposon copies. **(A)** Illustration of LTR (long-terminal repeat) sequence divergence processes for retrotransposons: at the time of a new LTR-retrotransposon insertion, the 5’-LTR and 3’-LTR sequences are identical within the progeny copy and identical to the 5’-LTR of the parental copy (1). The LTR sequences then gradually diverge because they accumulate distinct mutations (2). The divergence between the 5’-LTR sequences of the two copies (N1) should thus be equal to the divergence between the 5’- and 3’-LTR sequences of the progeny copy (N2), the two measures being proportional to the age of the new copy. Following a conversion event, the 5’-LTR sequences between copies will be identical (3a) while the divergence between the 5’- and 3’-LTR sequences within copies remain high (3b). **(B)** and **(C)** Comparison of the observed number of different substitutions, between LTR sequences within (N2) and between copies (N1), in color, to the expected distribution under a Poisson model, in gray, with a lambda parameter equal to the average number of substitutions between 5’- and 3’-LTR sequences within copies, for the *Copia* retroelements in *Microbotryum lychnidis-dioicae* (B) and *M. violaceum paradoxa* (C). If no gene conversion occurred, we expect a cloud of points around the first bisector (y=x, dashed line) when plotting the number of substitutions between 5’-LTR sequences of the pairs of most similar copies within putative bursts against their 5’ - 3’-LTR sequence number of substitutions. In case of gene conversion, we expect points at the bottom right (low divergence between copies but old transposition ages). The points on the top left likely correspond to copies for which the closest related copy could not be identified. The colors of the points match the genomic location of the copy pairs. The observed distributions of the number of substitutions between 5’-LTR sequences of most similar copies and between 5’ - 3’ LTR sequences within copies are shown at the top and at the right, respectively.

In all species, we found that only a minority of points fell at the bottom right outside of the Poisson-simulated cloud of points, for the clusters of similar *Copia* or *Ty3* copies (Figures 5B and 5C, Supplementary Figures S11). This suggests that conversion events are rare and indicates that the clusters of similar copies correspond mostly to bursts of transposition, due to increased activity or less efficient selection against insertions, rather than gene conversion among old copies. For several species, a few points fell on the top left corner outside of the Poisson-simulated cloud of points, likely corresponding to copies for which the closest related copy could not be identified. The colors indicating the compartment of origin of the pairs of copies are scattered on the plots, showing that conversion is not more frequent in any genomic compartment (Figures 5B and 5C, Supplementary Figures S11).

The recent bursts of proliferation show that *Copia* and *Ty3* retrotransposons are active in *Microbotryum* genomes. In several species, the centromeric TE copies tended to cluster together (Figure 4 and Supplementary Figure S9 and S10). In a few species, such as *M. v. paradoxa* (Figure 4B), we identified *Copia* TE bursts specific to non-recombining regions. However, the clustering of copies from autosomes and non-recombining regions in the same TE clade with low divergence indicates that TEs transpose from one compartment to the other. We found similar distributions of pairwise divergence times between TE copies in autosomes and in the non-recombining regions of the mating-type chromosomes (Supplementary Figure S12). This indicates that bursts occurred at the same time in the two genomic compartments, thus suggesting that non-recombining regions serve as reservoirs from which TEs can transpose in autosomes. Further supporting the reservoir hypothesis, we found fewer pairs of most closely related copies with both copies in the non-recombining regions than expected by chance given the number of copies located in the non-recombining regions in the genealogies (decreasing by 67% compared to random distribution on average, ranging from 95.62% to 40.6% across species, Supplementary Figure S13).

In order to assess whether these TE bursts occurred following recombination suppression, we inferred TE genealogies for *Copia* and *Ty3* retroelements with the sequences of all species. The TE copies resulting from the recent TE bursts often clustered by species and by group of species deriving from the same recombination suppression event, supporting the inference that TE accumulation has followed recombination suppression (Figures 6 and Supplementary Figure S14). The bursts of retrotransposon accumulation occurred at different times across lineages, as shown by the different phylogenetic distances of the clades with short branches to the root (Supplementary Figure S15A and S15B). The only exception was the occurrence of three *Copia* bursts at the same time, in the three species of the *M. v. gracilicaulis* clade, as the three bursts appeared equidistant to the root (Supplementary Figure S15A, pointed by arrows and aligned on the green circle).

**Figure 6:**
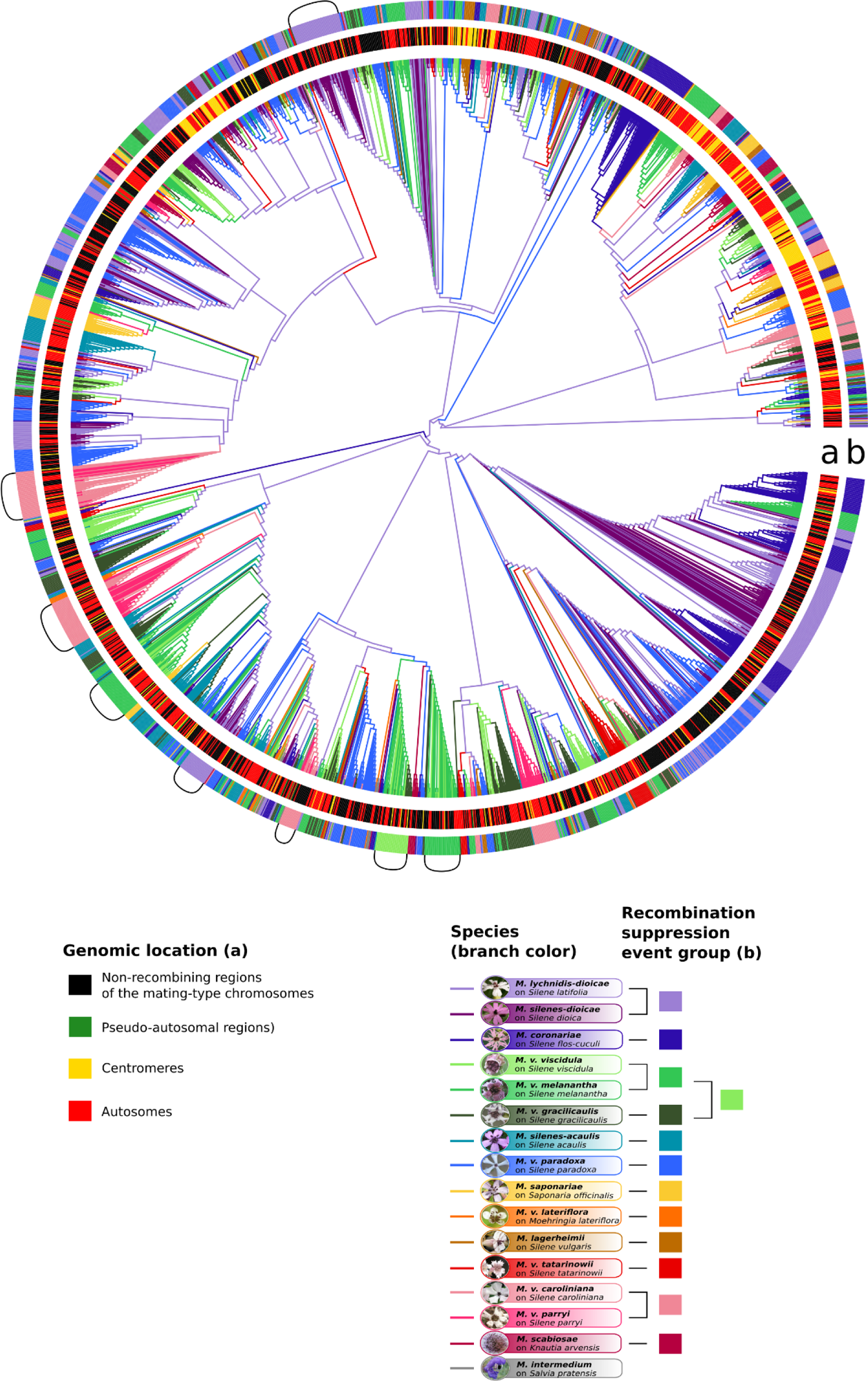
Multi-species genealogies of *Copia* retroelement copies in *Microbotryum* genomes. Multi-species *Copia* tree of all *Microbotryum* species of this study. The branch color corresponds to the species, the first inner track (a) corresponds to the genomic location of the TE copies, the second outer track (b) corresponds to the linkage event group of the species carrying the TE copy. Brackets highlight bursts of TEs. Multi-species *Ty3* tree of all *Microbotryum* species is shown in Supplementary Figure S14. Multi-species *Copia* and *Ty3* genealogies with branch lengths are shown in Supplementary Figure S1.

### Temporal dynamics of TE accumulation in non-recombining regions on mating-type chromosomes

The multiple independent events of recombination suppression across the *Microbotryum* phylogeny, with a range of ages, provides a unique opportunity to investigate the temporal dynamics of TE accumulation. The large non-recombining regions of different ages formed by the stepwise recombination suppression between the HD and PR loci were highly rearranged in *M. v. viscidula*, *M. v. melanantha*, *M. v. gracilicaulis* and *M. v paradoxa,* rendering difficult the assignment of some TE copies to a given age of recombination suppression. In order to be conservative, we therefore only considered TE copies that could be assigned to a stratum with high confidence, by taking into account only blocks of at least 80 kb (size of the pink stratum, the smallest stratum of the dataset) and encompassing at least two genes of a given stratum. The resulting blocks represented 66.52 to 90.52% of the total non-recombining region sizes (Supplementary Figure S16), thus likely yielding good estimates of their TE content. We excluded from analyses the orange stratum (Figure 1) shared by all or most of the species (Branco *et al*., 2017, 2018; Carpentier *et al*., 2019, 2022; Duhamel *et al*., 2022), as it is small and might not represent independent events of recombination suppression. We also excluded from analyses the small strata that were too much fragmented (for example the red stratum in *M. lychnidis-dioicae* and *M. silenes-dioicae;* Figure 1; Branco *et al*., 2018). To avoid pseudoreplication, we considered the mean of the percentages of base pairs occupied by TEs in the a_1_ and a_2_ genomes for each species. Similarly, for the strata shared by multiple species, we took the mean values across the species derived from the same recombination suppression event. We used the TE content in the autosomes of *M. intermedium* as the point at time zero, *i.e.,* before recombination suppression, as its mating-type chromosomes still recombine over almost their entire length, thereby precluding any potential TE reservoir effect.

We tested whether the percentage of base pairs occupied by TEs in the non-recombining regions could be explained by the time since recombination suppression and the ancestral size of the non-recombining region. We compared generalized additive models assuming linear or logarithmic relationships with the time since recombination suppression, with or without smoothing splines (*i.e.*, phases), as well as a non-linear negative exponential model. The percentage of base pairs occupied by TEs in the non-recombining regions was best explained by the negative exponential model with the age of the stratum as an explanatory variable of the percentage of TEs in this stratum (Figure 7 and Supplementary Table 6). We found that TEs rapidly accumulated in the first 1.5 million years following recombination suppression and then reached relatively abruptly a plateau at around 50% occupancy of TEs. A similar curve was obtained when fitting TE accumulation with the average synonymous divergence (d_S_) between a_1_ and a_2_ associated alleles of the genes within strata, as expected considering the quasi-linear relationship between d_S_ and stratum age (Supplementary Figure S17). The ancestral size of the non-recombining region did not significantly impact TE content in the retained negative exponential model.

**Figure 7:**
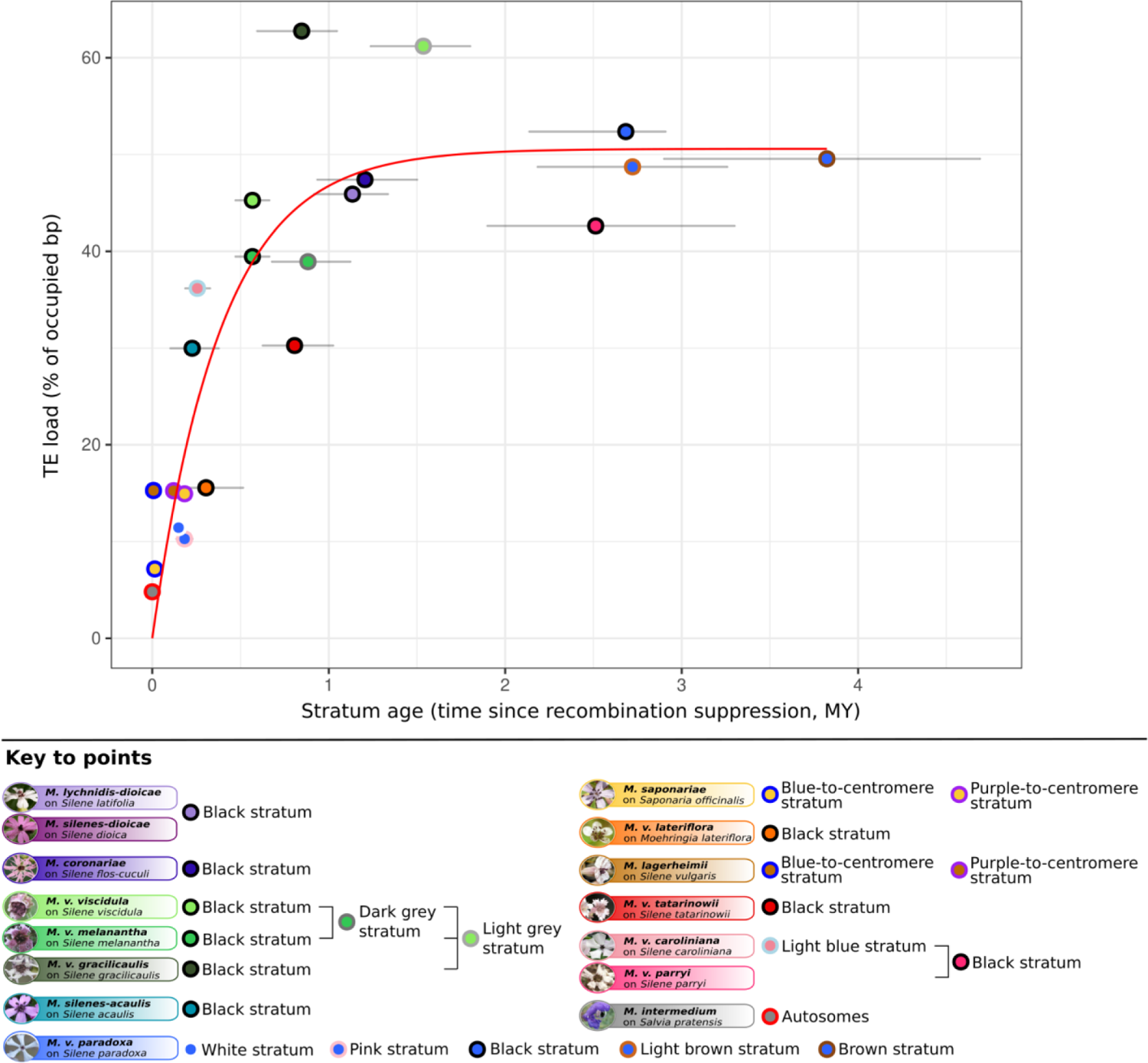
Tempo of transposable element (TE) accumulation in the non-recombining regions of the *Microbotryum* mating-type chromosomes. Percentage of base pairs occupied by TEs in non-recombining regions (means in a_1_ and a_2_ genomes and means across species sharing the same strata) as a function of their age in million years (MY) with confidence intervals. The red curve shows the prediction of the best model corresponding to a negative exponential model. Each dot corresponds to an independent evolutionary stratum.

We found that the relative abundance of RIP-like footprints (as assessed by the normalized RIP-index metric, see Methods) was highly variable in the young strata and remained constant with time since recombination suppression, with normalized RIP-index values greater than zero, suggesting that a process similar to RIP efficiently mutates TE copies in young non-recombining regions (Supplementary Figure S18). There was indeed no significant correlation between the age since recombination suppression and the abundance of RIP-like footprints in the TEs of the strata (two-sided Pearson’s correlation test, *p-value* = 0.9). The fraction of detected intact *Helitron* copies decreased with time since recombination suppression and even reached zero after 4 MY (Supplementary Figure S19). The proportion of intact copies of *Copia* and *Ty3* retroelements decreased rapidly following recombination suppression, from 20% down to 10%, and then remained stable at around 10% (Supplementary Figure S19).

## Discussion

We found a large degree of variation among *Microbotryum* species in their TE load, and the key to understanding this variation seems to be the evolution of the recombination landscape associated with regions determining reproductive compatibility. Moreover, there appears to be a significant feedback of the TE accumulation on non-recombining mating-type chromosomes where the buildup is rapid, and the rest of the genome is then burdened with greater repetitive element insertions. Leveraging on a unique dataset with 21 regions having independently evolved recombination suppression, we found that TEs accumulated rapidly following recombination cessation but then reached a plateau at ca. 50% of occupied base pairs, after 1.5 MY.

We indeed revealed more details on the degree of variation in genomic TE loads among *Microbotryum* genomes than previously recognized, ranging from 6.5 to 30% base pairs occupied by TEs, with an intermediate TE content in the most studied species, *M. lychnidis-dioicae* (19.4%). We were also able to annotate a large proportion of the detected TE copies, *e.g.* 89% in *M. lychnidis-dioicae*. In comparison, a previous study using Illumina paired-end sequencing estimated the genomic TE content of *M. lychnidis-dioicae* at 14.1% and only 60% of the TE copies were annotated (Amselem *et al*., 2015). Relative to other fungi, *Microbotryum* genomes carry intermediate TE contents, generally lower than their close relatives in the Pucciniomycotina (Raffaele & Kamoun, 2012). We found that LTR retrotransposons, especially from the *Copia* and *Ty3* superfamilies, represented the majority of the base pairs occupied by TEs in *Microbotryum* genomes. These LTR retrotransposons summed to 87% of the base pairs occupied by TEs in *M. intermedium* and 70% in *M. lychnidis-dioicae*, which was much higher than previously estimated and among the highest LTR retrotransposon content in fungi, relative to other TE categories (Amselem *et al*., 2015).

Our results also indicate that the lack of recombination has allowed the accumulation of TEs by a lower efficacy of selection against deleterious insertions, as shown by the higher TE loads in non-recombining regions of mating-type chromosomes compared to recombining autosomal or pseudo-autosomal regions. The accumulation of TEs has been widespread in non-recombining regions of sex chromosomes of plants and animals (Erlandsson *et al*., 2000; Bachtrog, 2003, 2005; Marais *et al*., 2008; Ahmed *et al*., 2014; Śliwińska *et al*., 2016; Mawaribuchi *et al*., 2017). In the absence of recombination, the average number of TE copies can only increase with time in the population, because chromosomes devoid of new TE copies cannot be generated by recombination and the chromosomes carrying fewer TE copies can be randomly lost, a phenomenon known as Muller’s ratchet (Muller, 1918; Charlesworth & Charlesworth, 2000; Bachtrog, 2003; Dolgin & Charlesworth, 2008). In *Drosophila miranda*, TEs played an early and important role in the degeneration of the neo-Y chromosome (Bachtrog, 2005; Bachtrog *et al*., 2008). We detected RIP-like footprints in *Microbotryum* genomes, but this genome defense mechanism was not sufficient to control TE proliferation in the absence of recombination, as previously noted in some *Microbotryum* species (Johnson *et al*., 2010; Horns *et al*., 2017).

Transposable elements from the *Copia*, *Ty3* and *Helitron* superfamilies were particularly involved in the repeat accumulation in non-recombining regions. *Copia* and *Ty3* retrotransposons proliferated independently and repeatedly following the multiple independent events of recombination suppression across *Microbotryum* lineages. *Copia* and *Ty3* superfamilies are predominant in *M. intermedium* and in autosomes of the species with non-recombining mating-type chromosomes, suggesting that they were already abundant before recombination suppression, and then proliferated due to transposition and Muller’s ratchet. *Helitron* elements were in contrast rare in *M. intermedium* and in autosomes of all species, and yet have also accumulated in the multiple non-recombining regions, as observed in the non-recombining regions of the sex chromosomes of *Drosophila pseudoobscura* and *D. affinis* (Nguyen *et al*., 2022). *Helitrons* transpose by a mechanism similar to rolling-circle replication via a single-stranded DNA intermediate and have been suggested to preferentially insert in AT-rich regions; RIP and RIP-related mechanisms increase the percentage in AT and are more abundant in non-recombining regions, which could have facilitated the accumulation of *Helitrons* (Grabundzija *et al*., 2016). It is striking that the same superfamilies preferentially expanded repeatedly in multiple non-recombining regions of *Microbotryum* fungi. Their specific mechanisms of transpositions may be particularly efficient in regions where recombination is suppressed, perhaps due to the chromatin landscape (Quadrana *et al*., 2019). LTR retroelements for example preferentially target heterochromatin (Marsano & Dimitri, 2022).

Through the inferred TE copy genealogies of the most abundant superfamilies in non-recombining regions (*Copia* and *Ty3* superfamilies), we found recent accumulation of TE copies through bursts occurring in all genomic compartments, *i.e.*, the non-recombining regions, the autosomes and the centromeres, many of them being shared by species deriving from the same recombination suppression event. This insight into the dynamics of TE accumulation in the different genomic compartments suggests that TE accumulation in *Microbotryum* genomes followed recombination suppression, not only in the non-recombining regions, but also in the rest of the genome. TE accumulation in non-recombining regions therefore have genome-wide impacts. Our TE genealogies were based on a fraction of the *Copia* and *Ty3* retrotransposon copies, due to the necessity to identify the 5’-LTR sequences. However, the 5’-LTR sequences were sometimes lost in degraded TE copies or could not be distinguished from the 3’-LTR sequence when the coding sequences were degenerated and could not be oriented. Nevertheless, considering the very low divergence between the TE copies within the genealogies, the burst patterns should not be affected by the sub-sampling of TE copies. Moreover, we found independent TE bursts across multiple *Microbotryum* species, indicating that the sampling of the TE copies used for the genealogies was good enough to capture accumulation events older than species divergence. Furthermore, the comparison of LTR sequence divergence within and between copies indicated that the clusters of similar copies were due mostly due to recent bursts of transposition (due to increased TE activity or less efficient selection against their insertion), with only rare conversion events among copies. This analysis thus informs on TE dynamics in genomes of species with a large range of ages of recombination suppression. Supporting the TE reservoir hypothesis (Hood *et al*., 2004; Wei *et al*., 2020), we found a similar pattern of copy near identity between the non-recombining regions and the autosomes of the *Copia* and *Ty3* retroelements and a strong significant positive correlation between TE load in the non-recombining regions and the autosomes. The TE bursts likely result from relaxed purifying selection following recombination suppression rather than from increased intrinsic TE activity, as shown also following polyploidization (Baduel *et al*., 2019).

We found similar TE contents in pseudo-autosomal regions compared to autosomes, without preferential accumulation of TE copies at the margin of the non-recombining regions. TEs have nevertheless accumulated everywhere in genomes following the formation of a non-recombining region acting as a TE reservoir. TE copies thereby accumulating at the margin of the non-recombining regions may impact their evolution (Maloisel & Rossignol, 1998; Ben-Aroya *et al*., 2004; Yelina *et al*., 2012; Kent *et al*., 2017). TEs can trigger chromosomal rearrangements (Balachandran *et al*., 2022), and they recruit silencing marks, such as DNA methylation and heterochromatin formation, that can suppress recombination (Maloisel & Rossignol, 1998; Ben-Aroya *et al*., 2004; Yelina *et al*., 2012, 2015; Bachtrog, 2013; Li *et al*., 2016; Kent *et al*., 2017). TEs can also induce deleterious mutations, which can select for recombination suppression that ensures the sheltering of the genetic load they create (Jay *et al*., 2022). A DNA fragment evolving recombination cessation that captures fewer deleterious mutations than the population average should be selected for, but this region will have its genetic load exposed to selection in the homozygous state more often when it rises in frequency unless it also captures a permanently heterozygous allele such as the haploid self-incompatibility of a mating-type allele (Jay *et al*., 2022). The more often that deleterious mutations segregate in genomes then the more this effect can act: a given fragment evolving recombination cessation and capturing a permanently heterozygous allele will have a greater chance of having a higher fitness differential with the average locus if the distribution of the number of deleterious mutations in this fragment is wider in the population, resulting in stronger selection (Jay *et al*., 2022). We found that TEs did not accumulate preferentially in the margin of the non-recombining regions, in contrast to our prediction based on a recent theoretical model showing that, in automictic species such as *Microbotryum* fungi, a mating-type locus can shelter deleterious mutations in its flanking regions (Tezenas *et al*., 2022). As TE insertions can be deleterious, one may have expected that selfing would have promoted their purge more easily farther away from the mating-type locus (Tezenas *et al*., 2022). Not all TE insertions are however deleterious and their active spread may counteract the purging effect.

We found that TEs have rapidly accumulated in mating-type chromosomes following recombination suppression, but TE content reached a plateau relatively abruptly at about 50% of occupied base pairs after 1.5 MY. The rapid initial accumulation of TEs is expected: under the lack of recombination, new TE insertions cannot be purged, and each new copy can further make new copies (Quadrana *et al*., 2019; Legrand *et al*., 2019). The deceleration of TE accumulation after 1.5 MY can be due to several, non-exclusive phenomena: i) selection against new insertions or for a more efficient control may be stronger when the load becomes higher, particularly if its cost increases with copy number, or if TE insertions are more deleterious with time because genes become hemizygous. Indeed, TE insertions in genes would be sheltered in a heterozygous state early after recombination suppression while there are two copies of each gene in the diploid (or dikaryotic) state, but importantly not following the loss of function for one copy or the gene becoming hemizygous, as often occurs in sex chromosomes and in particular in *Microbotryum* fungi (Bachtrog, 2013; Badouin *et al*., 2015); ii) extant control mechanisms against TE proliferation can become more efficient with time by a cumulative effect, e.g. if methylation accumulates with generations (Teixeira & Colot, 2010), or could even evolve to become more efficient; iii) new TE insertions nested in more ancient copies can further contribute to inactivate TEs, as suggested by the decreased proportion of *Helitrons* intact copies with time since recombination suppression in non-recombining regions; *Helitrons* in particular often display high proportions of non-autonomous copies (Sweredoski *et al*., 2008); iv) the chromatin landscape of non-recombining regions may become less favorable to new insertions, as for example histone can induce a preferential integration of C*opia* retrotransposons (Quadrana *et al*., 2019).

The rate of degeneration for non-recombining regions in *Microbotryum* fungi has been previously analyzed in terms of non-synonymous substitutions and the frequency of optimal codons (Carpentier *et al*., 2022). Purifying selection on non-synonymous substitutions was lower in non-recombining regions than in autosomes and remained nearly constant over time (Carpentier *et al*., 2022). However, the substitutions toward non-optimal codons increased rapidly following recombination suppression and then decelerated to reach a level below random codon usage. It is not unexpected to find different temporal dynamics of degeneration for DNA substitutions and TE insertions, as they correspond to very different mechanisms of evolution.

## Conclusion

In conclusion, this study leverages on a unique dataset of 21 independent evolutionary strata of different ages in closely related species to assess the dynamics of TE accumulation with time following recombination suppression. We show that TEs have rapidly accumulated following recombination cessation and that the TE accumulation slowed down abruptly after 1.5 MY. We further show that some superfamilies repeatedly expanded in independent non-recombining regions, and in particular *Helitrons* that were not yet abundant before recombination suppression. *Copia* and *Ty3* superfamilies were also over-represented in non-recombining regions and have accumulated through bursts of proliferation, both in the non-recombining regions of the mating-type chromosomes and in the autosomes of *Microbotryum* species at the same time; this finding supports the TE reservoir hypothesis, i.e., that the TEs accumulated in non-recombining regions have a genome-wide impact by transposing to recombining regions. This study thus sheds light on important processes to improve our knowledge on genome evolution and in particular on the consequences of recombination suppression.

## Material and methods

### Detection and annotation of transposable elements

Transposable elements (TEs) were detected *de novo* in the a_1_ and a_2_ haploid genomes of 16 *Microbotryum* species and the red yeast *Rhodothorula babjavae* using LTRharvest (Ellinghaus *et al*., 2008) from GenomeTools 1.5.10, performing long-terminal repeat (LTR) retrotransposons detection and RepeatModeler 1.0.11 (Smit & Hubley, 2008) combining results from three other programs: RECON (Bao & Eddy, 2002), RepeatScout (Price *et al*., 2005) and Tandem Repeats Finder (Benson, 1999). The TE detection was enriched by BLASTn 2.6.0+ (Altschul *et al*., 1990) using the genomes as a database and the previously detected TE models as queries. To fulfill the repeat criterion, a TE sequence detected by RepeatModeler or LTRHarvest and its BLAST hits were retained only if the query matched at least three sequences in the same species with an identity ≥ 0.8, a sequence length > 100 bp and a coverage ≥ 0.8 (defined as the query alignment length with removed gaps divided by the query length). When these criteria were met, we retained the other query matches for the following parameters: identity ≥ 0.8, sequence length > 100 bp, e-value ≤ 5.3e−33 (25th percentile of e-value distribution) and coverage ≥ 0.8.

We performed TE annotation (Wicker *et al*., 2007) using the fungal Repbase database 23.05 (Bao *et al*., 2015) based on sequence similarity. We first performed a BLASTn search and a BLASTx search using the fungal Repbase DNA and protein sequence database, respectively, with TE sequences as queries. For each of these similarity-based searches, we set the minimum e-value score at 1e−10 and minimum identity at 0.8. Then, we performed protein domain detection using pfam_scan.pl (ftp://ftp.ebi.ac.uk/pub/databases/Pfam/Tools/) on TE sequences and compared them to protein domain detection in the fungal Repbase database, keeping matches with e-values ≤ 1e−5. We also considered the annotation found by RepeatModeler. In order to determine the best annotation of a TE sequence, we applied a majority rule: we chose the most frequent annotation, and, in case of equality, we kept both annotations, generating a multiple annotation; these multiple annotations can be due to TEs being inserted within other TE copies, generating nested TEs. When we found no annotation, we assigned the TE sequence to an “unclassified” category; we discarded such annotations when overlapping with predicted genes to avoid false positives. We also discarded predicted TE sequences annotated as rRNA or overlapping *Microbotryum* ribosomal sequences (downloaded from https://www.arb-silva.de/ and http://combio.pl/rrna/taxId/5272/) and mitochondrial DNA sequences (NC_020353) with e-value ≤ 1e-10. We used a python script for annotation, and the TE detection and annotation pipeline is available at https://gitlab.com/marine.c.duhamel/microtep.

### Phylogenomic species tree reconstruction

After removing the genes overlapping with annotated TEs, we inferred a species phylogenomic tree from the coding sequences of the 3,669 single-copy genes present in all *Microbotryum* species and the outgroup *Rhodothorula babjavae*, as done previously (Duhamel *et al*., 2022).

### Evolutionary histories of *Copia* and *Ty3* superfamily retroelements

We inferred the evolutionary histories of the most abundant elements, *i.e.,* in the *Copia* and *Ty3* retroelement superfamilies, by reconstructing their genealogies based on the sequence of their long terminal repeat (LTR) region at the 5’ end of the elements (5’-LTR), the rationale being that, at the time of transposition, the LTR sequences of the new copy are identical to the 5’-LTR sequence of its progenitor (Zhang *et al*., 2014). We only retained sequences harboring one or two LTR sequences identified by LTRHarvest (Ellinghaus *et al*., 2008), thus avoiding nested copies. We distinguished the 5’-from the 3’-LTR based on the orientation of their coding sequences detected using RepeatProteinMask (Smit & Hubley, 2008). Because some retrotransposon sequences were degenerated, their coding sequences could not be identified and thus oriented, so that only a subset of the retrotransposons were used for building the genealogies. We built multi-species genealogies using the TE copies of *Copia* and *Ty3* retrotransposons in all genomes (2,468 and 694 copies, respectively). We built single-species (*i.e.*, within genome) genealogies, only for species with at least 100 copies remaining after filtering (see proportion of TE copies used for the genealogies in Supplementary Figure S8). We then aligned the 5’-LTR sequences of *Copia* or *Ty3* elements using MAFFT v7.310 (Katoh *et al*., 2002) (alignment length ranging from 816 to 2,767 bp, 1,678 bp on average). We used IQ-TREE 2.0.4 (Minh *et al*., 2020) to infer the maximum likelihood tree from the 5’-LTR sequences alignment under the GAMMA + GTR model of substitution. We assessed tree robustness with 1000 replicates for ultrafast bootstrap and SH-aLRT. Trees were plotted using phytools (Revell, 2012) and ggtree (Yu *et al*., 2017). Trees were displayed as rooted using midpoint rooting (*midpoint.root*, phytools, (Revell, 2012). Pairwise sequence divergence values in trees were obtained using *cophenetic.phylo* from ape 5.0 (Paradis & Schliep, 2019).

We wanted to assess whether the identified clusters of TE copies with low divergence to each other originated from bursts of transposition or conversion events among copies. In case of gene conversion, multiple copies would be similar while their age of insertion would be ancient, if an old copy had copied itself by gene conversion into several other copies. In contrast, in case of recent bursts, all copies should have a young age of insertion in addition to be similar to each other. Because the 5’-LTR and 3’-LTR sequences are identical at the time of insertion and then diverge (SanMiguel *et al*., 1998), we estimated the time since transposition of TE copies in the putative bursts by computing the number of substitutions between the 5’-LTR and 3’-LTR sequences, in the TE copies for which the 3’-LTR sequence could also be identified. For clusters of genetically similar copies, we compared the number of substitutions between 5’-LTR and 3’-LTR sequences per copy to the number of substitutions between the 5’-LTR sequences of pairs of most similar copies within the same cluster. For both measures, we computed the gamma-corrected Kimura 2-model parameter (**γ**-K2P), using MEGA11 (Tamura *et al*., 2021), and multiplied the divergence by the number of aligned sites. To compare the two values, we normalized the number of substitutions between the 5’- and 3’-LTR sequences by the number of aligned sites in the 5’-LTR sequences alignment. In case of bursts of transposition without conversion events, the 5’-LTR and 3’-LTR sequences of each copy diverge gradually and at the same rate as the 5’-LTR sequences between copies from the same burst (Figure 5A). Plotting the number of substitutions between 5’-LTR sequences of the most similar copies against the number of substitutions between the 5’-LTR - 3’-LTR sequences of each copy should therefore give a cloud of points around the first bisector. In case of conversion events, we expect that old copies (with divergent 5’-LTR and 3’-LTR sequences) have low divergence between one another while they have a high divergence between their 5’-LTR and 3’-LTR sequences, and therefore fall at the bottom right of the plot, far from the first bisector (Figure 5A). To assess whether clusters of highly similar copies were caused by conversion events within clusters, we therefore plotted the number of substitutions between 5’-LTR sequences between closest copies as a function of the number of substitutions between their 5’- and 3’-LTR sequences (normalized by the same alignment length), for TE copies within clusters of genetically similar elements. As the number of substitutions in a sequence are rare discrete events following a Poisson law (Felsenstein, 2003), we compared for each species this observed cloud of points to a cloud of points generated by drawing their two coordinates from a Poisson distribution, with a lambda parameter equal to the observed average number of substitutions between the 5’- and 3’-LTR sequences within copies, as this value is not affected by potential conversion events among copies, but only by divergence since insertion. The points outside the simulated cloud, in the right bottom sector (low number of substitutions between copies but high number of substitutions within copies) would result from conversion events.

### Search for potential horizontal transfers of transposable elements between *Microbotryum* species and their host plants

To search for potential horizontal transfers of transposable elements between *Microbotryum* fungi and their host plants, we compared the nucleotide sequences from the Repbase database 23.05 (Bao *et al*., 2015) of dicotyledones and *Arabidopsis thaliana* with the *Copia* and *Ty3* TE annotations from the *Microbotryum* genomes using BLASTN 2.6.0+ (Altschul *et al*., 1990).

### Detection of centromeres and telomeres in the autosomes of *Microbotryum* genomes

We identified centromeres (Supplementary Table 7) as in Duhamel *et al*. (2022) by blasting (BLASTN 2.6.0+, Altschul *et al*., 1990) the centromeric repeats previously described in *Microbotryum* fungi (Badouin *et al*., 2015) on autosomal contigs larger than 20 kb; the largest stretch of centromeric repeats on a given contig was annotated as its predicted centromere. We delimited the predicted centromere by recursively extending the focal region by 1 kb intervals as long as gene density remained lower than 0.25 in the focal window. We detected telomeres (Supplementary Table 8) within 100 bp regions at the end of contigs carrying at least five times the telomere-specific TTAGGG motif (CCCTAA on reverse complementary strand), as done previously (Badouin *et al*., 2015). Subtelomeric regions were defined as the 20 kb regions adjacent to the telomeres, plus the 100 bp telomeres.

### RIP-like index

We computed values for a RIP-like index in non-overlapping 1 kb genomic regions along the mating-type chromosomes and autosomal contigs as previously described (Beckerson *et al*., 2019). The RIP-like index was calculated as the ratio *t/n*, *t* being the ratio of the RIP-affected sites (TTG + CAA trinucleotides, forward and reverse; Hood *et al*., 2005) over the non RIP-affected sites (TCG + CGA, minus overlapping tetra-nucleotides TGCA) for the RIP target sites, and *n* being the same ratio for non-RIP target sites ([A,C,G]TG + CA[C,G,T] over [A,C,G]CG + CG[C,G,T] - [A,C,G]CG[C,G,T]), to control for sequence composition. The RIP index was normalized as *t/n - 1*, so that a normalized RIP index greater than 0 was indicative of an excess of RIP-like mutations compared to random expectations. The script used to compute the RIP index is available at https://gitlab.com/marine.c.duhamel/ripmic. We used the *SlidingWindow* function of the *evobiR* package (mean values, 10 kb steps - Blackmon & Adams, 2015) and *ggplot2* (Wickham, 2009) to plot the distribution of RIP along the mating-type chromosomes and autosomal contigs and identify the RIP-affected regions. We used the same parameters to plot the density of genes and TEs in the same non-overlapping 1 kb windows. In order to test the relationship between the abundance of RIP footprint in TEs in non-recombining regions and the age of evolutionary strata, we calculated the average of the RIP index of the TEs in each stratum and performed a two-sided Pearson’s correlation test.

### Estimates of TE content in non-recombining regions and their flanking regions

The distinct evolutionary strata within non-recombining regions have been previously distinguished based on the following criteria: i) the distribution of recombination suppression across the phylogenetic tree to assess when recombination suppression evolved and thereby identifying distinct evolutionary strata, *i.e.*, genomic regions that stopped recombining at different times, and ii) the level of trans-specific polymorphism (*i.e.*, alleles clustering by mating type rather than by species), which is also a strong indicator of the time of recombination suppression (Branco *et al*., 2017, 2018; Carpentier *et al*., 2019, 2022; Duhamel *et al*., 2022) (Supplementary Table 1). Young non-recombining regions, formed after the linkage of the PR and HD loci (*i.e.*, the white and pink strata in *M. v. paradoxa* and the light blue stratum in *M. v. caroliniana*; Figure 1) were not rearranged with other strata, so that it was straightforward to compute their percentages of base pairs occupied by TEs. In contrast, the ancient blue and purple strata (Figure 1), having evolved at the base of the *Microbotryum* clade, were too much rearranged and intermingled with other strata for their specific TEs to be identified in species with large non-recombining regions, but they were very small (*i.e.*, encompassing on average only 12 genes). We therefore pooled the ancient blue and purple strata with the larger strata with which they were intermingled. For the multiple independent evolutionary strata corresponding to the linkage of the HD locus with the PR locus in a single step (black strata; Figure 1), it was straightforward to compute their percentage of base pairs occupied by TEs. In *M. v. viscidula*, *M. v. melanantha*, *M. v. gracilicaulis* and *M. v paradoxa*, recombination suppression extended gradually to link the HD and PR loci and then underwent multiple rearrangements that intermingled their genes and TEs. In order to avoid misassignment of TEs to these intermingled strata, we only considered regions spanning at least 80 kb (corresponding to the size of the smallest stratum from the dataset, the pink stratum in *M. v. paradoxa*) and with only genes from a single stratum. We defined the limit between two strata as the middle point between two genes from two distinct rearranged strata. The resulting blocks assigned to strata overall represented 66.52 to 90.26% of the total non-recombining regions formed by intermingled strata (Supplementary Figure S15), which should thus give a good estimate of the stratum TE contents. We used circos (Krzywinski *et al*., 2009) to visualize the blocks assigned to strata. For the non-recombining regions linking the HD or PR loci and their respective centromere (blue-to-centromere and purple-to-centromere respectively, in *M. lagerheimii* and *M. saponariae*), the percentage of base pairs occupied by TEs was calculated in the region between the end of the centromere and the mating-type locus. For each stratum, we took the mean percentage of TEs across a_1_ and a_2_ haploid genomes, and for the strata shared across several species, the mean of the a_1_ and a_2_ values across all the species sharing the same recombination suppression event. The ancestral size of the non-recombining regions were estimated by calculating the average size of the corresponding genomic region in the a_1_ and a_2_ haploid genomes of *M. intermedium*, *i.e.,* the *Microbotryum* species most distant from the other species in our dataset, with unlinked mating-type loci and with little recombination suppression (Duhamel *et al*., 2022).

To test whether TEs also preferentially accumulate at the margin of the non-recombining regions, we compared the fraction of base pairs occupied by TEs within the 100 kb of the pseudo-autosomal regions flanking the non-recombining regions to the distribution of TE load in 100 kb non-overlapping windows in fully recombining autosomes (100 kb away from the centromeres and after removal of 20 kb corresponding to subtelomeric regions). We only considered the species with a length of the pseudo-autosomal region at least equal to 100 kb (thus excluding *M. v. paradoxa*) and with at least 30 non-overlapping windows of 100 kb on their autosomes (thus excluding *M. scabiosae* and *M. v. parryi*, for which the genome assemblies were too much fragmented). We set the window length at 100 kb because this is the DNA fragment size at which linkage disequilibrium decreases below r² = 0.2 in *Microbotryum* populations (Badouin *et al*., 2017). The distribution of the 100kb-window values on autosomes were plotted using the Sturges binning method.

We estimated the fraction of intact *Copia*, *Ty3* and *Helitron* copies in each stratum based on their length. For each stratum, we calculated the ratio of putative intact TE copies (5-7.5 kb for *Copia* and *Ty3* and 5-11 kb for *Helitron*) over the total number of *Copia* and *Ty3* copies or *Helitron* copies. For each stratum, we also calculated the mean fraction of intact TE copies across a_1_ and a_2_ haploid genomes and for the strata shared by several species, the mean across species deriving from the same recombination suppression event.

### Estimates of non-recombining region ages

In *Microbotryum* fungi, sexual reproduction occurs before the infection of a new plant, which will produce spores in its flowers at the next flowering seasons, so that one year corresponds to one generation for these fungi. We used the dates of recombination suppression estimated in previous studies (Duhamel *et al*., 2022; Carpentier *et al*., 2022), for 21 genomic regions corresponding to independent events of recombination suppression distributed across 15 *Microbotryum* species. The *M. scabiosae* non-recombining regions used in a previous study (Carpentier *et al*., 2022) was not considered here due to the large confidence interval on its estimated age. To this dataset of 15 *Microbotryum* species, we added the TE content in the autosomes of *M. intermedium* as the zero time point; indeed, the autosomes are recombining and there is little potential for a reservoir effect in *M. intermedium* as the non-recombining regions are very small, being restricted to close proximity around the separate mating-type loci (Duhamel *et al*., 2022); we set the age since recombination suppression at 0.00000003 MY for *M. intermedium* autosomes, corresponding to one day since the recombination suppression event, to avoid the zero value for logarithm computation in temporal dynamics analyses.

### Statistical analyses

We tested whether the genomic percentage of base pairs occupied by TEs was different between the a_1_ and a_2_ mating-type haploid genomes using a paired two-sided Wicoxon’s rank test. To compare the percentages of base pairs occupied by TEs in the autosomes, the pseudo-autosomal regions and the non-recombining regions of the mating-type chromosomes, we performed a variance analysis (ANOVA) after testing the normality of the TE percentage distributions (Kolmogorov-Smirnov test). We then performed post-hoc Tukey’s range tests to identify the pairs of genomic compartments which displayed significantly different TE loads, using the *TukeyHSD* R function providing adjusted p-values for multiple comparisons. We assessed the correlation between TE loads in autosomes and non-recombining regions using the two-sided Pearson’s test.

In order to assess the shape of the functions for accumulation of TEs in the non-recombining regions of the *Microbotryum* mating-type chromosomes, we performed regressions using generalized additive models (*gam* function of R, Wood, 2011) and non-linear regression models (*drm* function of the drc package, Ritz *et al*., 2019). Based on the Akaike information criterion (AIC), we evaluated the model fits for predicting the percentage of base pairs occupied by TEs in the non-recombining regions as a function of the time since recombination suppression, with the ancestral size of the non-recombining region as a covariable, and assuming either a linear or a logarithmic relationship with the time since recombination suppression, with or without smoothing spline, or a negative exponential model.

In order to assess the relationship between the fraction of intact *Copia*, *Ty3* and *Helitron* copies and time since recombination suppression, we performed a local regression using the *loess.as* function from the fANCOVA package, using the Akaike Information Criterion for automated parameter selection (Hurvich *et al*., 1998; Wang & Ji, 2020). We performed all R analyses in R version 4.2.1.

## Supplementary information

**Additional file 1.** Supplementary figures S1-S14.

**Additional file 2.** Supplementary figures S15-S19.

**Additional file 3.** Supplementary tables 1-8.

**Additional file 4.** Datasets published on Figshare.

## Declarations

**Ethics approval and consent to participate**: Not applicable.

**Consent for publication**: Not applicable.

## Availability of data and materials

The datasets supporting the conclusions of this article are included within the article (Supplementary Tables 1 - 8). All data generated or analysed during this study are included in this published article and the supplementary information files. The TE detection pipeline is available at https://gitlab.com/marine.c.duhamel/microtep. The script used to compute the RIP index is available at https://gitlab.com/marine.c.duhamel/ripmic.

## Competing interests

The authors declare that they have no competing interests.

## Funding

This work was supported by the National Institute of Health (NIH) grant R15GM119092 to M. E. H., the Louis D. Foundation award and the EvolSexChrom ERC advanced grant #832352 to T. G., and an IDEX Paris-Saclay Bochum University PhD grant to M.D.

## Author contributions

M.D., T.G., R.C.R.d.l.V. and M.E.H designed the study. T.G., R.C.R.d.l.V. and M.E.H supervised the study. M.D. performed the genomic and statistical analyses and produced the figures. R.C.R.d.l.V. advised in genomic analysis and data visualization. T.G. and M.D. wrote the original draft. All authors contributed to the manuscript.

## Acknowledgements

We thank Dominik Begerow, Sebastian Klenner, Lena Steins and Frederick Witfeld for providing useful comments on the manuscript. We thank Gilles Fisher, Paul Jay, Jacqui Shykoff and Olivier Gascuel for useful discussions.

---

**Supplementary figure S1:**
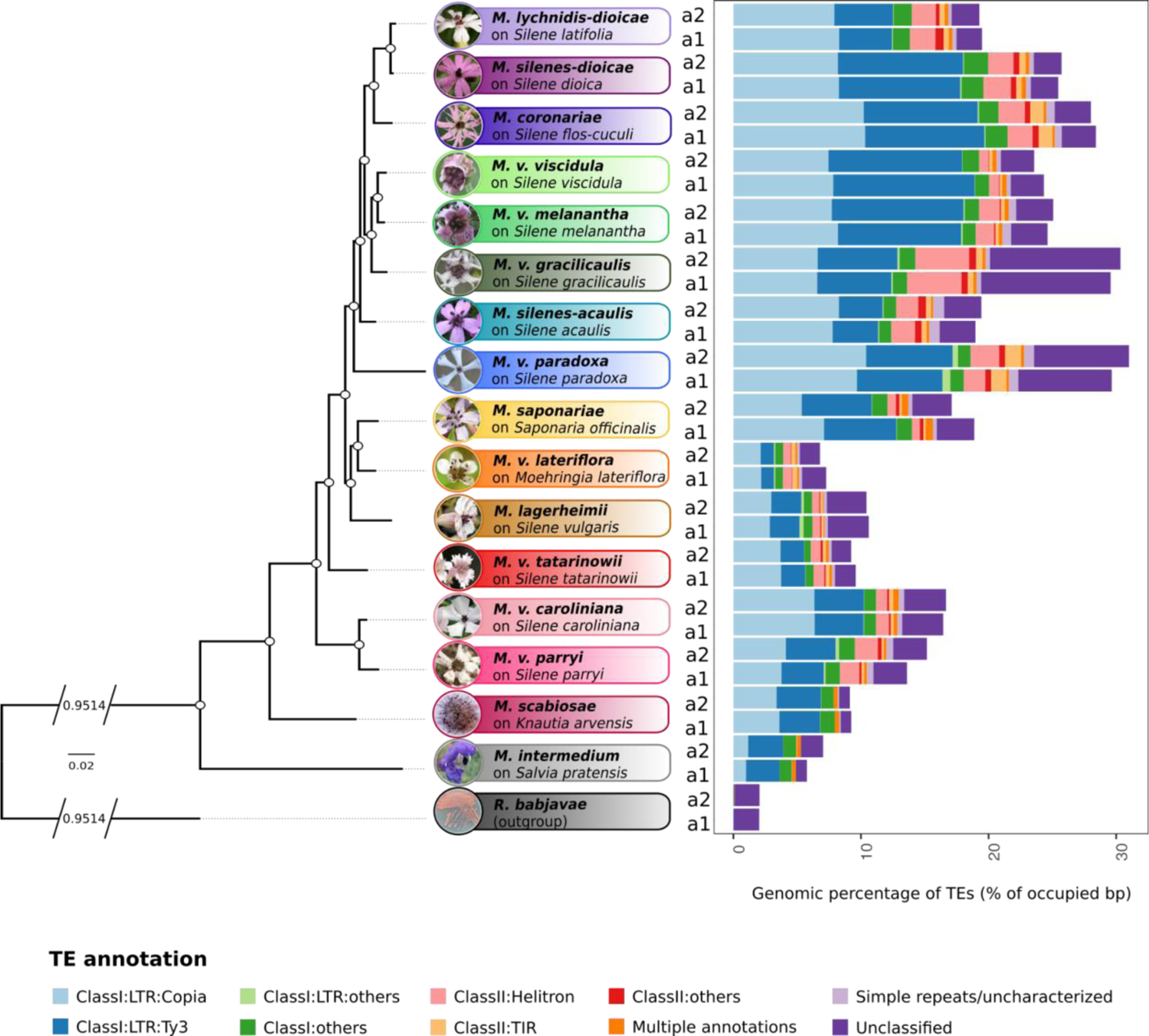
Percentage of base pairs occupied by transposable elements (TEs) in *Microbotryum* genomes, both for a_1_ and a_2_ mating-type chromosomes. Whole-genome TE content across the *Microbotryum* phylogeny, for each mating-type genome, each color representing a TE category.

**Supplementary figure S2:**
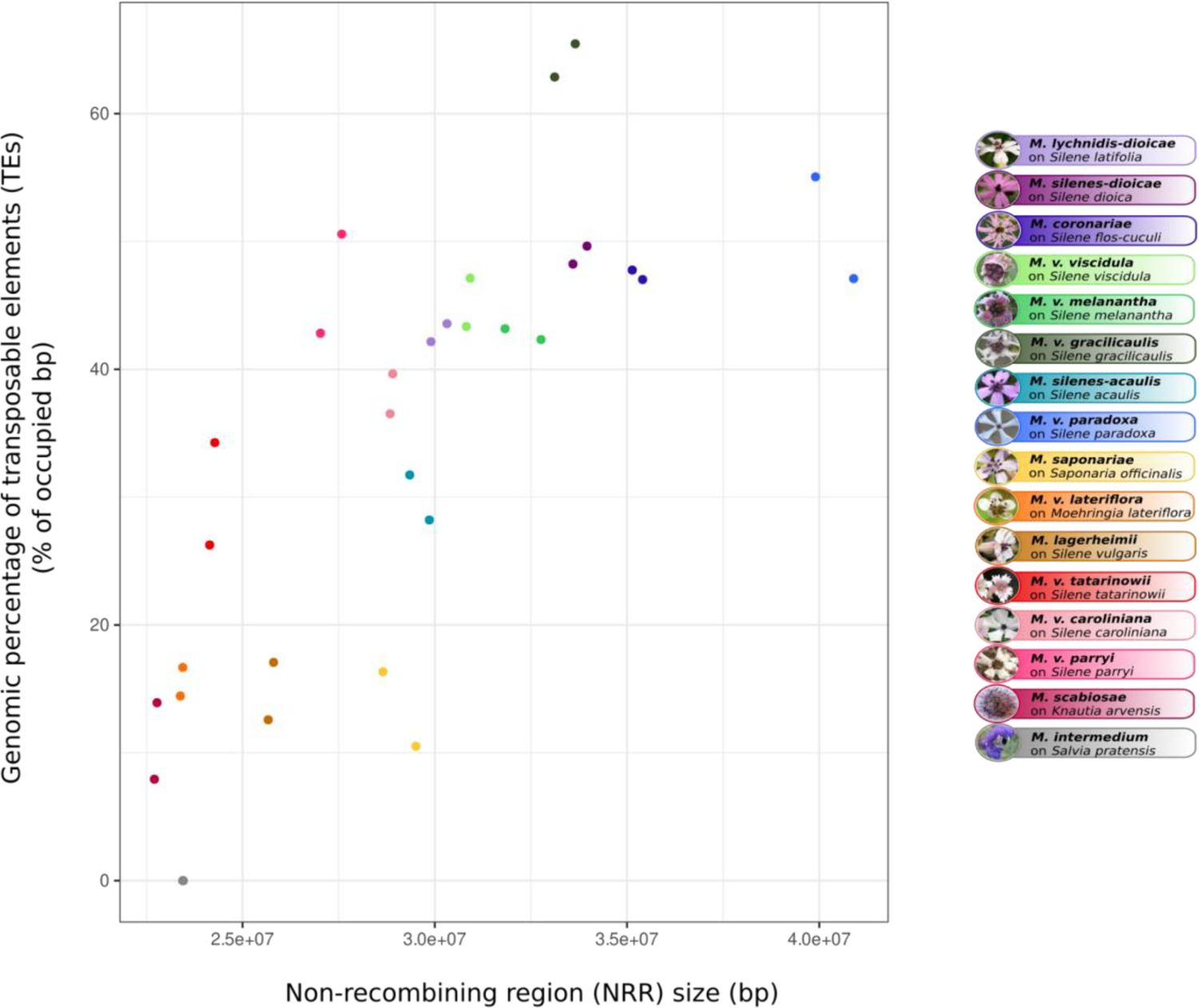
Relationship between the percentage of base pairs occupied by TEs and the total size of non-recombining regions. Each color represents one species, one point per mating type.

**Supplementary figure S3:**
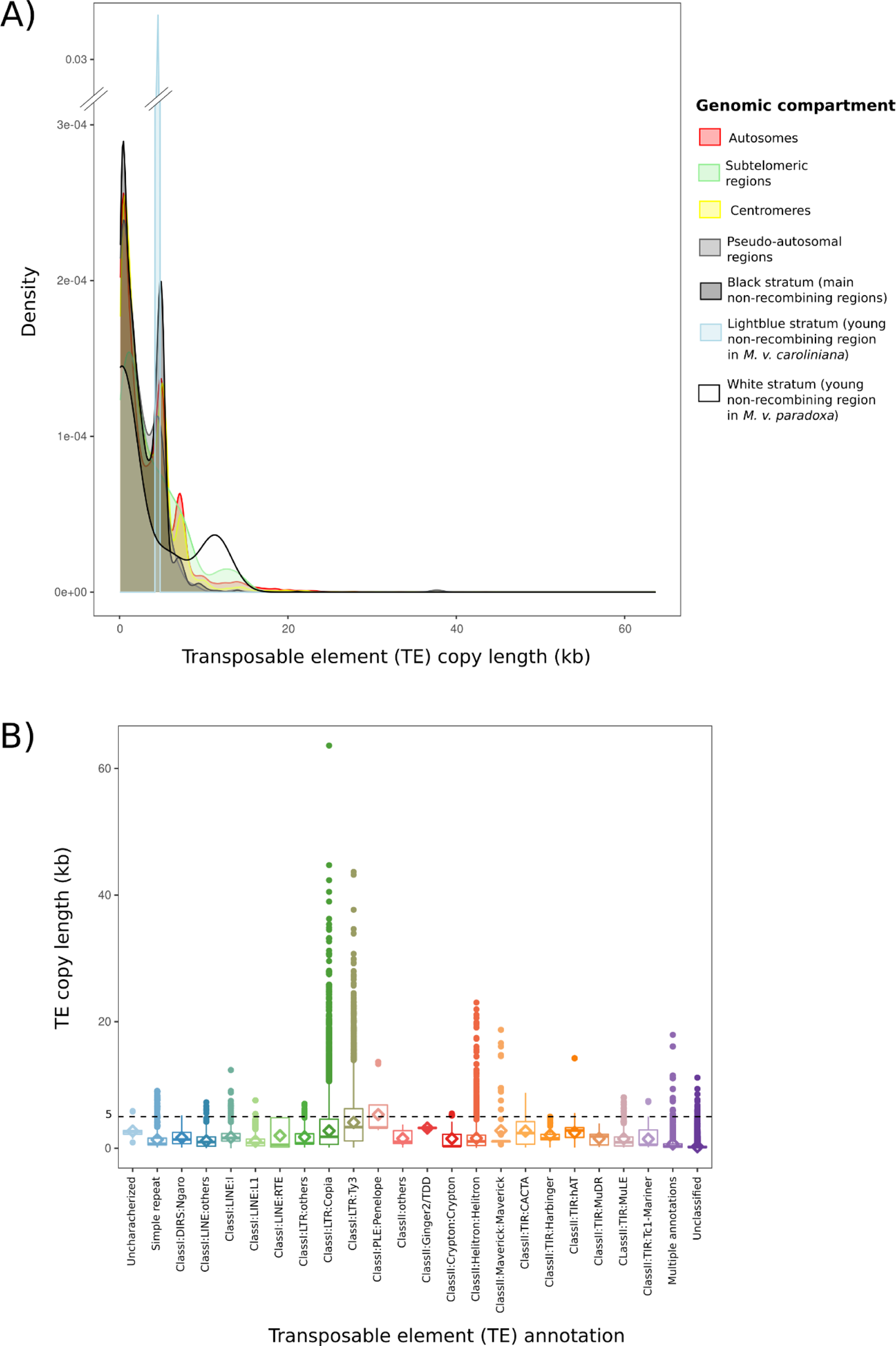
Distribution of transposable element (TE) copy length in *Microbotryum* genomes. **A)** Distribution of the *Copia* and *Ty3* retrotransposon copy length in the different genomic compartments in various *Microbotryum* species. Note that the Y axis is interrupted by parallel bars. **B)** Distribution of the length of individual TE copies for each TE annotation, across all *Microbotryum* genomes. Each color corresponds to a specific annotation. The dashed line indicates the 5 kb expected mean length of *Copia* and *Ty3* retroelements. Similar annotations representing a small proportion of the *Microbotryum* genomes were pooled in all the other figures; e.g., *ClassII:TIR* with two inverted repeats corresponds to all the *TIR* elements, i.e., *CACTA, Harbinger, hAT, MuDR, MuLE* and *Tc1-Mariner*.

**Supplementary Figure S4:**
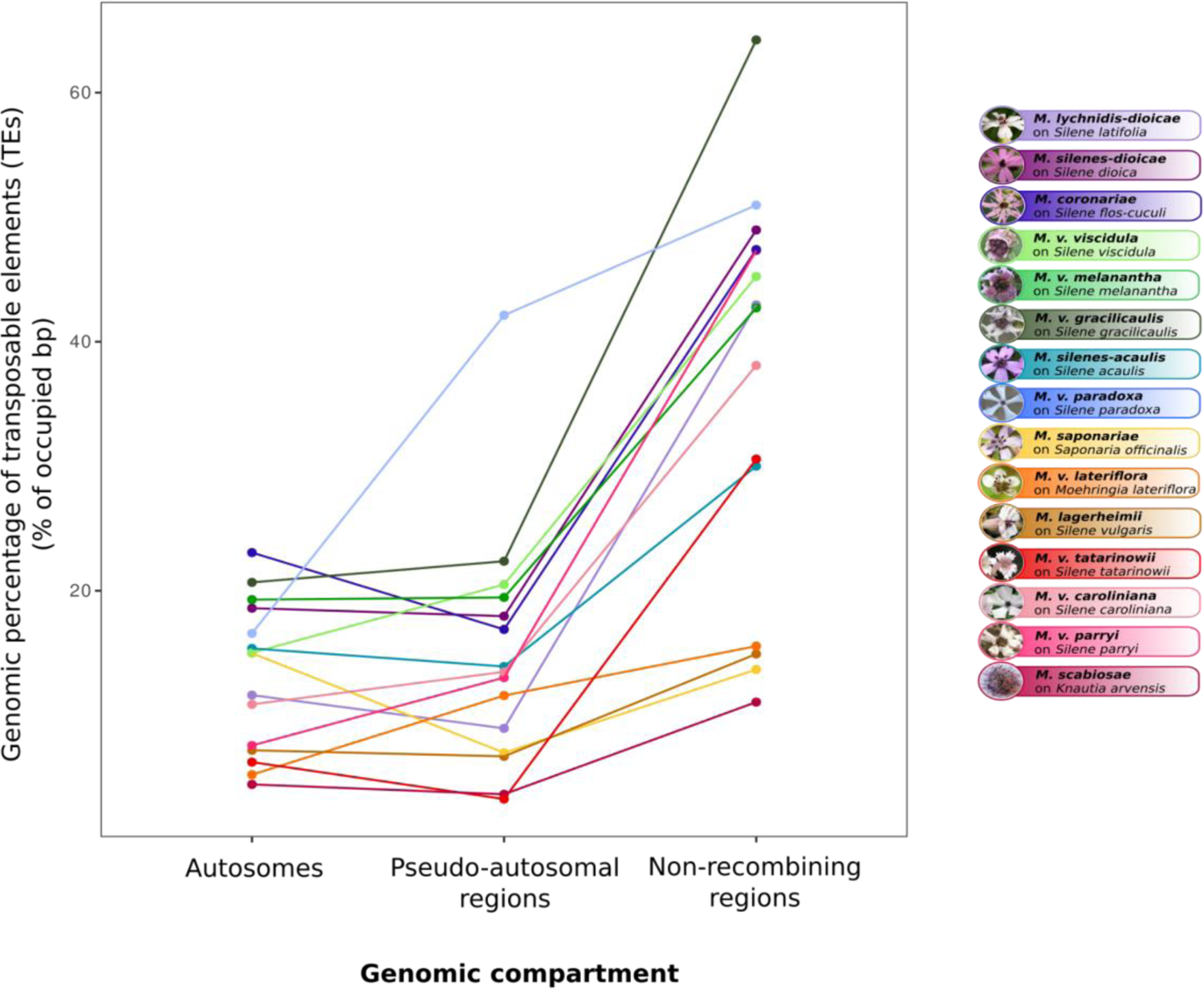
Transposable element (TE) proportions in the different genomic compartments of *Microbotryum* fungi. Total TE load per genomic compartment, i.e., autosomes (without the centromeres), pseudo-autosomal regions and non-recombining regions. The lines connect the values in the same species to facilitate comparisons. Each color represents a *Microbotryum* species.

**Supplementary Figure S5:**
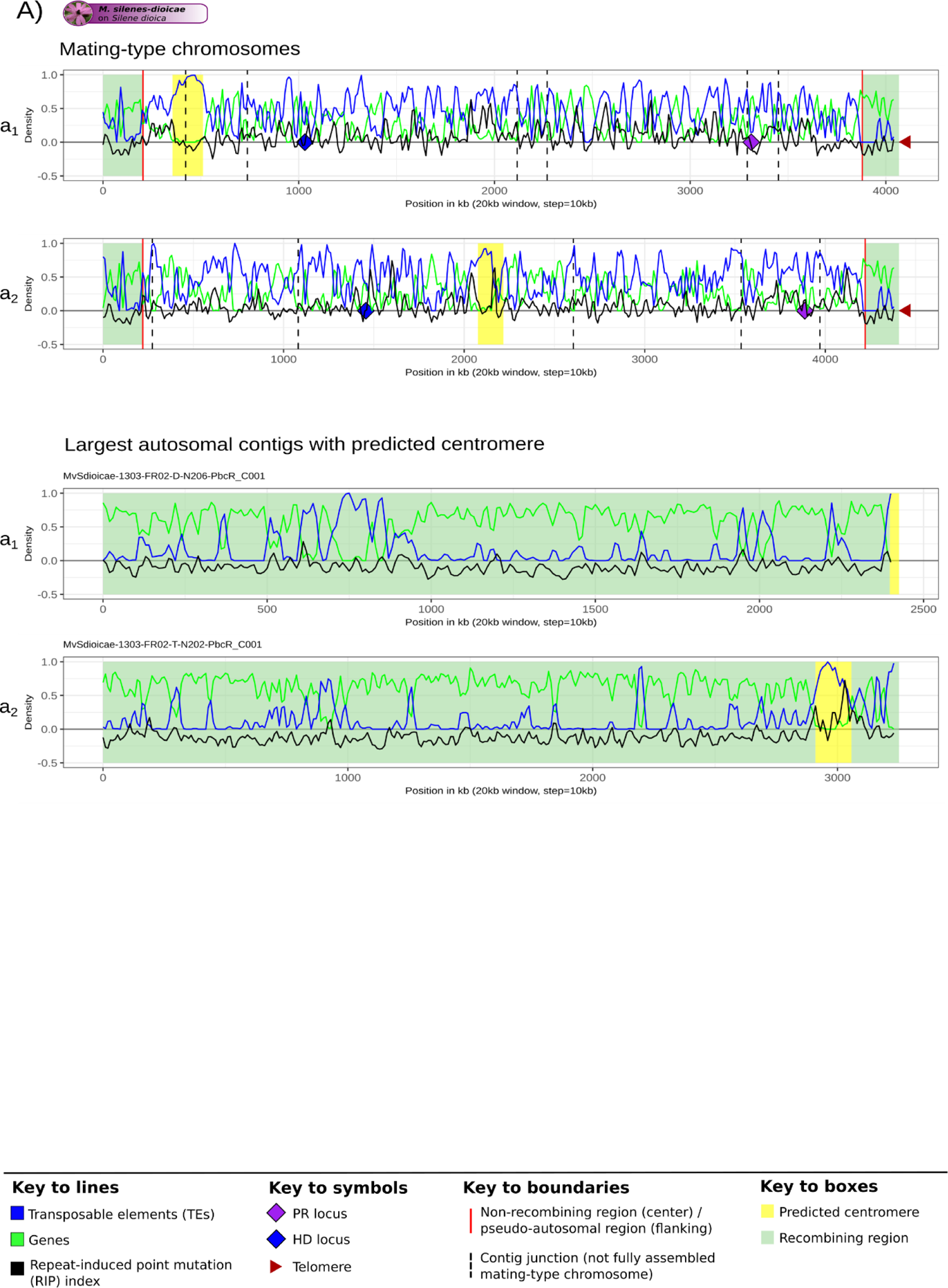
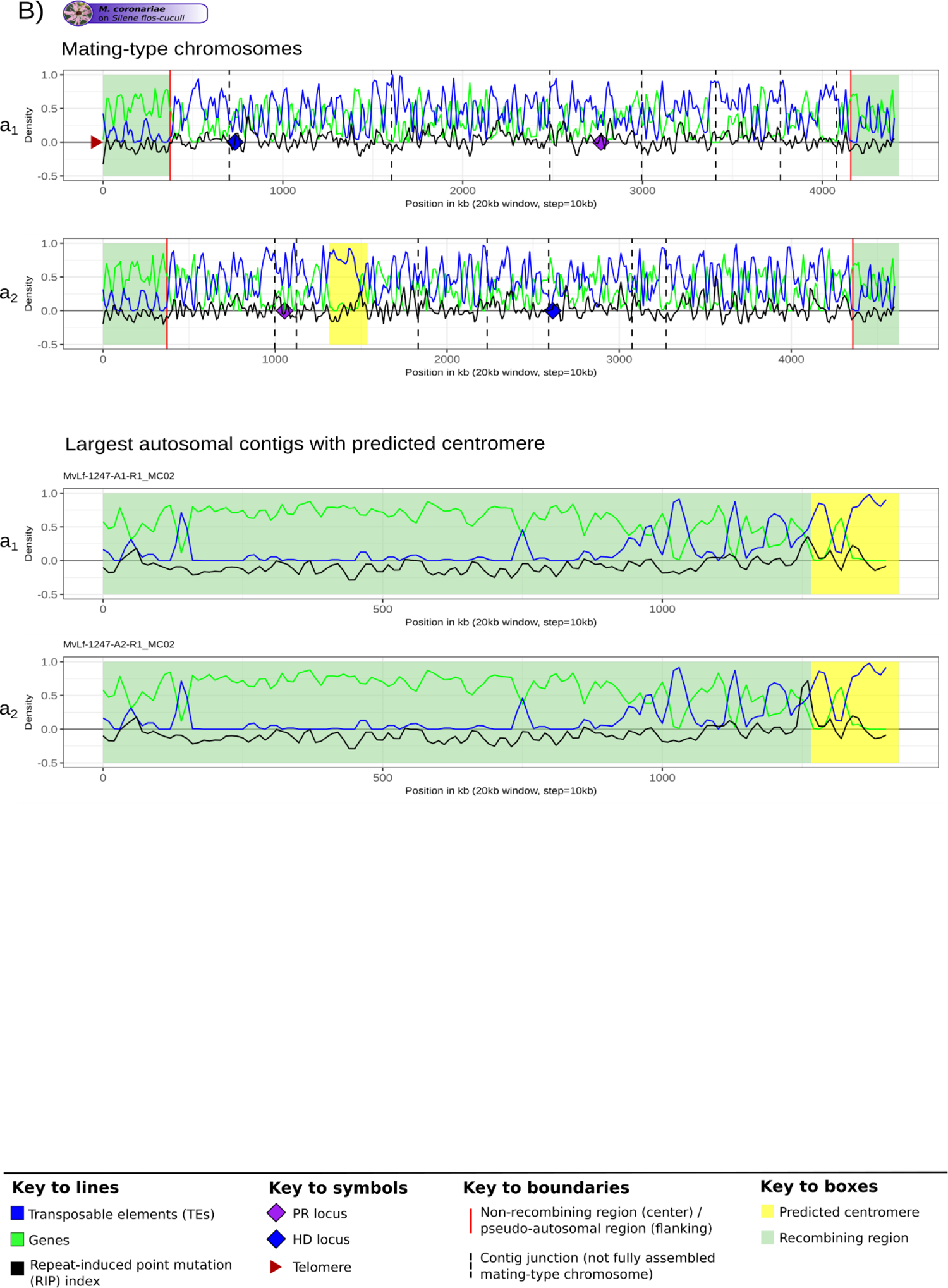
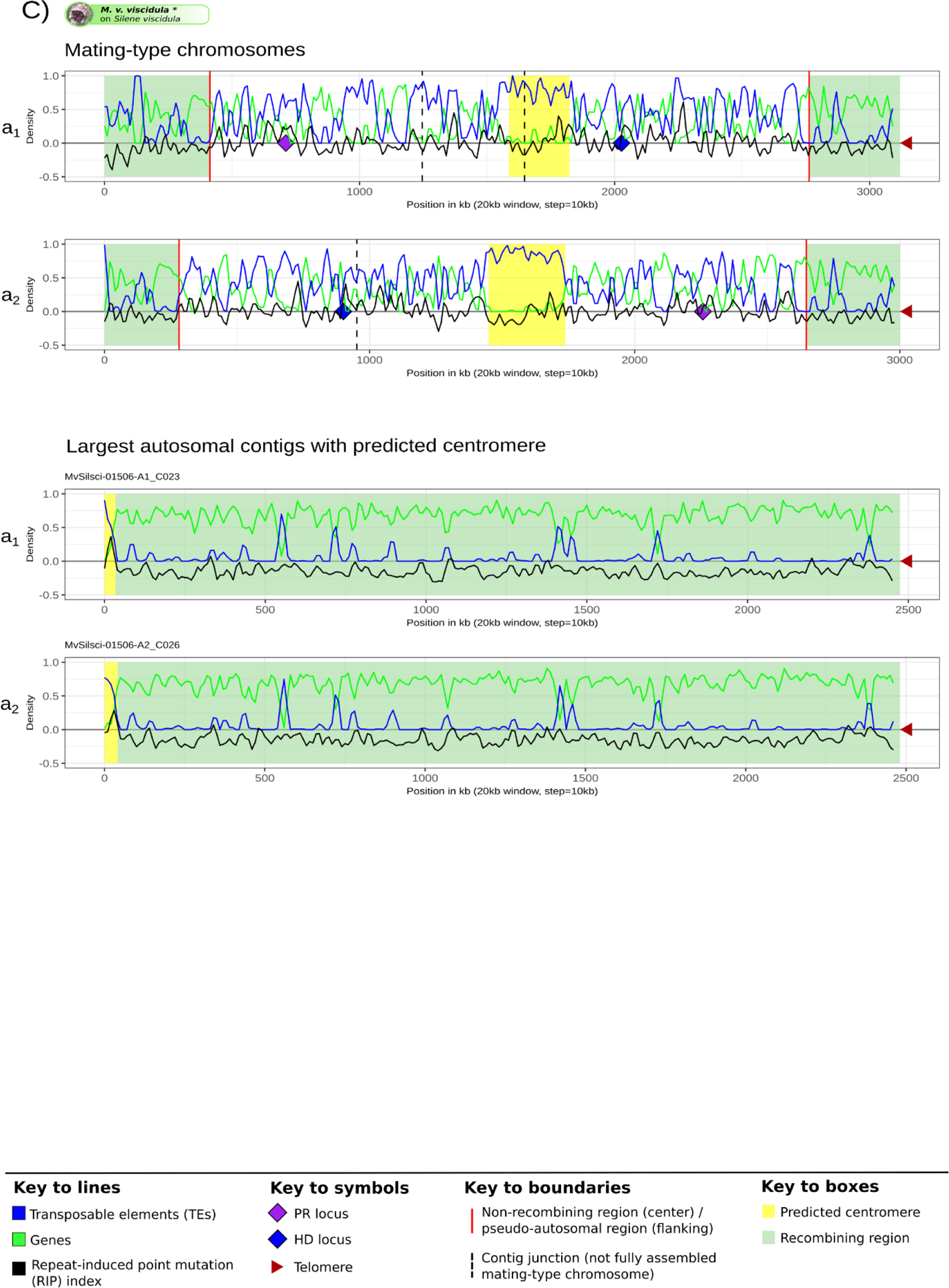
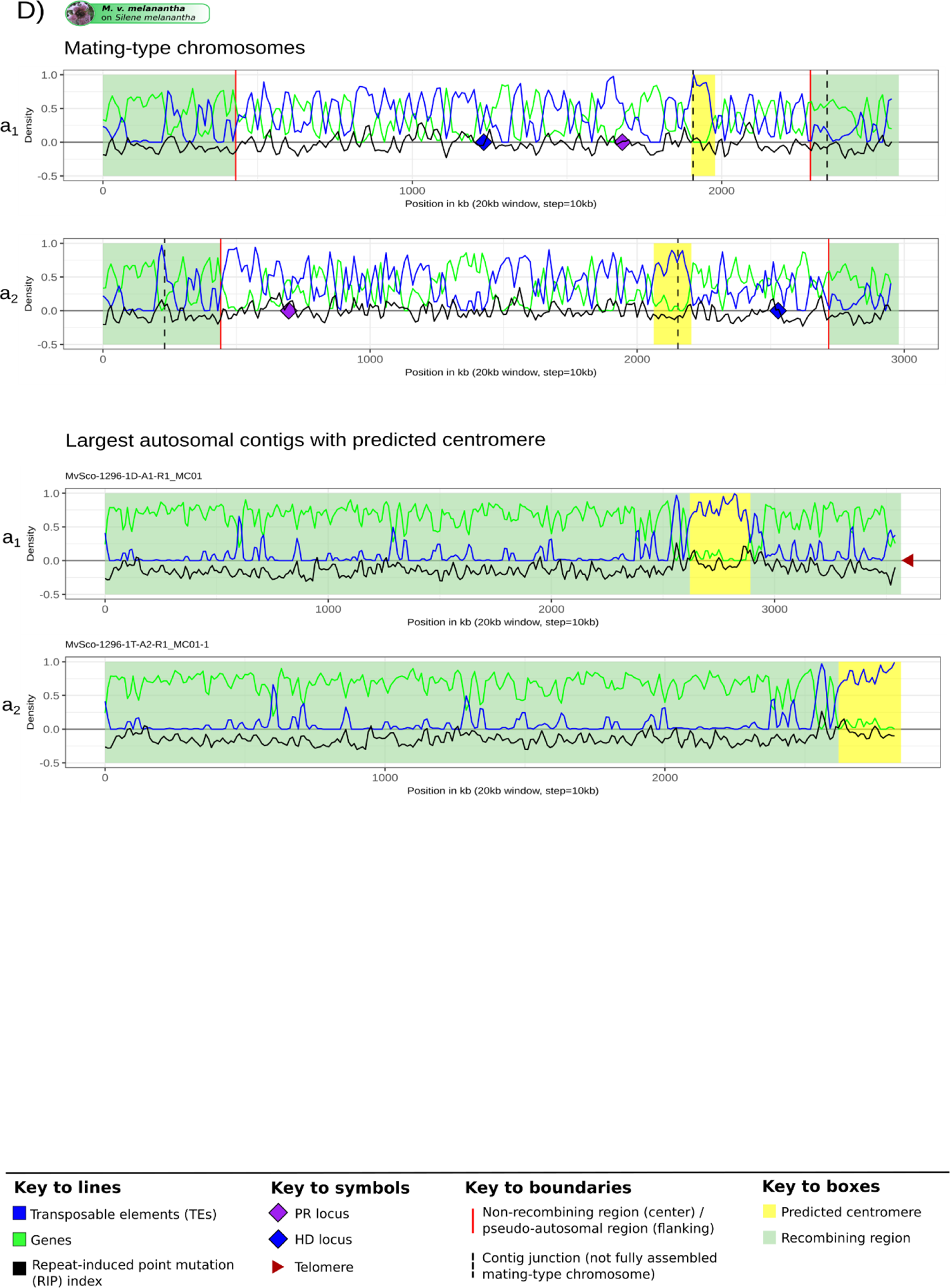
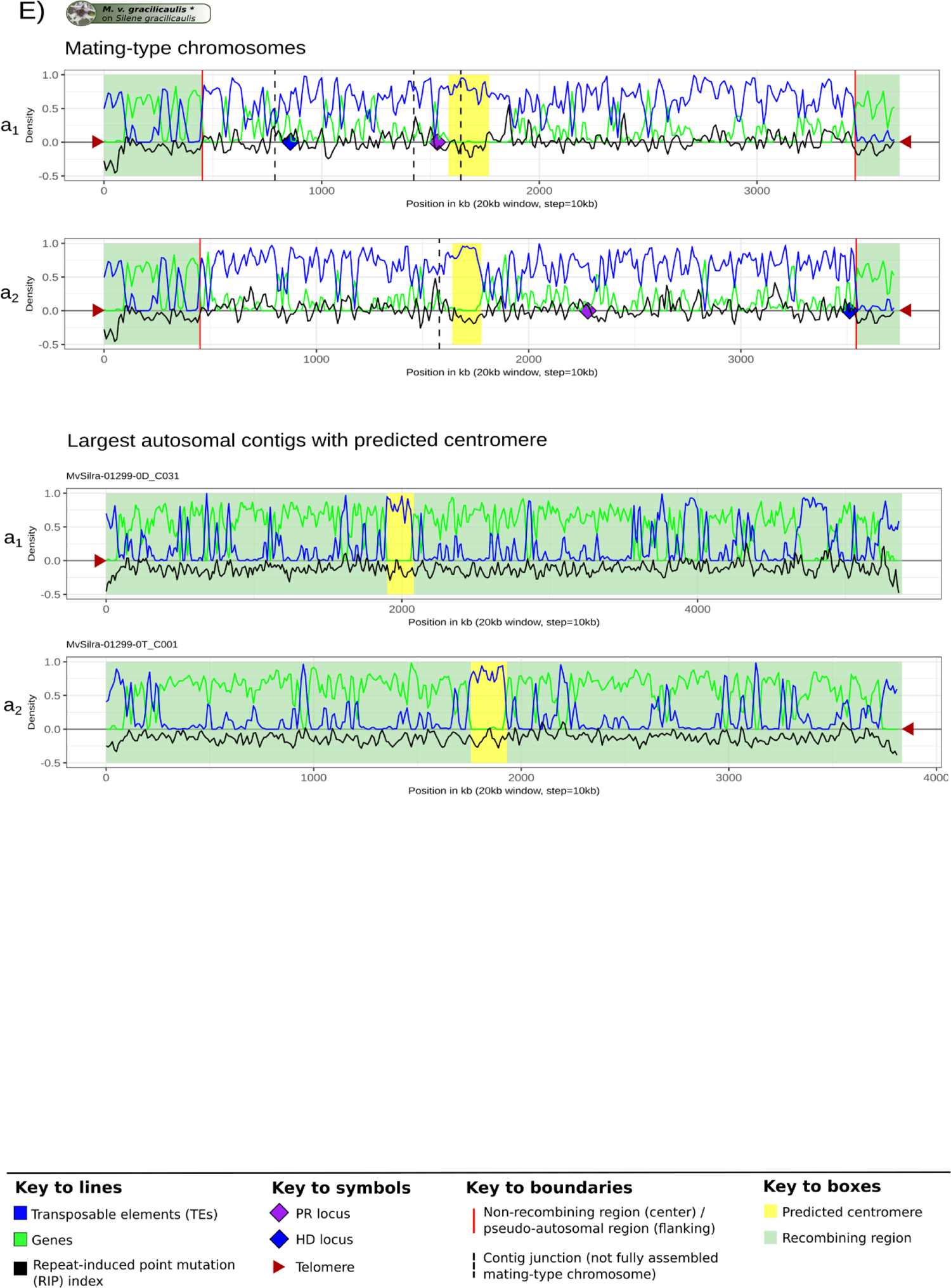
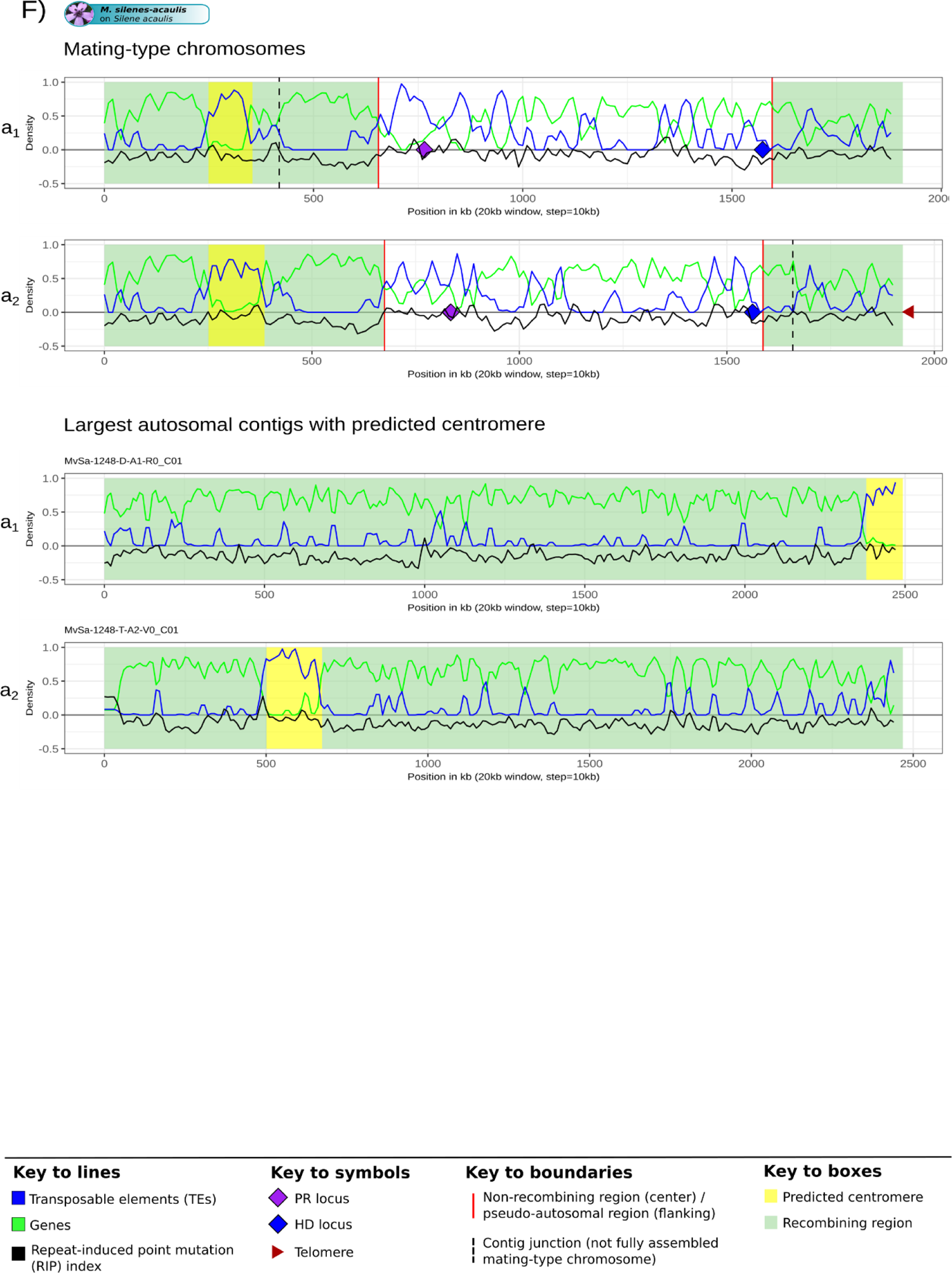
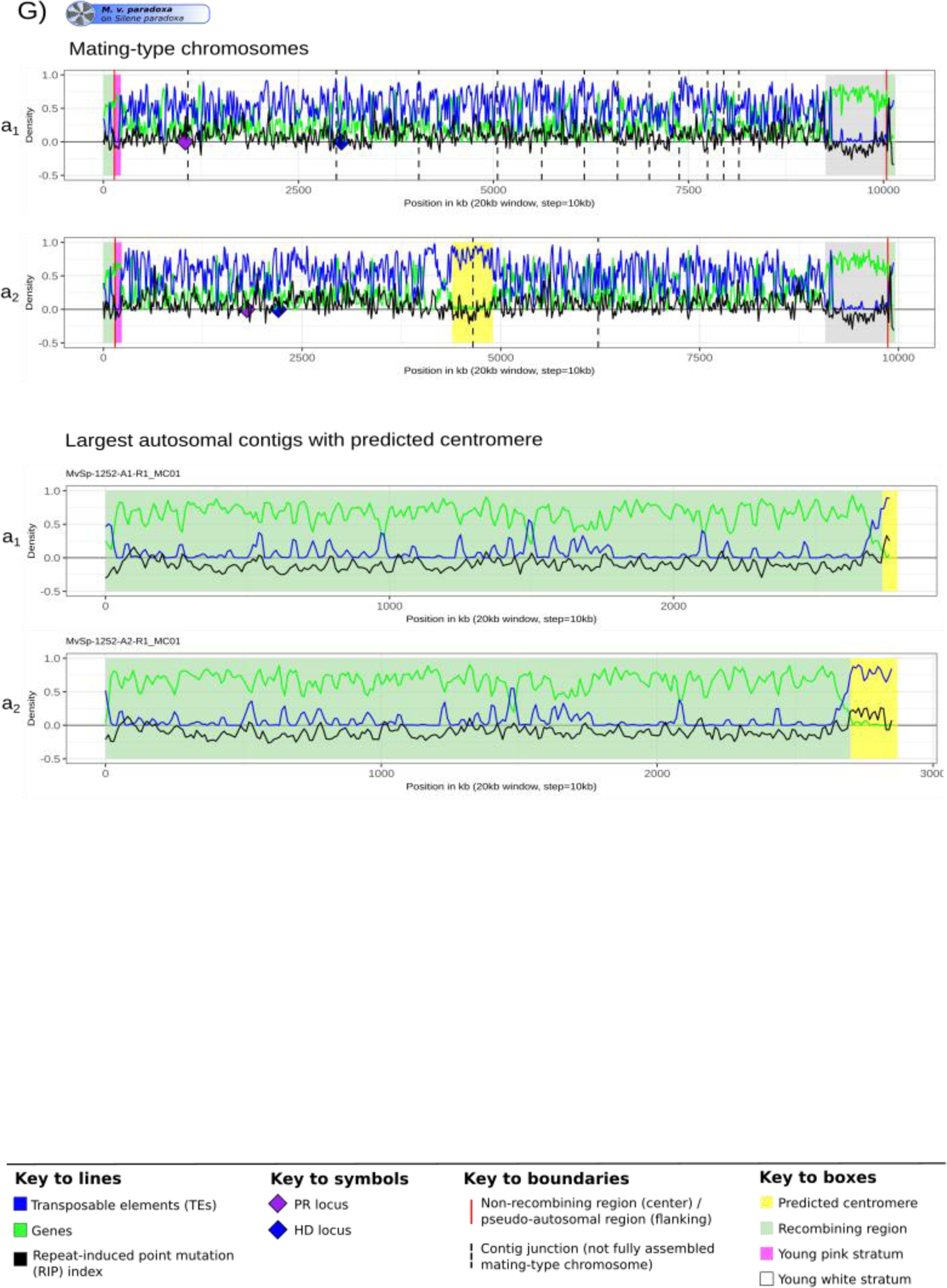
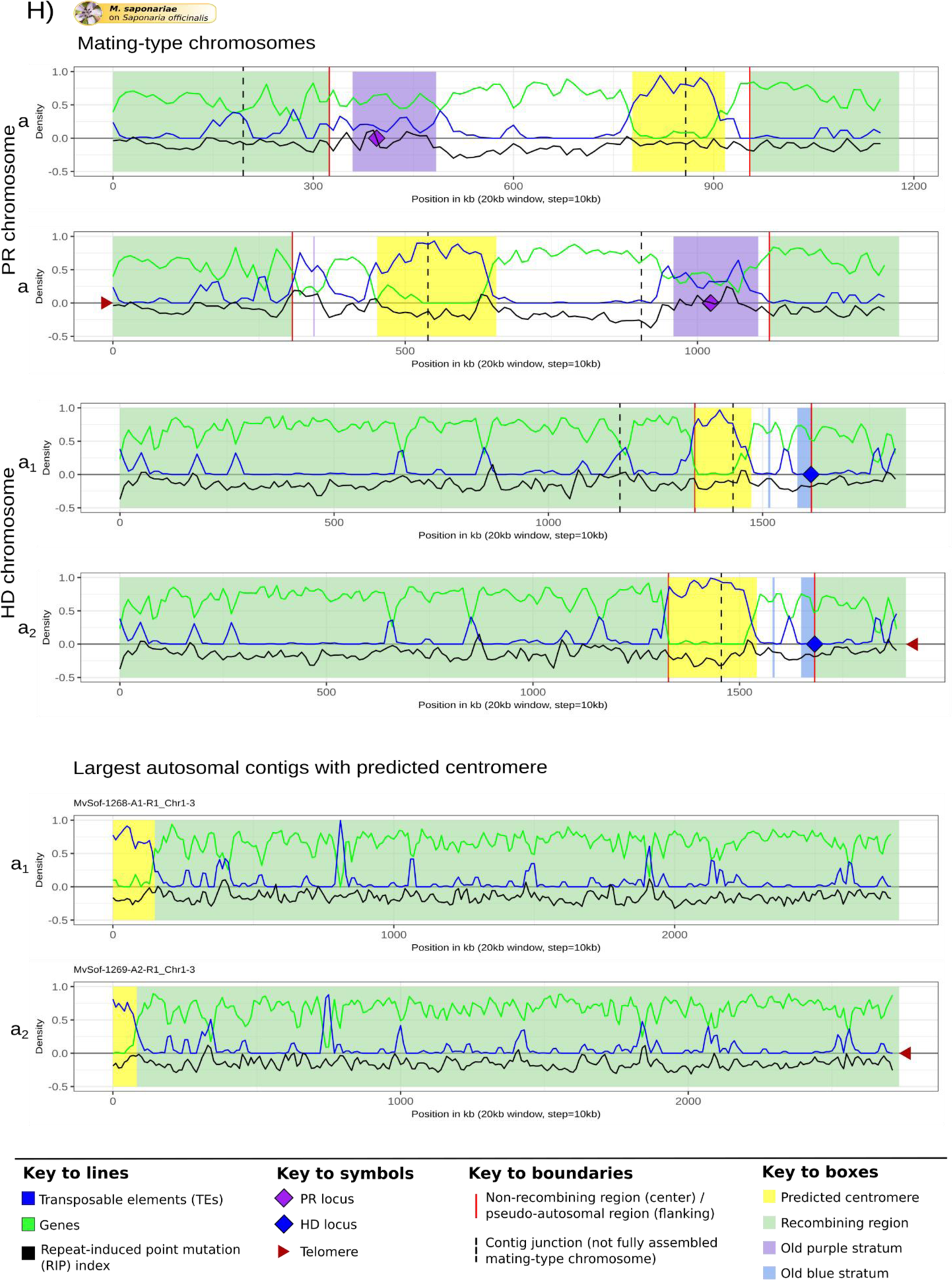
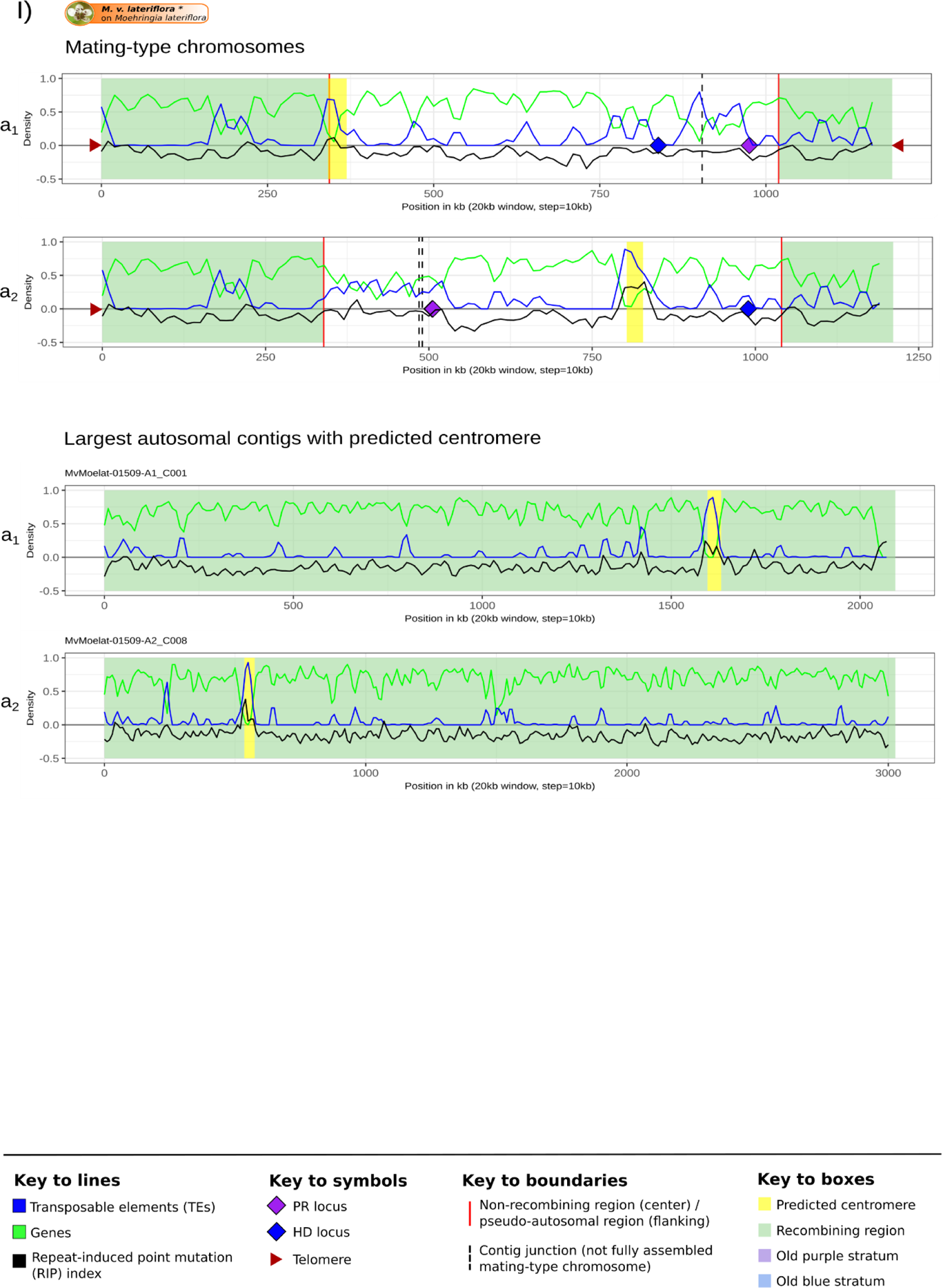
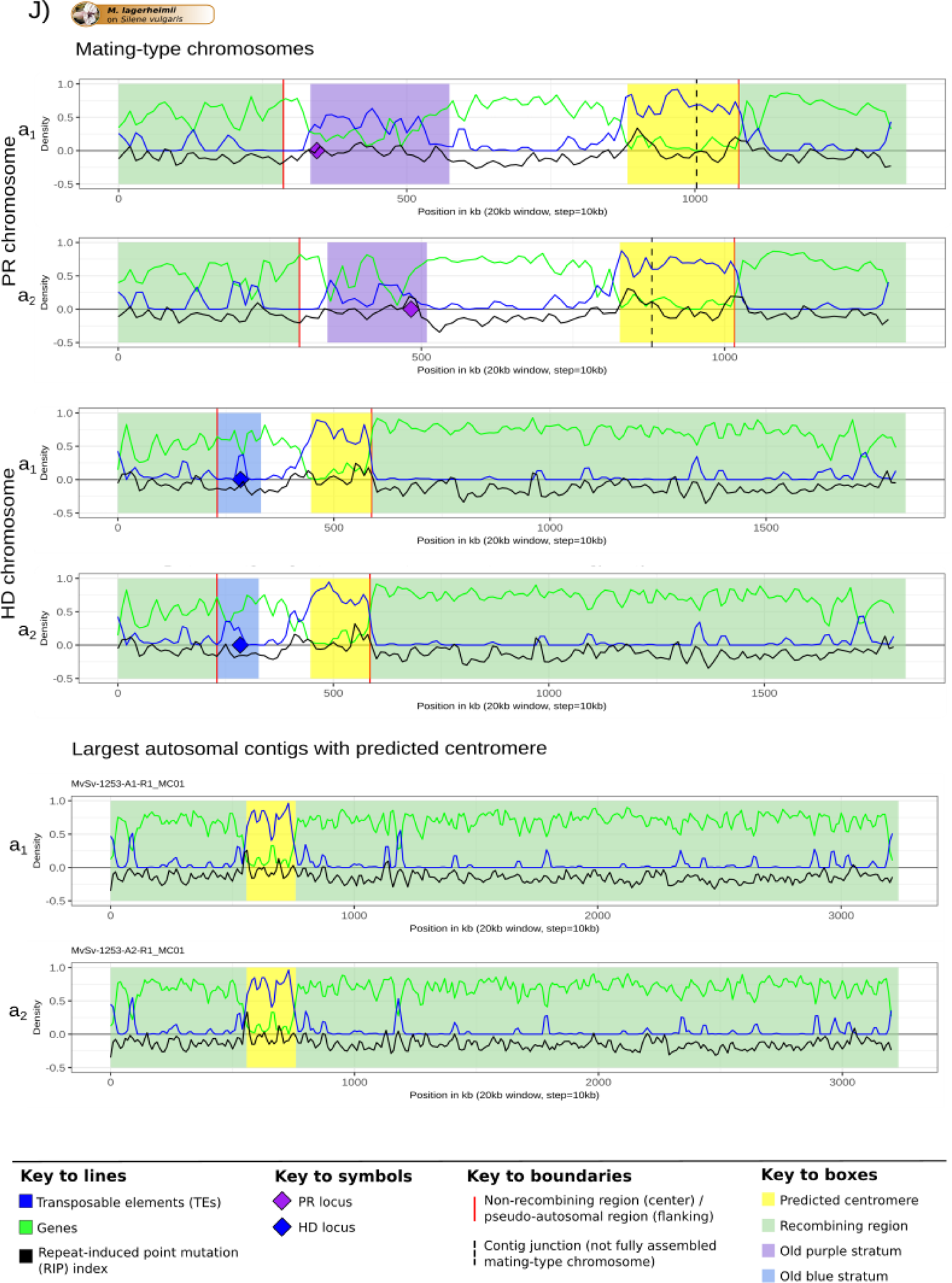
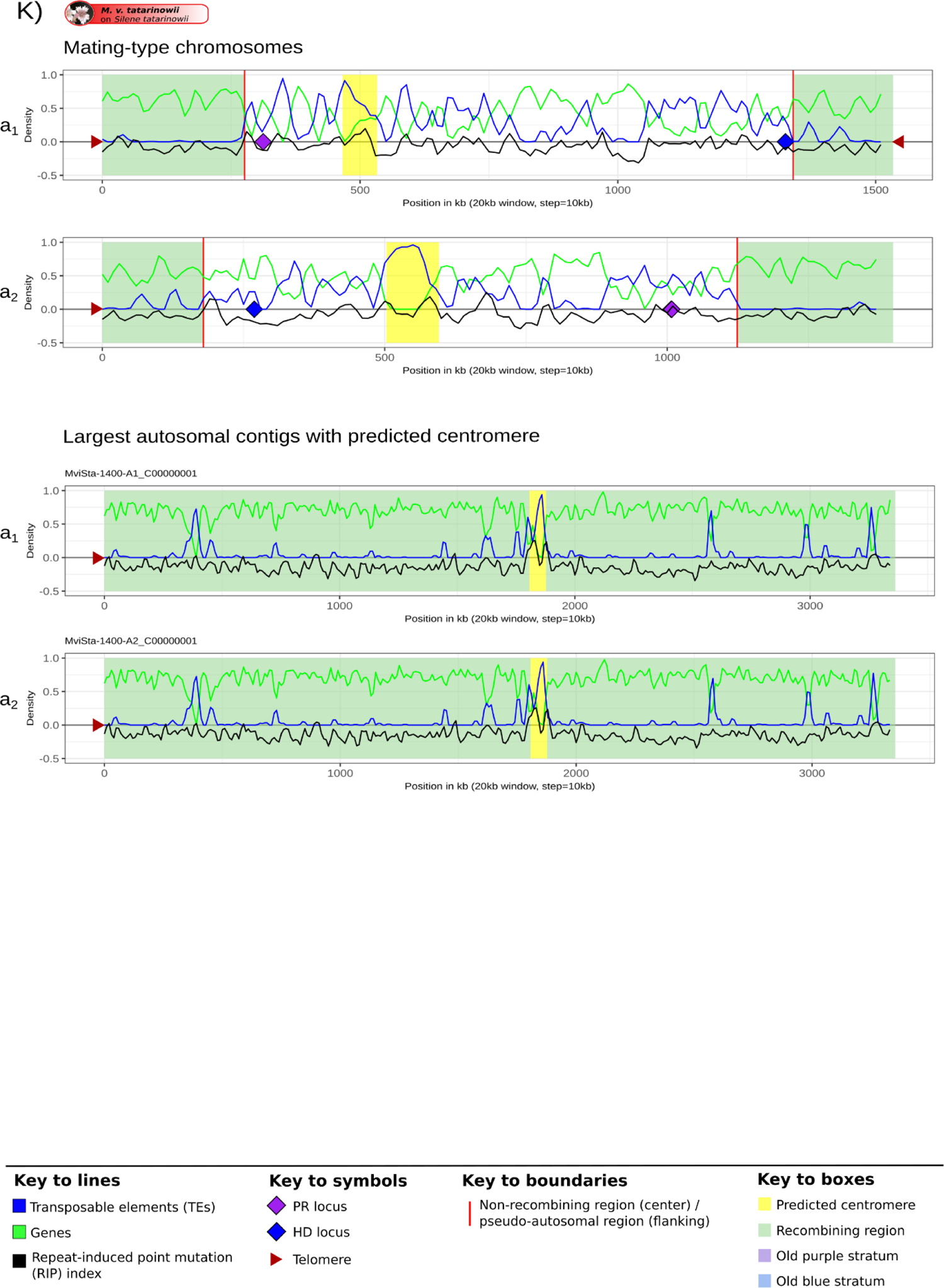
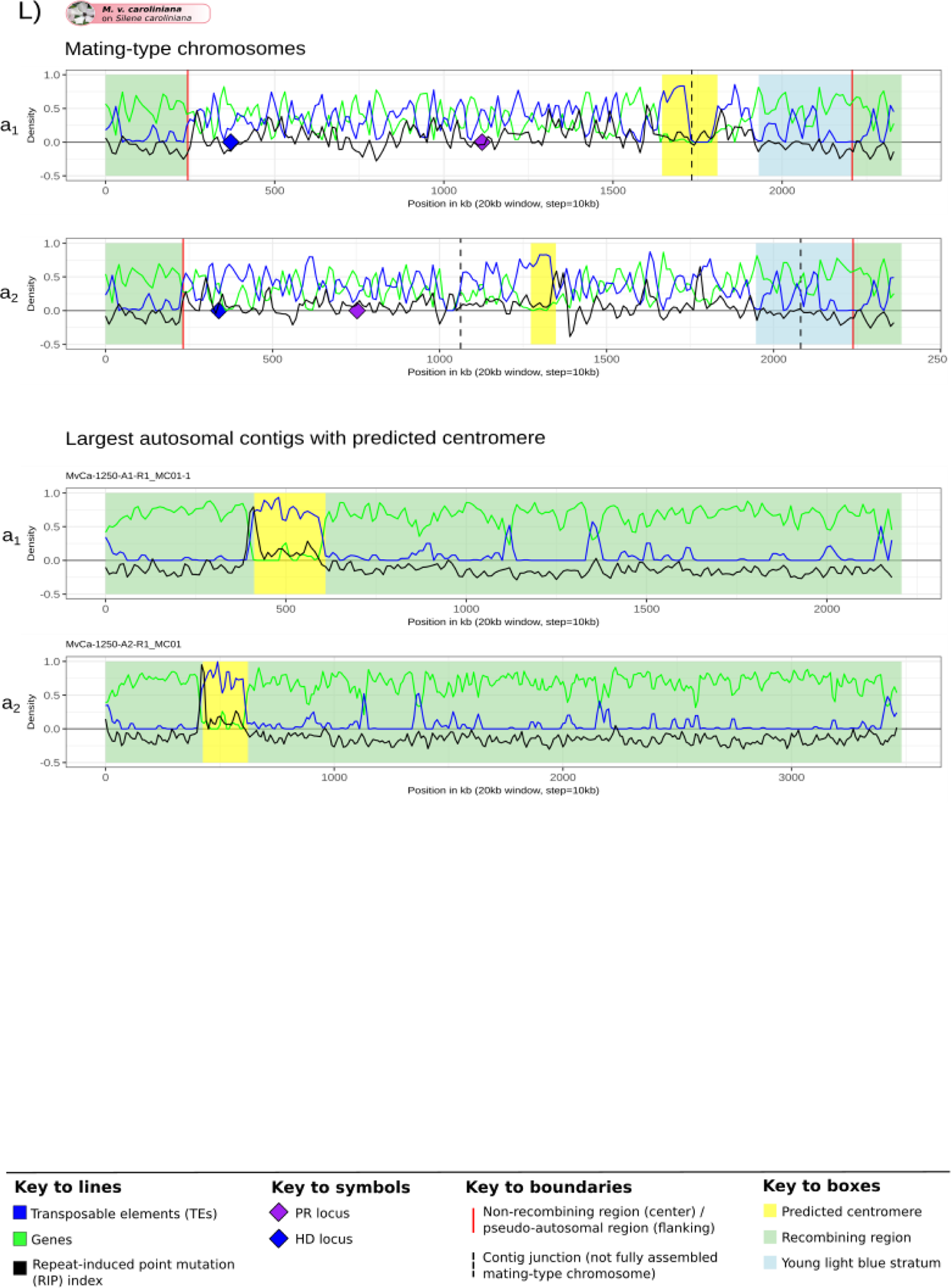
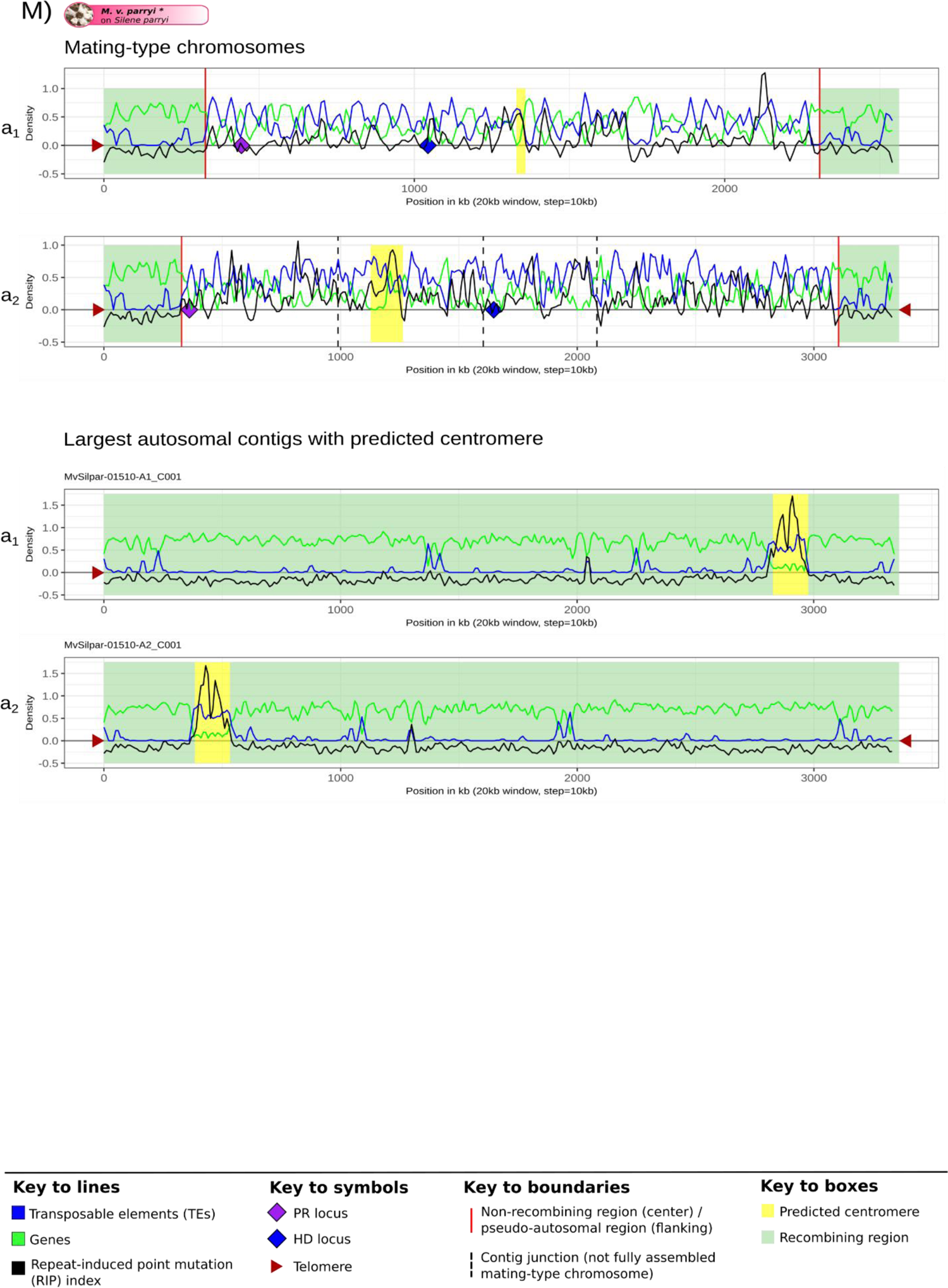
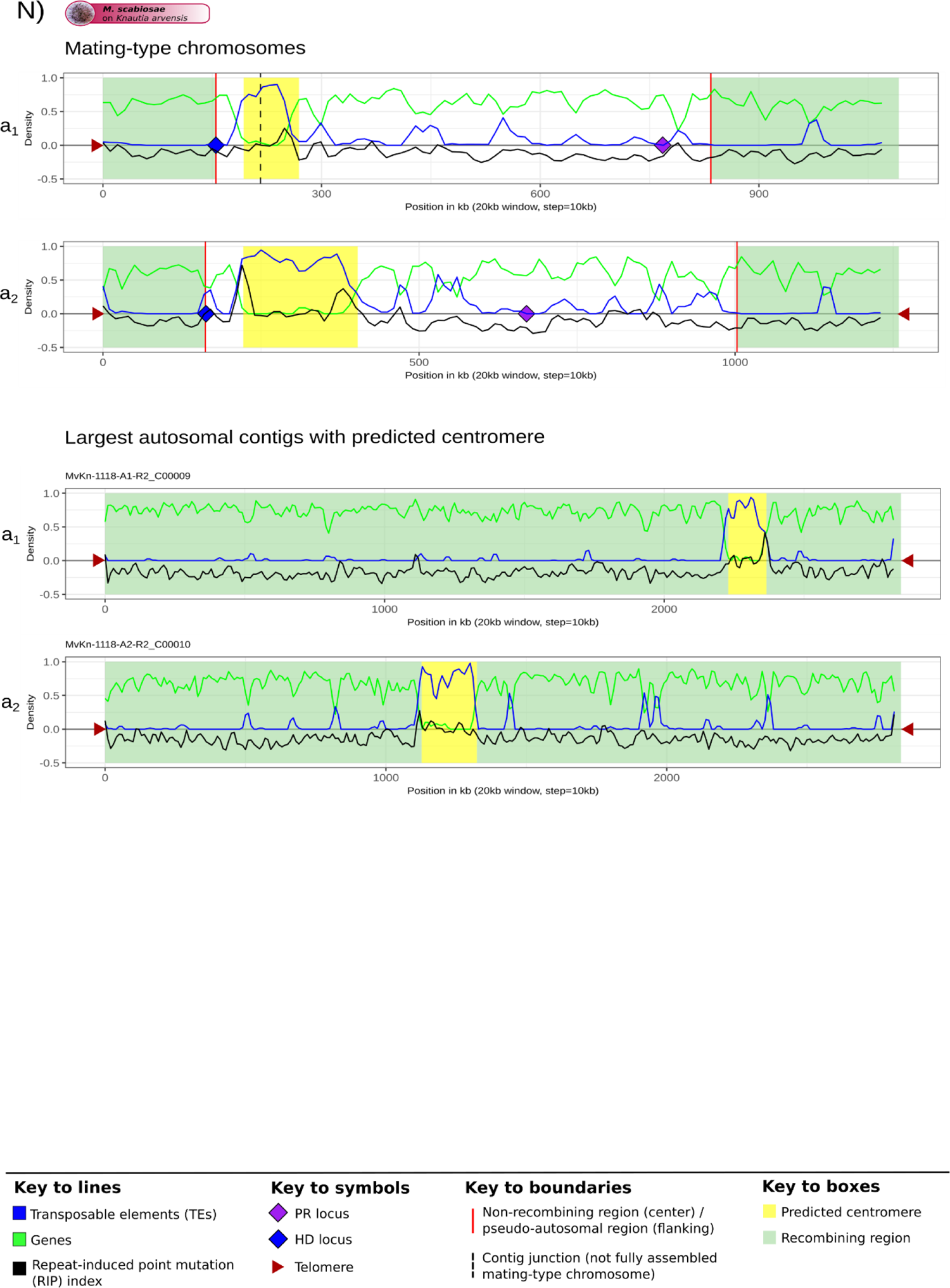
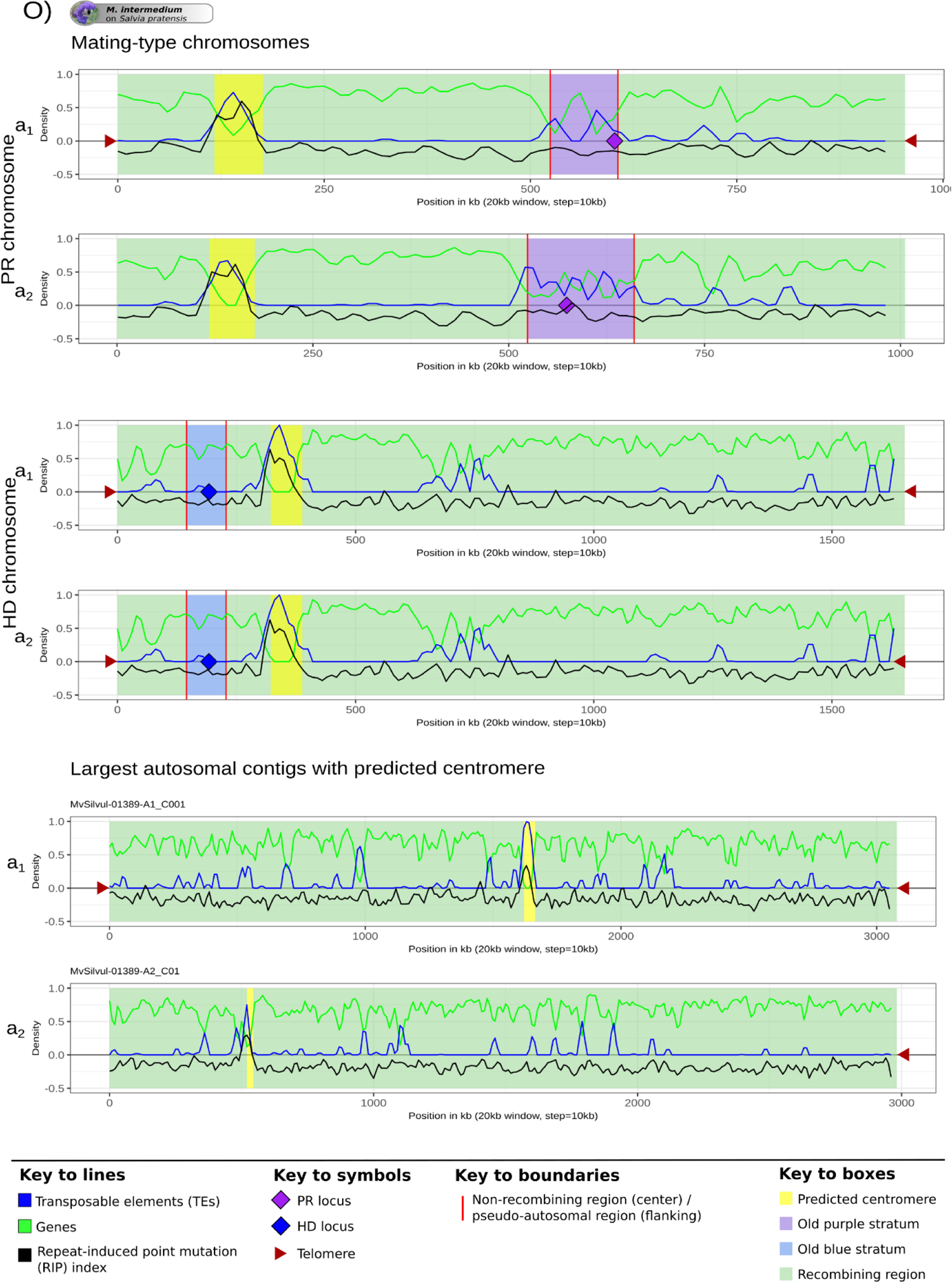
Density of transposable elements (TEs), genes and repeat-induced point mutations (RIP) along the mating-type chromosomes and the autosomes of *Microbotryum* genomes. Density of TEs, genes and RIP index were calculated in 20 kb non-overlapping windows. A RIP index greater than one indicates that the region is RIP-affected. Predicted centromeres are indicated in yellow. Recombining regions of the mating-type chromosomes (pseudo-autosomal regions) and the autosomes (largest autosomal contig harboring a predicted centromere in each genome) are highlighted in pale green. Non-recombining regions are delimited by red lines. Non-rearranged evolutionary strata are displayed in their corresponding color (see legend). Non-fully assembled mating-type chromosomes have been joined and the junctions are indicated by black dashed lines. Pheromone-receptor (PR) and homeodomain (HD) loci are indicated by purple and blue diamonds, respectively. Identified telomeres are indicated by brown triangles.

**Supplementary Figure S6:**
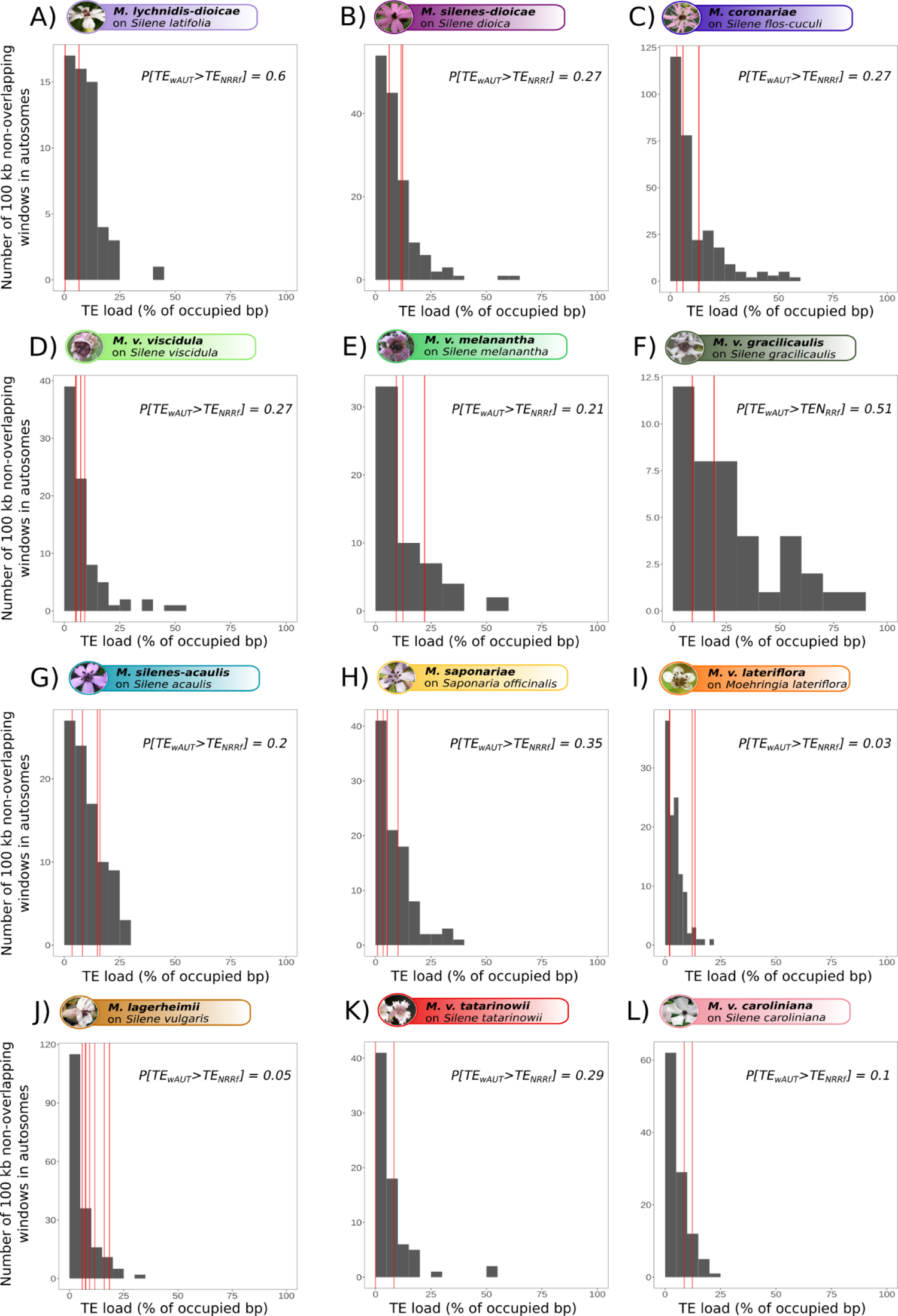
Transposable element (TE) load at the margin of the non-recombining regions compared to the distribution in recombining autosomal regions. The average values of TE content in the regions flanking the non-recombining region are indicated by red lines and the ones in autosomal windows constitute the grey distribution. The probability of finding a higher TE content in a 100 kb window on autosomes than in the 100 kb region directly flanking the non-recombining region (P[TE_wAUT_>TE_NRRf_]) is reported in the top right corner of each panel. The panels correspond to different species, and there are four or eight red lines (individual flanking regions of non-recombining regions) per species: one a_1_ and one a_2_ genome per species, and for each haploid genome one or two mating-type chromosomes, so four or eight flanking regions. Three species are not represented because their pseudo-autosomal regions were smaller than 100 kb (*Microbotryum violaceum paradoxa*) or their autosome assemblies were too fragmented (*M. v. parryi* and *M. scabiosae*).

**Supplementary Figure S7:**
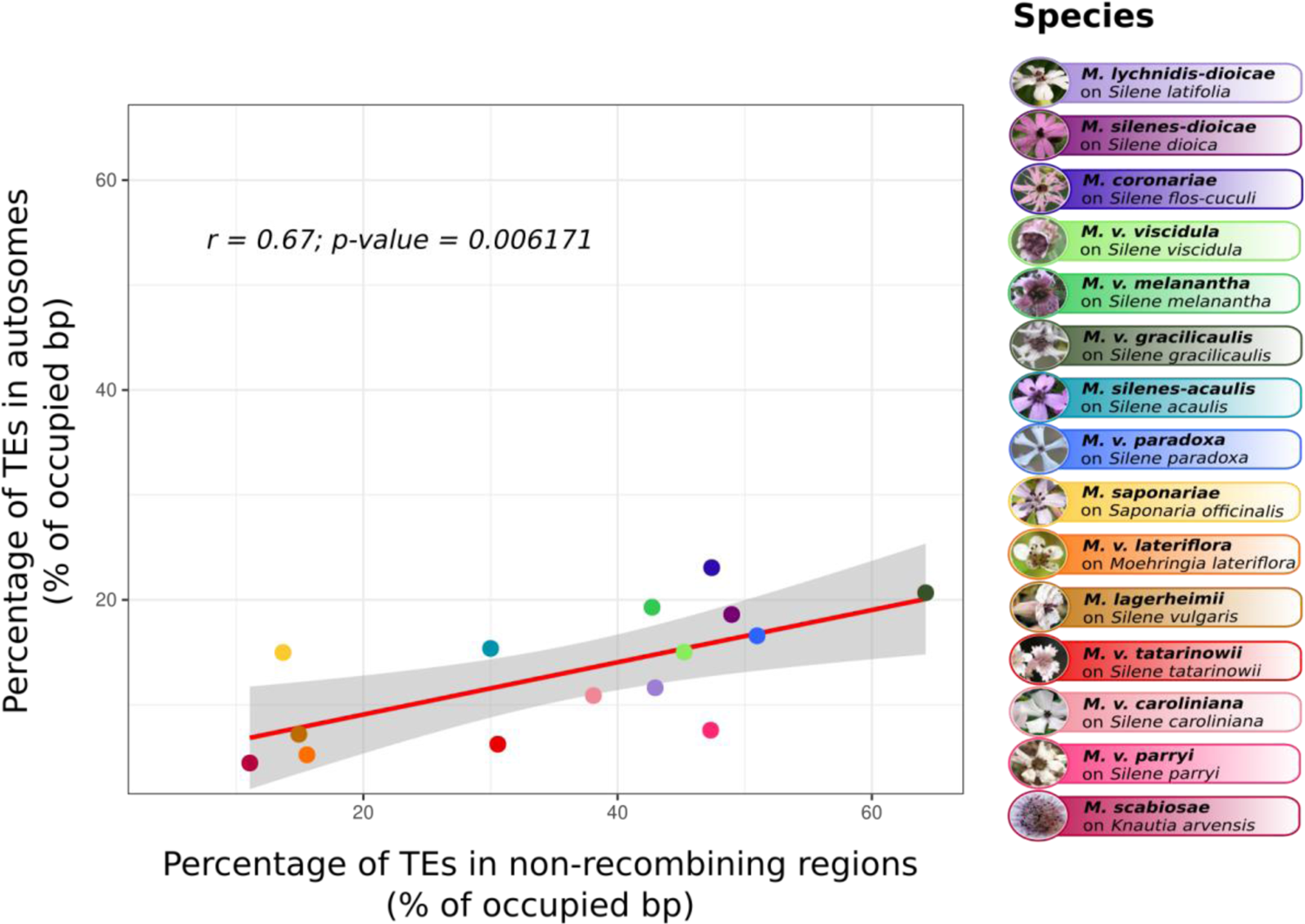
Correlation between the proportion of base pairs occupied by transposable elements (TEs) in the non-recombining regions and the autosomes.

**Supplementary Figure S8:**
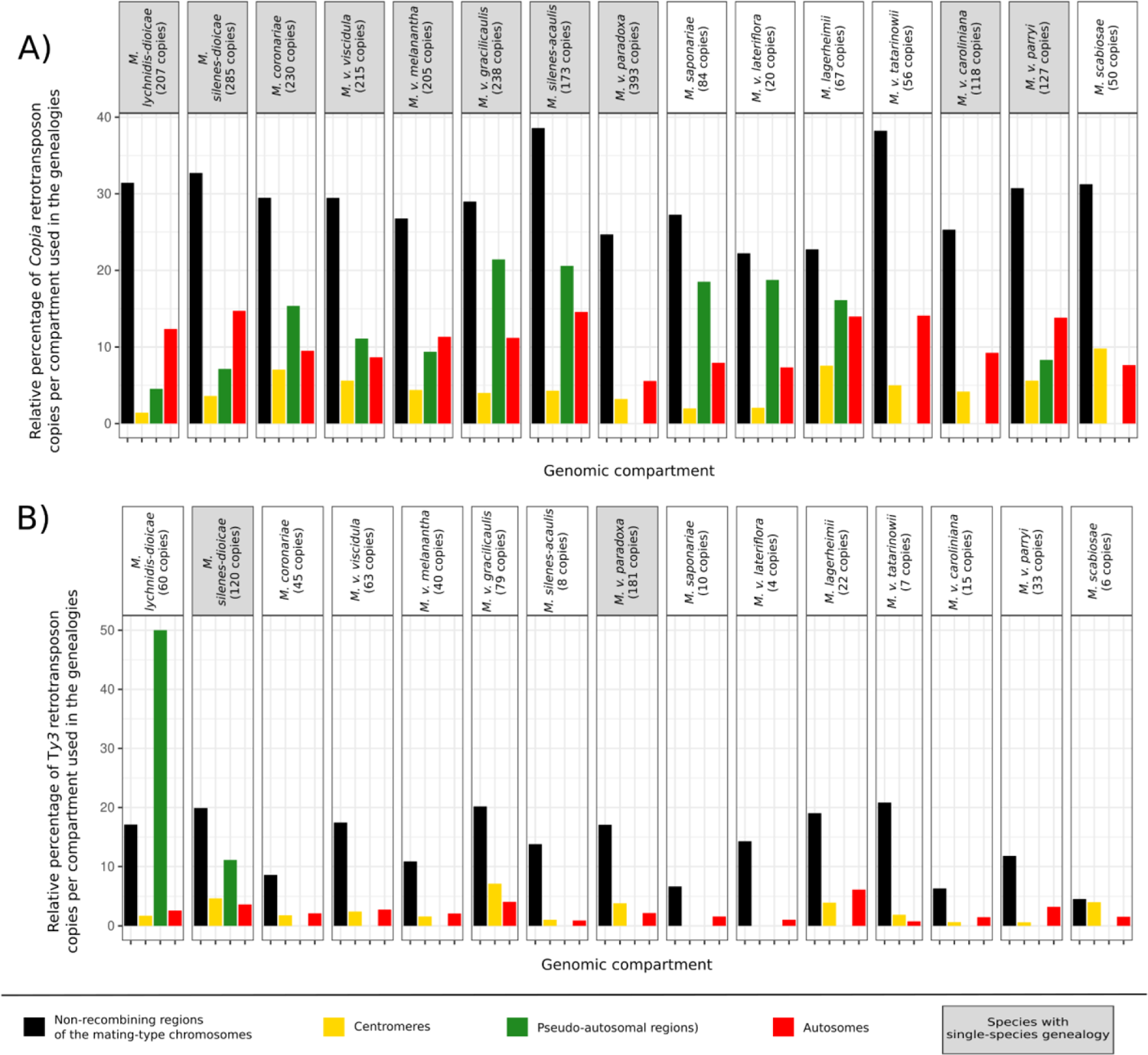
Relative percentage and number of transposable element (TE) sequences per genomic compartment used for the genealogies based on their 5’-LTR sequences, for *Copia* (A) and *Ty3* (B) elements. The total number of 5’-LTR sequences available per species is indicated in brackets under the species name. Single-species genealogies were only built for species with grey filled rectangles around their names.

**Supplementary Figure S9:**
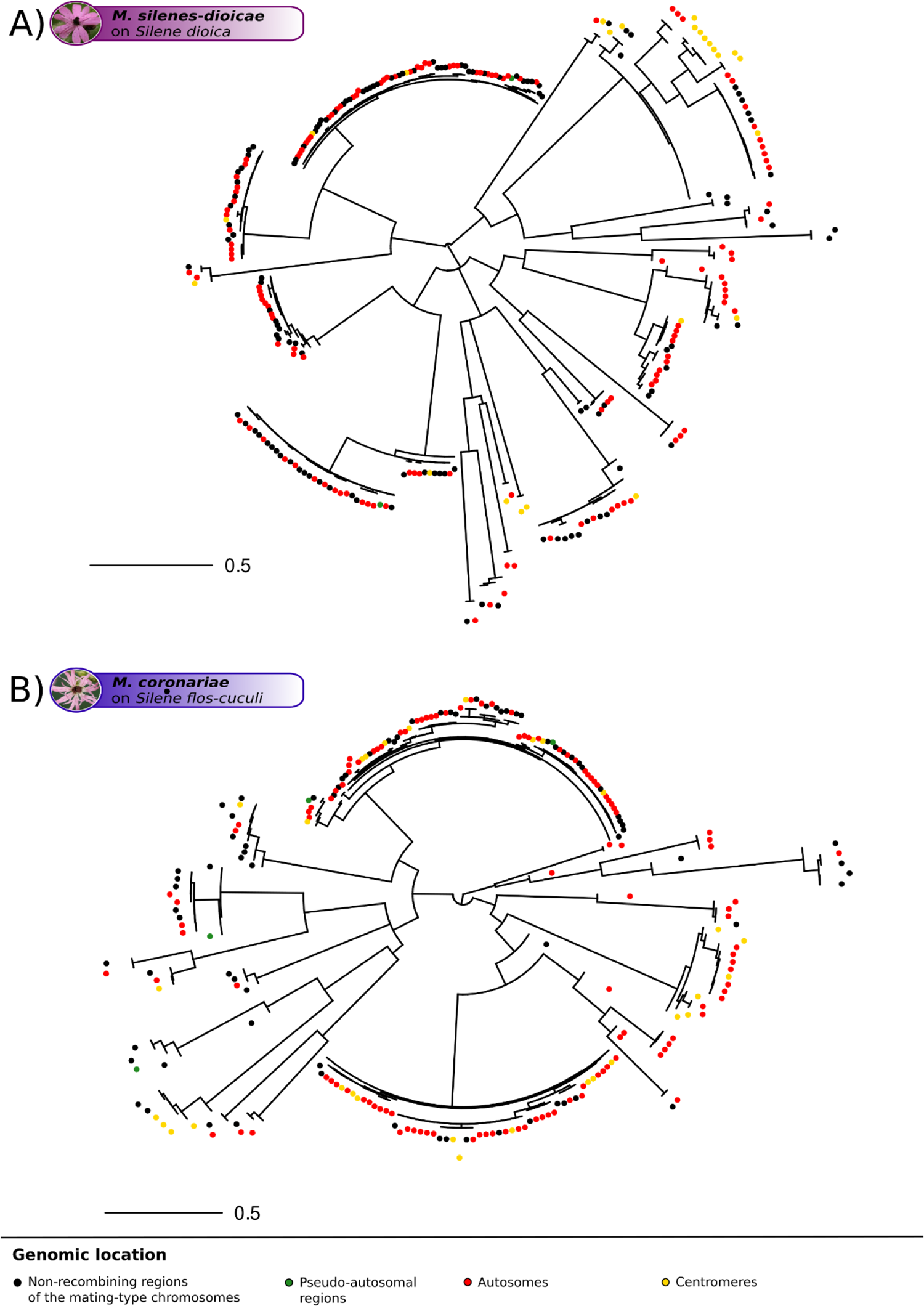
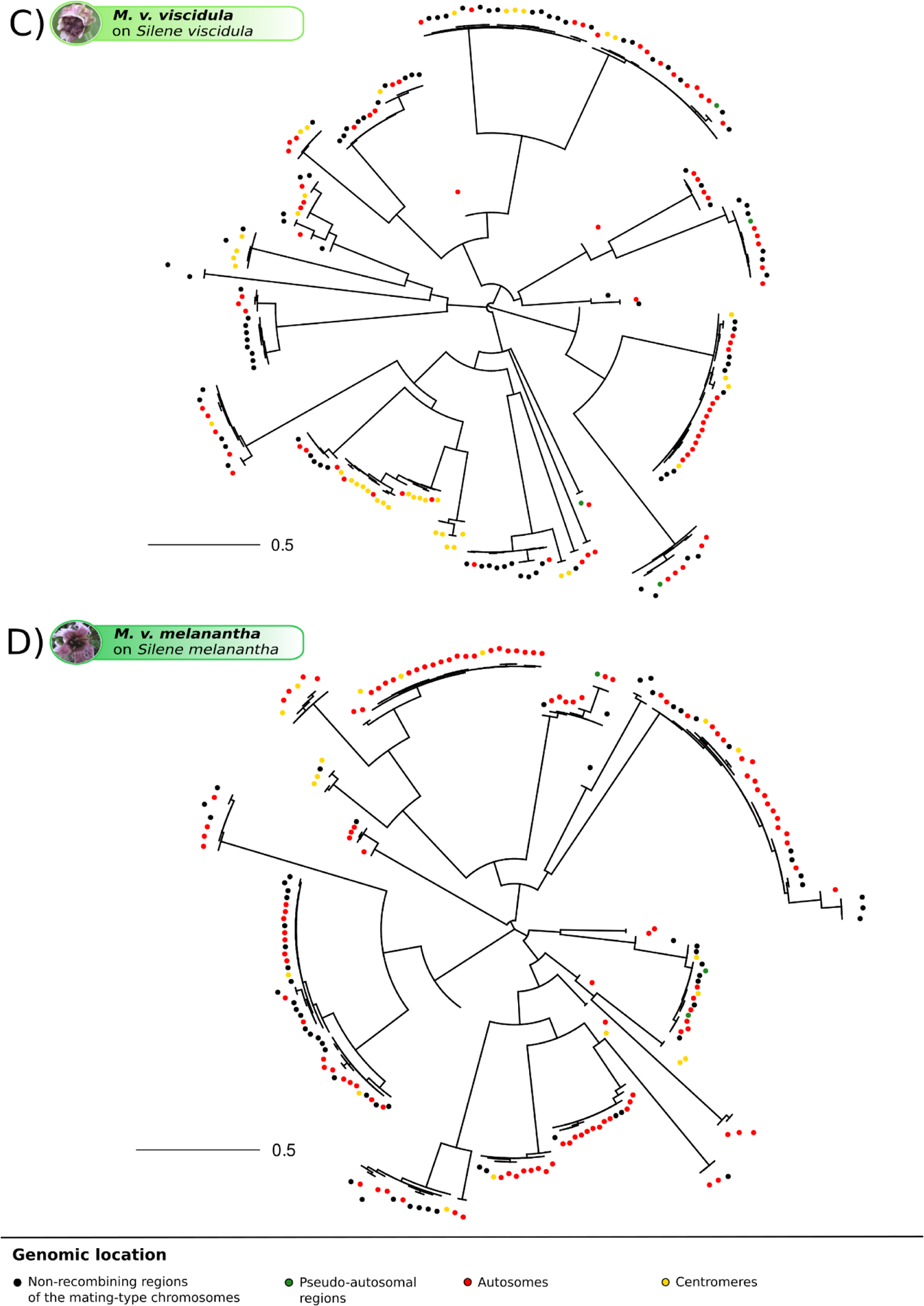
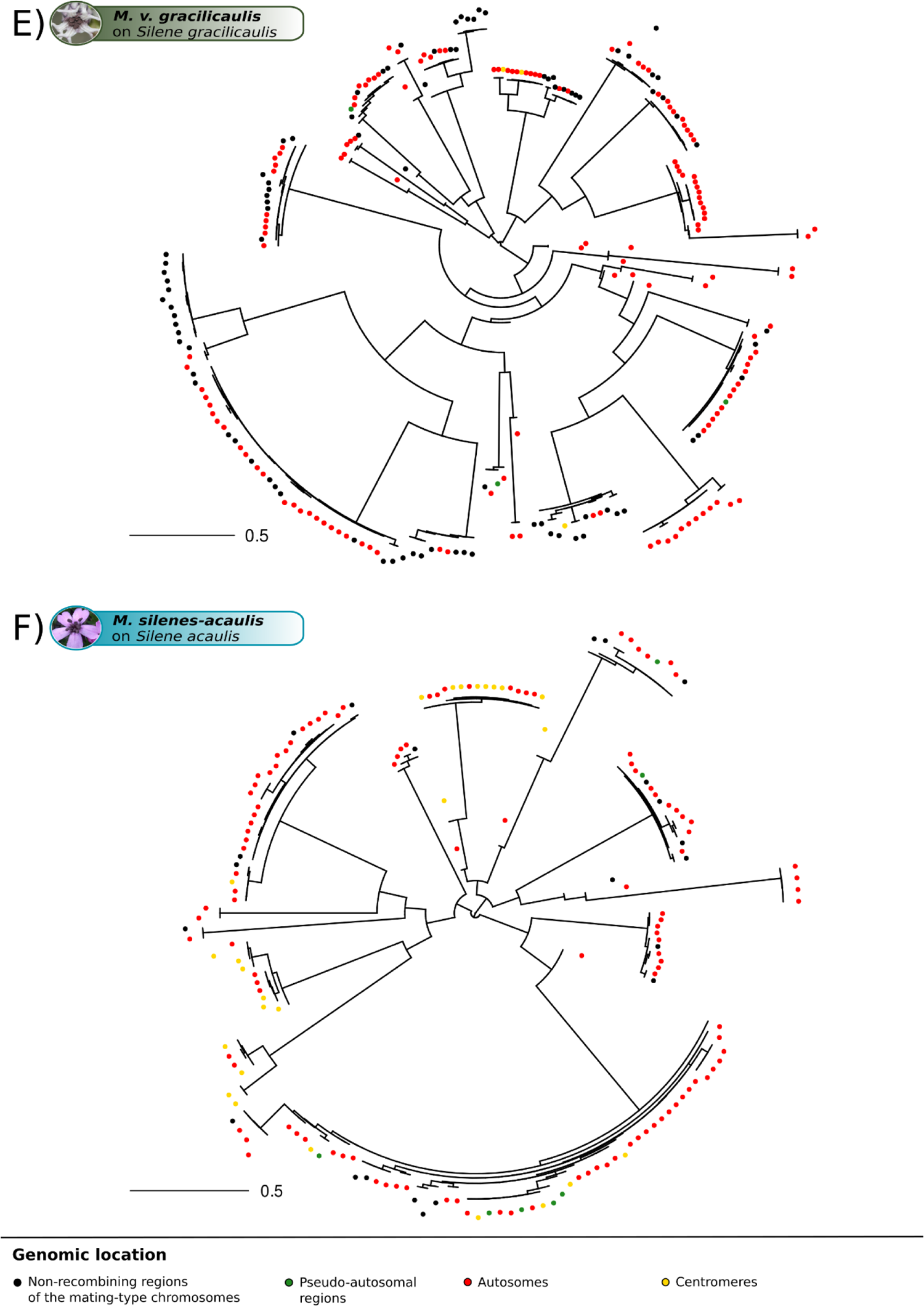
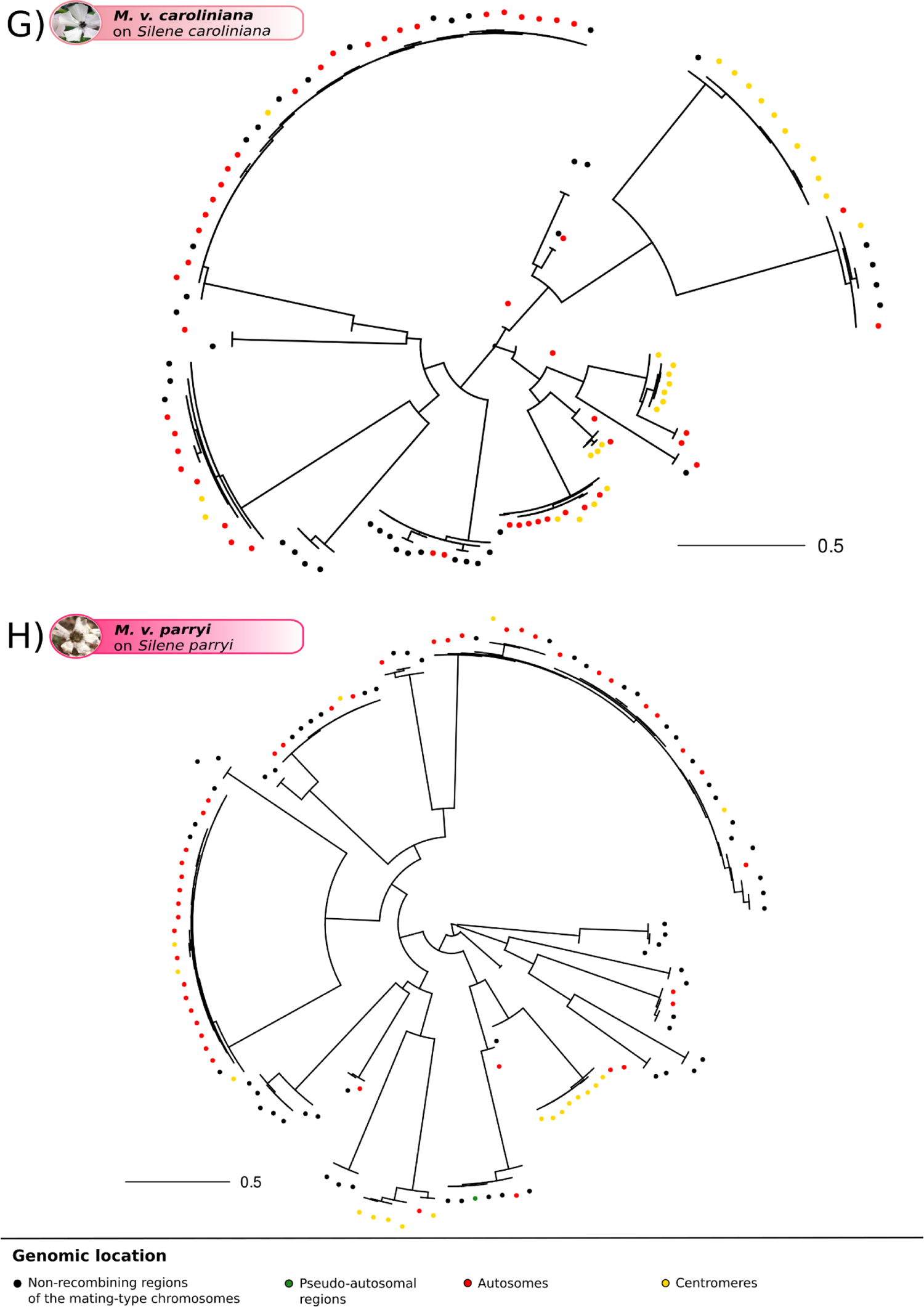
Single-species genealogies of *Copia* retroelement copies in *Microbotryum* genomes. Single-species genealogies of *Copia* copies in *Microbotryum silenes-dioicae* **(A)**, *M. coronariae* **(B)**, *M. violaceum viscidula* **(C)**, *M. v. melanantha* **(D)**, *M. v. gracilicaulis* **(E)**, *M. silenes-acaulis* **(F)**, *M. v. caroliniana* **(G)** and *M. v. parryi* **(H)**. The color of the dots at the tip of the branches corresponds to the genomic location of the transposable element (TE) copies.

**Supplementary Figure S10:**
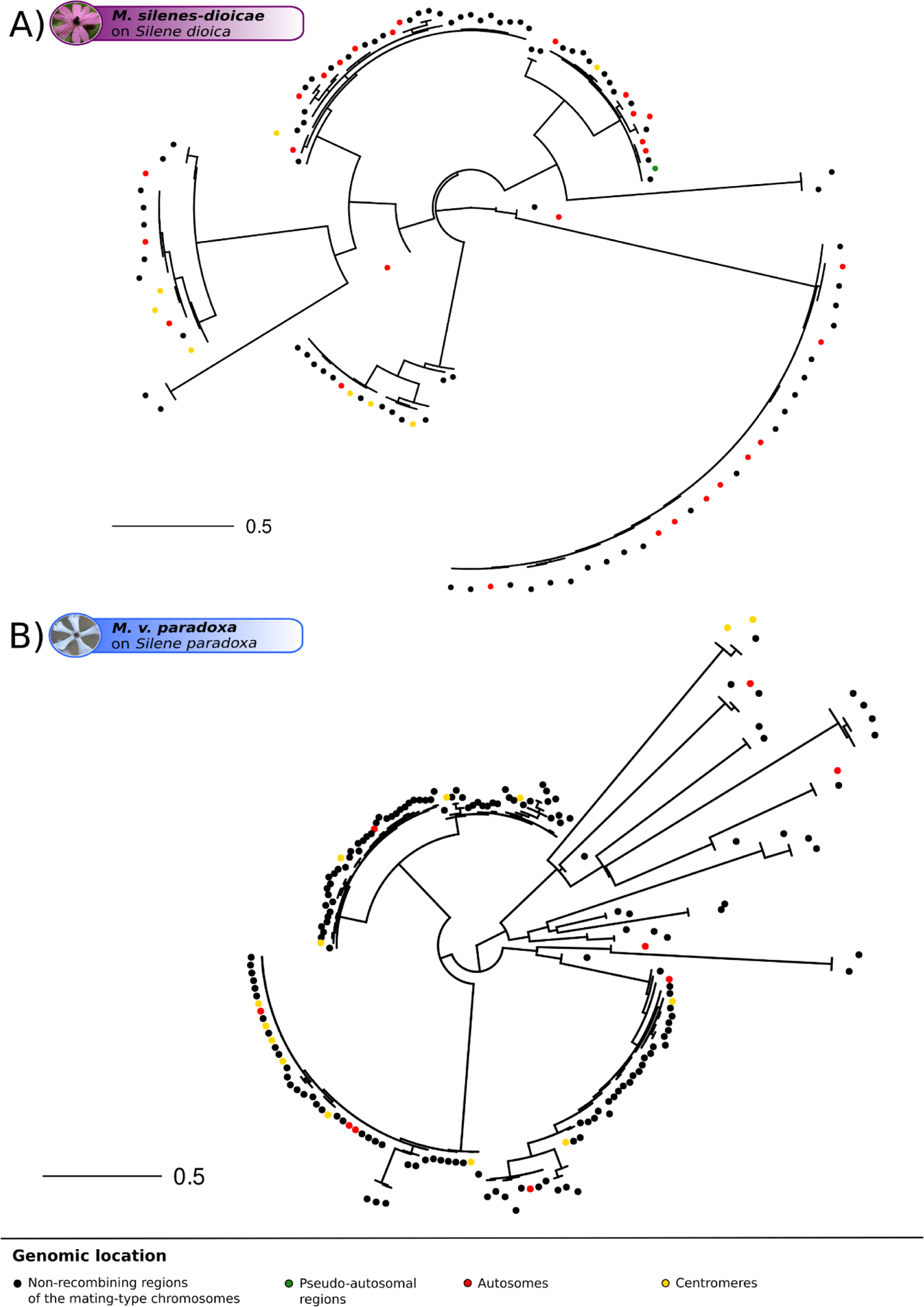
Single-species genealogies of *Ty3* retroelement copies in *Microbotryum* genomes. Single-species genealogies of *Ty3* copies in *Microbotryum silenes-dioicae* **(A)** and *M. v. paradoxa* **(B)**. The color of the dots at the tip of the branches corresponds to the genomic location of the transposable element (TE) copies.

**Supplementary Figure S11:**
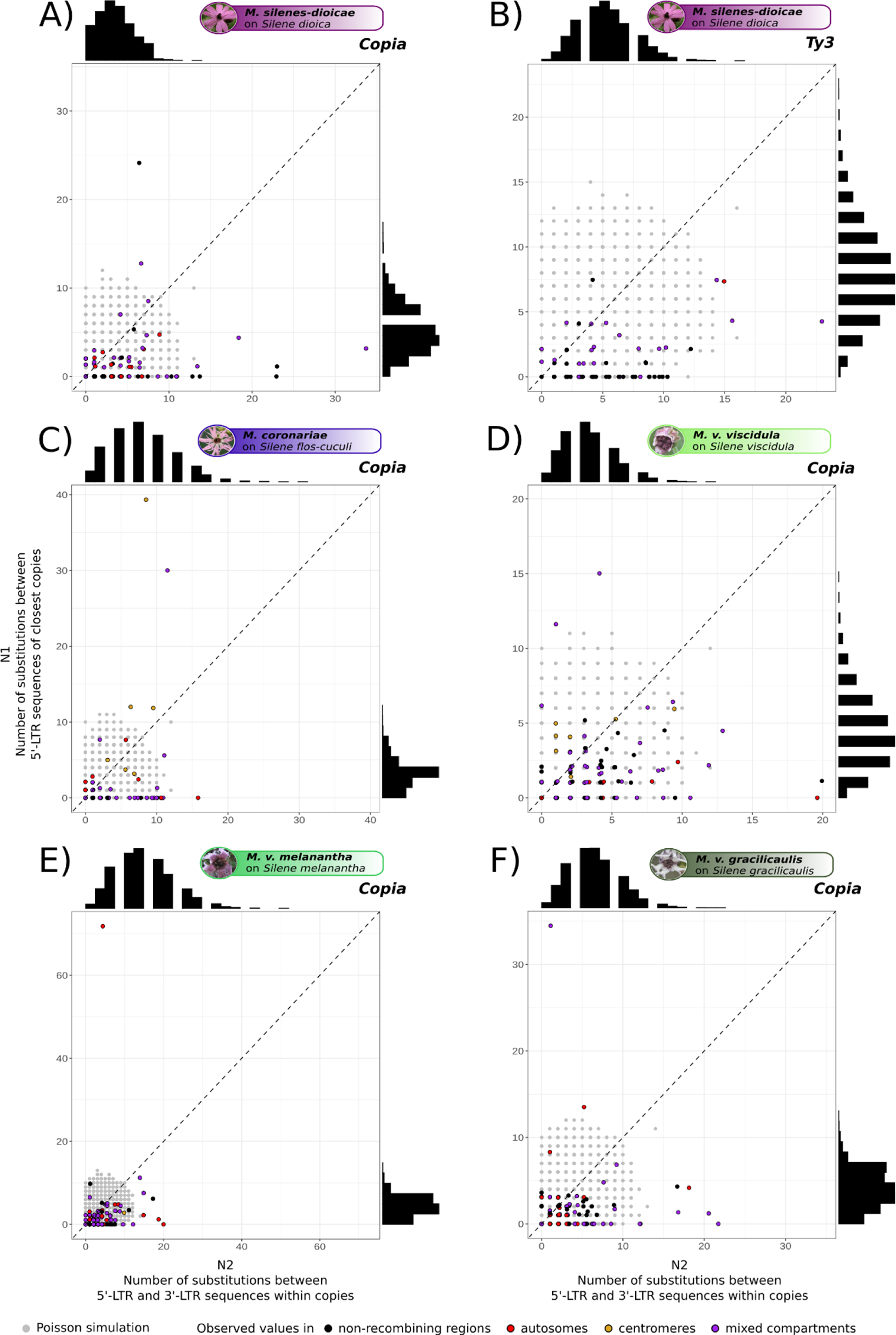
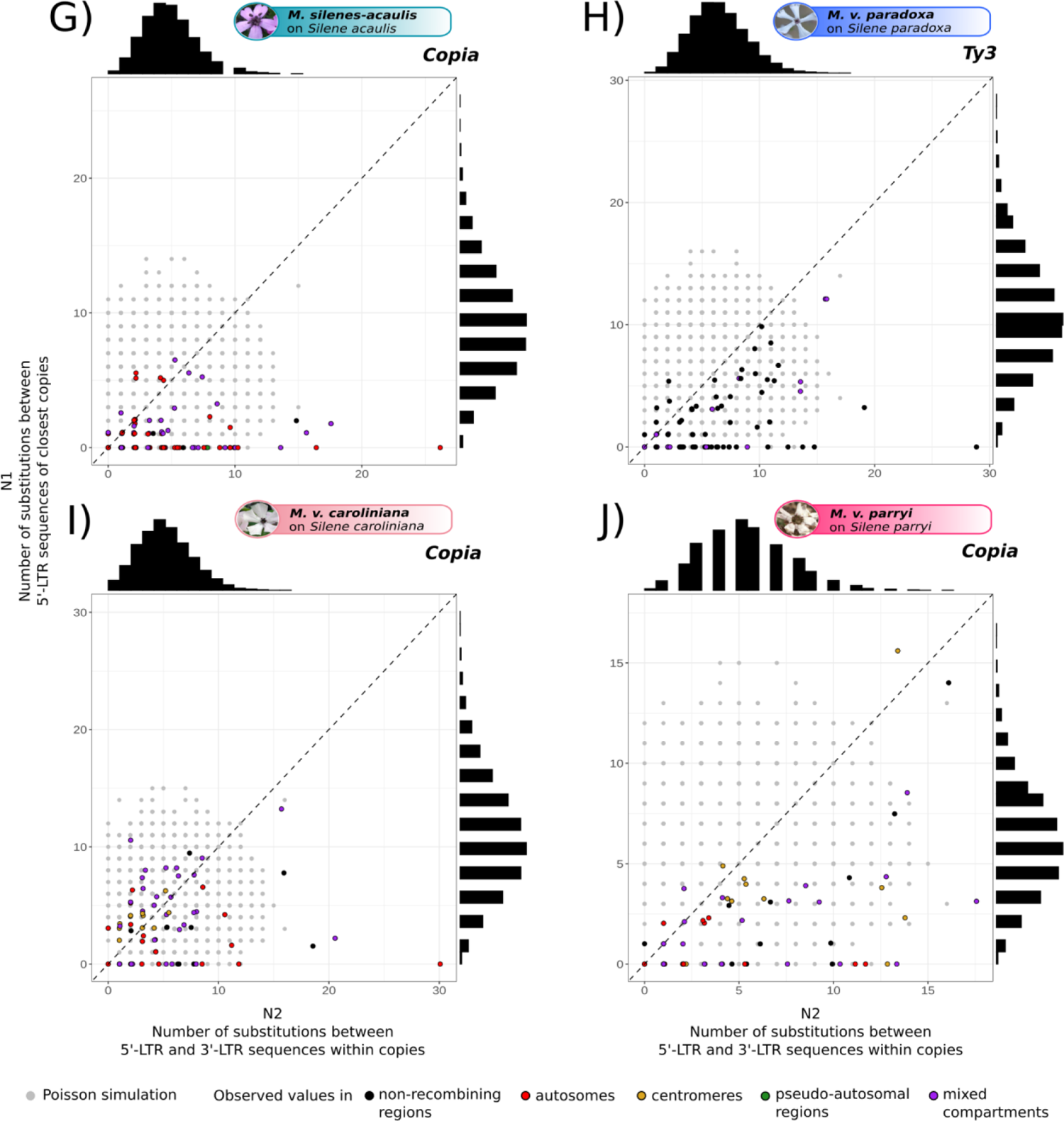
Investigation of the possible occurrence of conversion events within clusters of highly similar LTR-retrotransposon copies in *Microbotryum* fungi. We looked for footprints of conversion events as explained on Figure 5 and in the method section, for the *Copia* (A) and *Ty3* (B) retrotransposon copies of *Microbotryum silenes-dioicae*, for the *Copia* retrotransposons of *M. coronariae* (C), *M. violaceum viscidula* (D), *M. v. melanantha* (E), *M. v. gracilicaulis* (F), *M. silenes-acaulis* (G), for the *Ty3* retrotransposon copies of *M. v. paradoxa* (H), and for the *Copia* retrotransposon copies of *M. v. caroliniana* (I) and *M. v. parryi* (J). In colors are shown the number of substitutions between the 5’-LTR sequences of pairs of most similar copies (N2) within clusters of highly similar retroelement copies as a function of the number of substitutions between the 5’- and 3’-LTR sequences within copies (N1). The Poisson simulated numbers of substitutions (with lambda equal to the average number of substitutions between the 5’- and 3’-LTR sequences within copies) are plotted in gray. The colors of the points indicate the genomic location of the copy pairs. The observed distributions of the number of substitutions between 5’-LTR sequences and between 5’ - 3’ LTR sequences are shown at the top and at the right of each scatterplot.

**Supplementary Figure S12:**
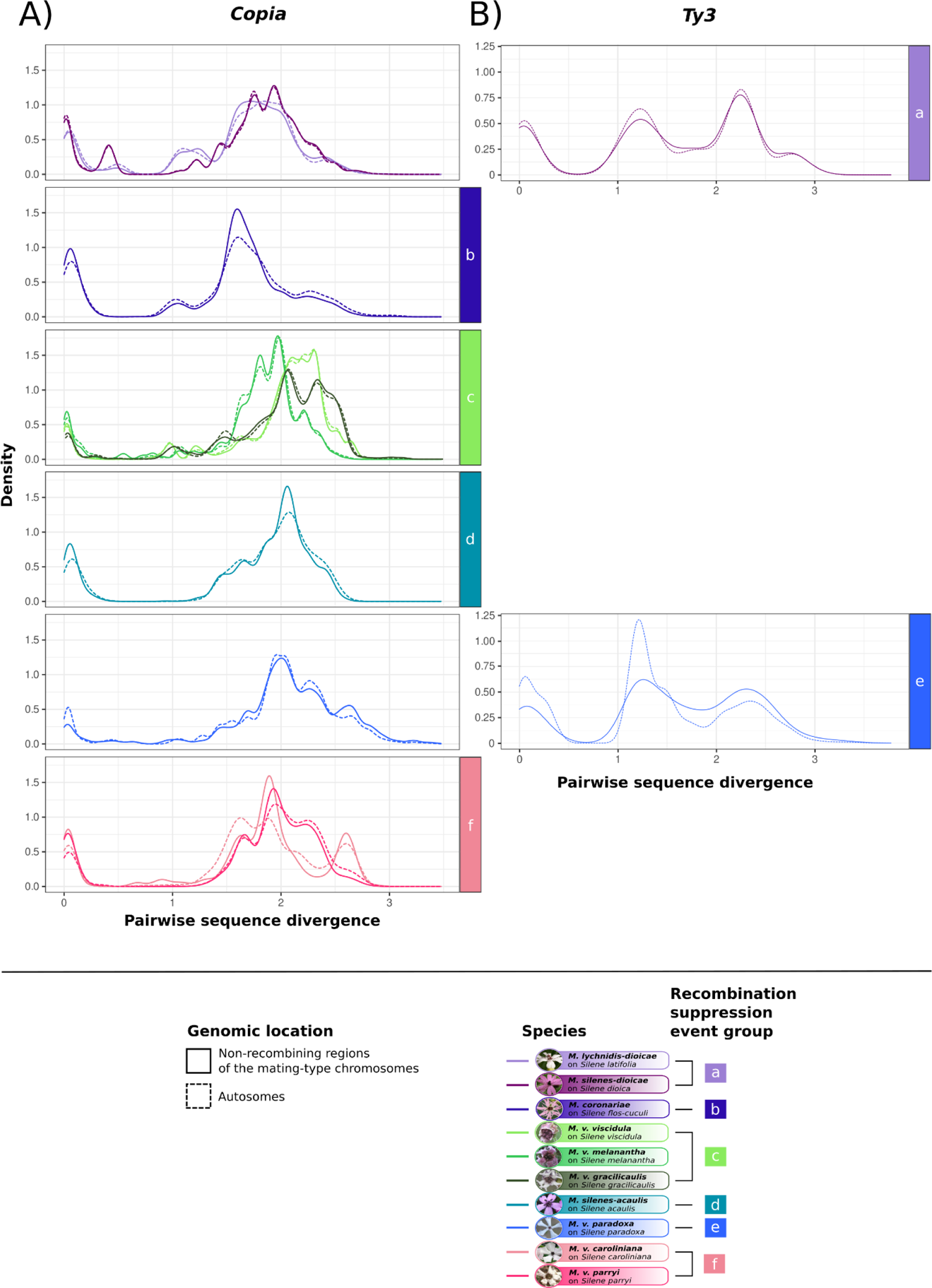
Distribution of pairwise divergence for retroelements in the non-recombining regions of the mating-type chromosomes and the autosomes in each species, plotted separately for the different independent events of recombination suppression. **(A)** *Copia* retrotransposons, **(B)** *Ty3* retrotransposons.

**Supplementary Figure S13:**
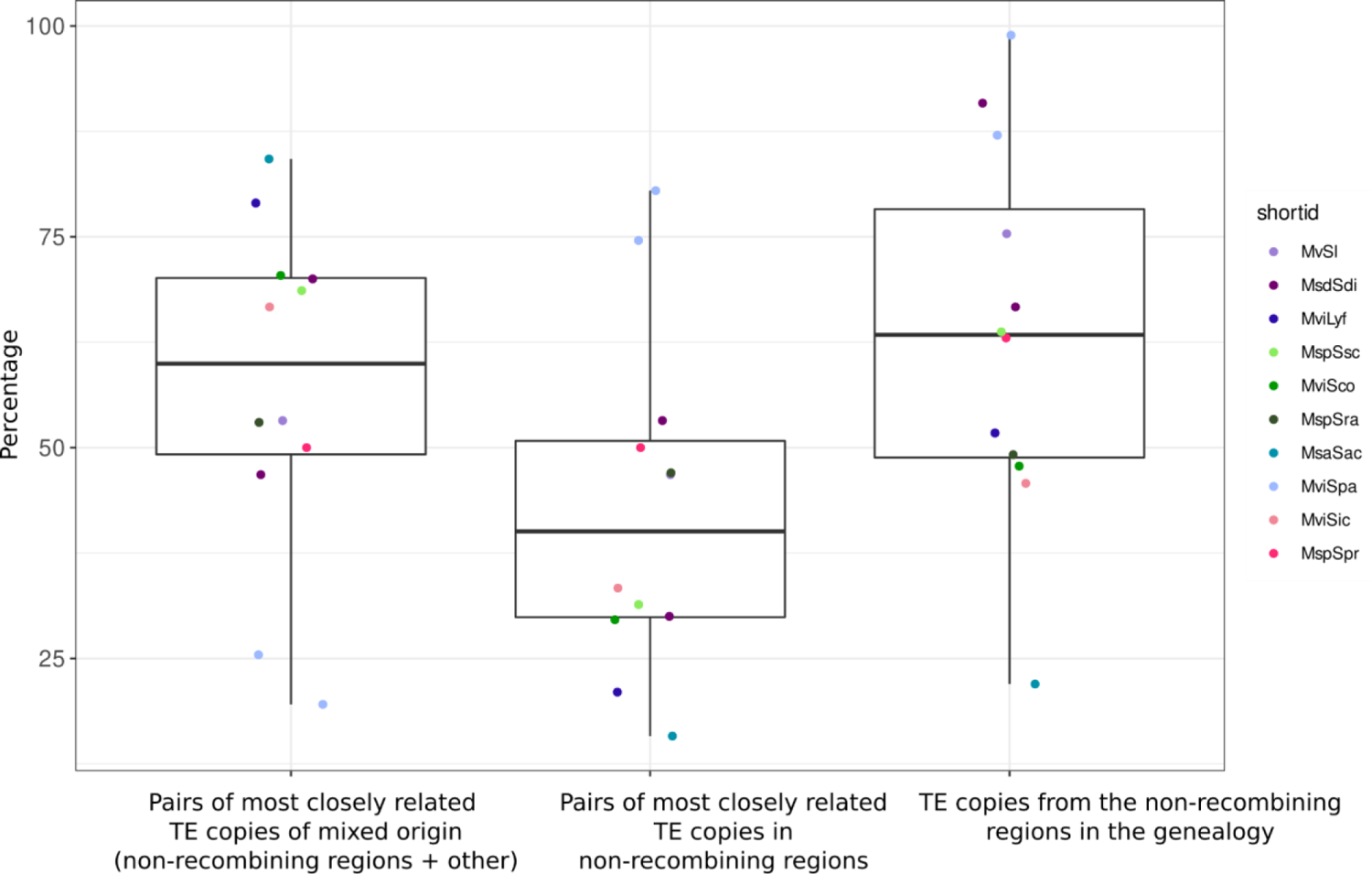
Percentage of pairs of most closely related long-terminal repeat (LTR) sequences with at least one copy in the non-recombining regions compared to their representation in their transposable element (TE) genealogy. Each colored point corresponds to a TE genealogy (*Copia* or *Ty3*) of a *Microbotryum* species (Figure 4 and Supplementary Figure S9 and S10), according to the species color code from Figure 1. From left to right are represented the percentages of pairs of most closely related copies i) involving one copy from the non-recombining regions and from another location (*e.g.*, the autosomes), ii) with both copies in the non-recombining regions, and the percentage of copies located in the non-recombining regions, in their TE genealogy.

**Supplementary Figure S14:**
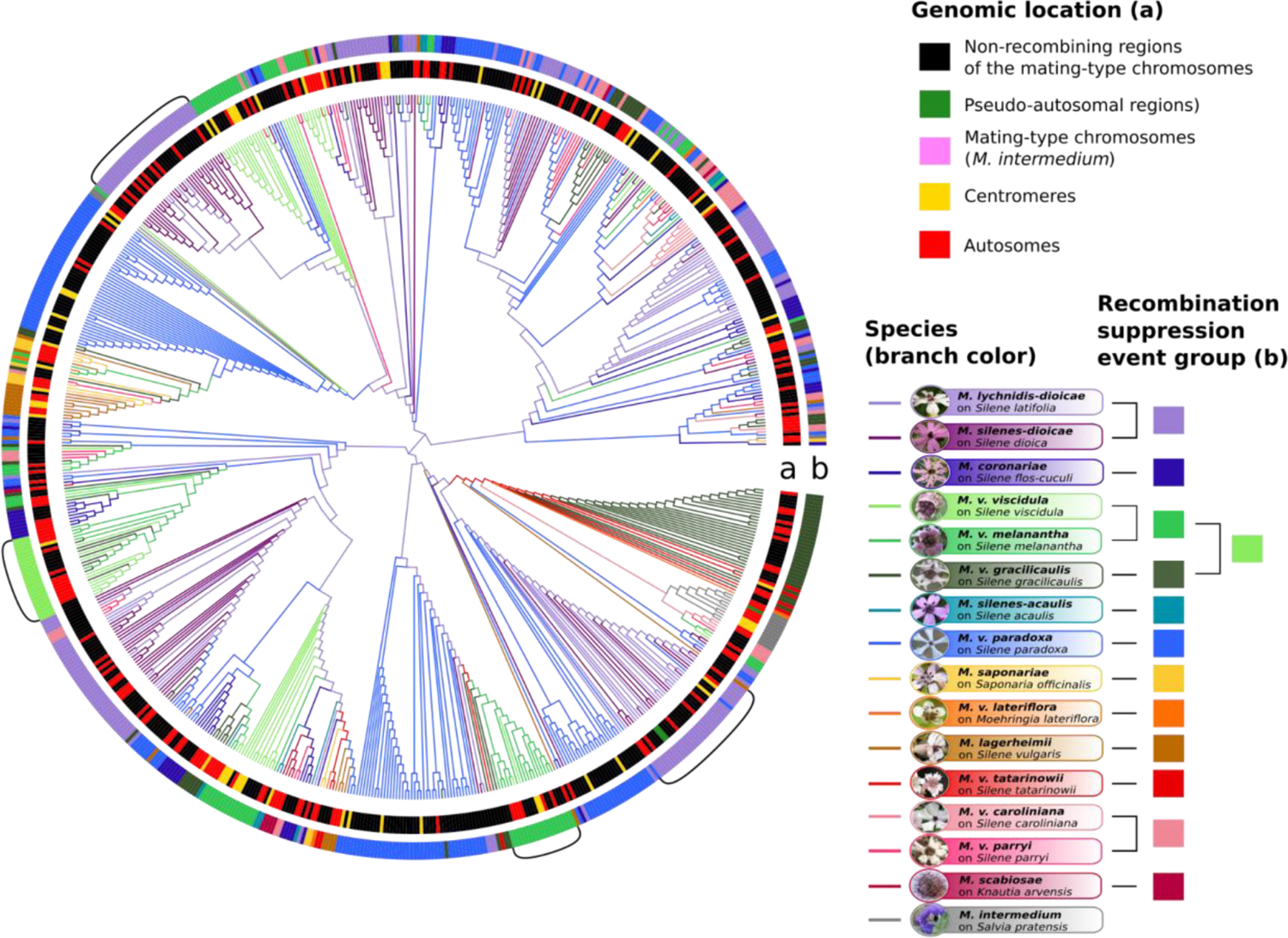
Multi-species genealogies of *Ty3* retroelement copies in *Microbotryum* genomes. Multi-species *Ty3* trees of all *Microbotryum* species analysed in this study. The branch color corresponds to the species, the first inner track (a) corresponds to the genomic location of the transposable element (TE) copies, the second outer track (b) corresponds to the linkage event group of the species carrying the TE copy. Brackets highlight bursts of TEs.

**Supplementary Figure S15:**
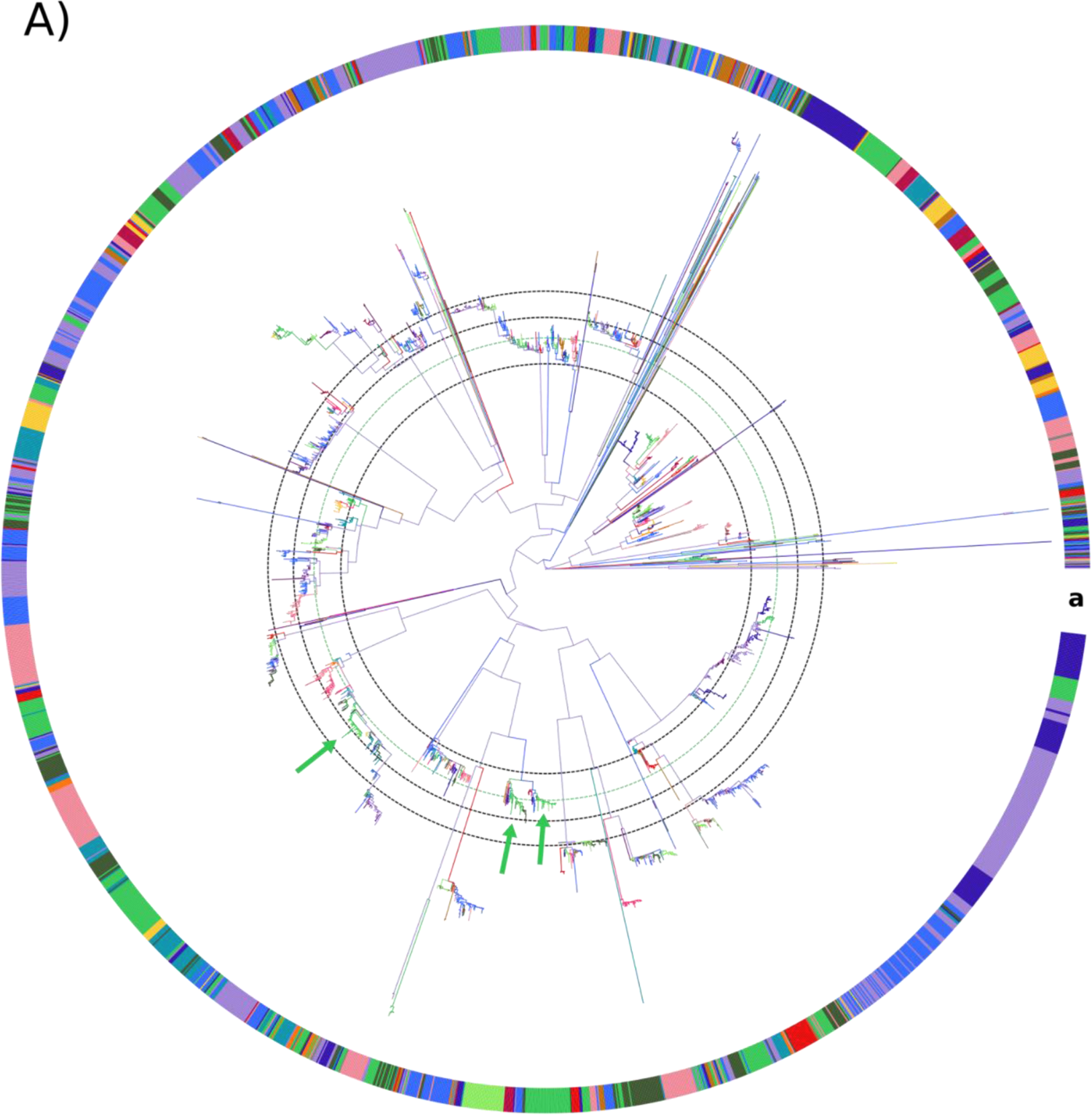
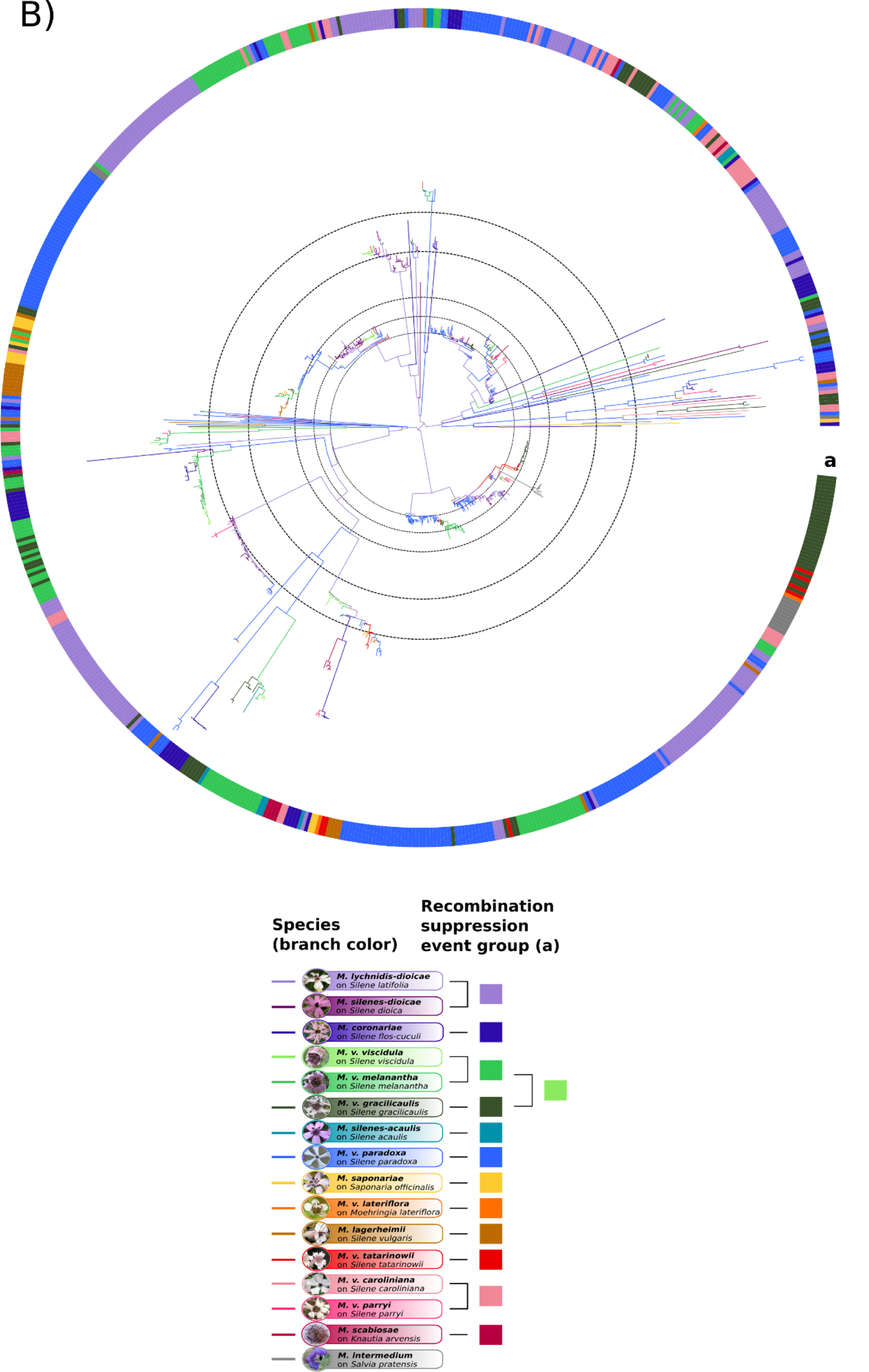
Multi-species genealogies of retroelement copies in *Microbotryum* genomes. The phylogenies of *Copia* **(A)** and *Ty3* **(B)** are the same as those displayed on Figure 5 and Supplementary Figure S14, respectively, but here plotted with branch lengths. The branch color corresponds to the species, the track (a) corresponds to the linkage event group of the species carrying the TE copy. The dashed circles are a help to visualize the timing of the bursts. The green dashed circle highlights the *Copia* bursts which are almost synchronized in the clade of *M. v. gracilicaulis*, pointed by green arrows.

**Supplementary Figure S16:**
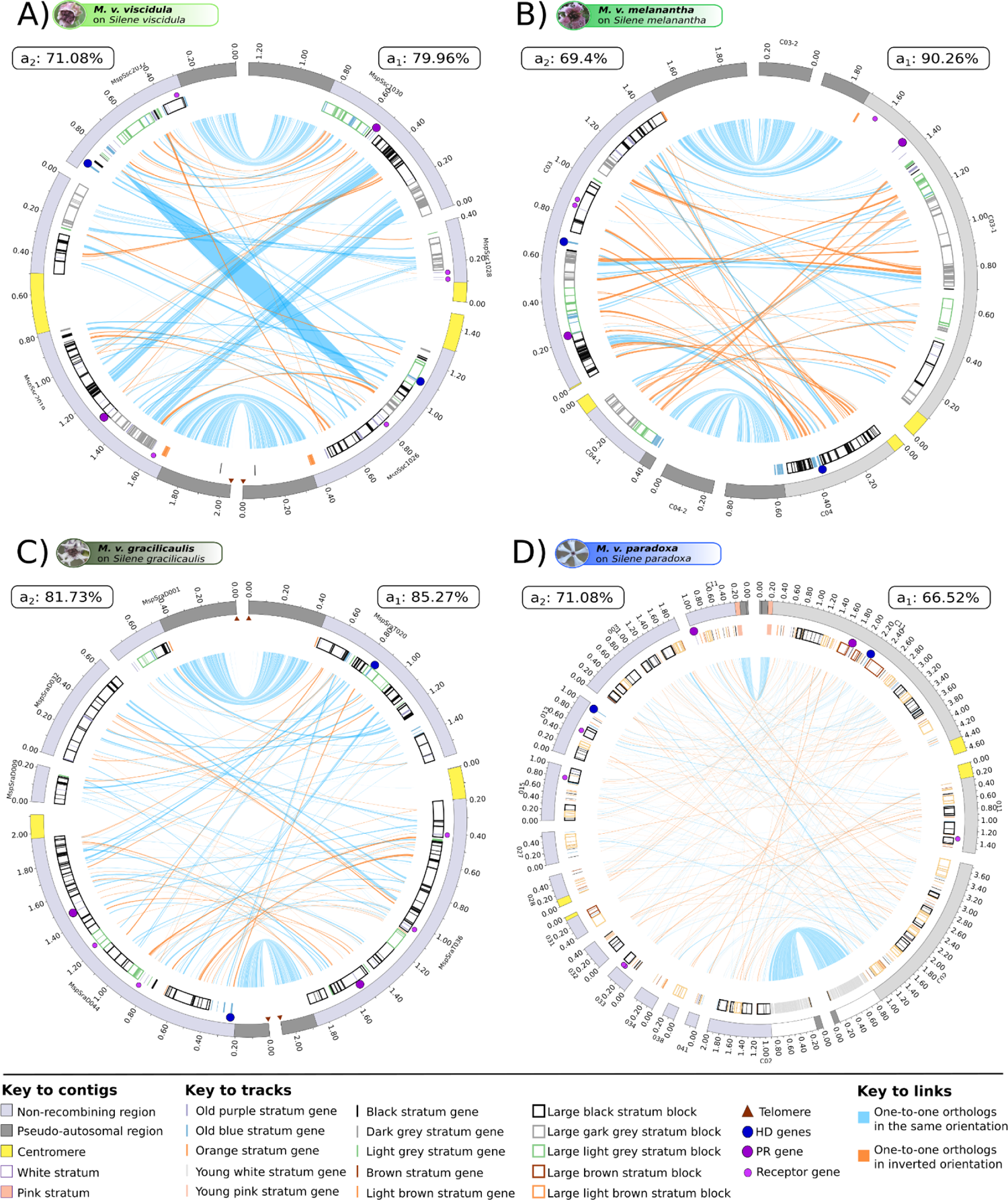
Delimitation of blocks for the rearranged evolutionary strata in mating-type chromosomes. The PR, HD and pheromone genes are represented by purple, blue and small light purple circles, respectively. The outer track represents contigs, with length ticks every 200 kb and centromeres indicated in yellow. Non-recombining regions are displayed in dark grey and pseudo-autosomal regions in light gray. The young, non-rearranged white and pink strata of *M. v. paradoxa* are highlighted on the outer track. Genes from the different strata are displayed by bars on the inner track (see Figure 1 for the stratum formation history). Large blocks (greater than 80 kb) of genes from a single stratum are indicated on the inner track by rectangles. The light grey stratum is displayed in green to facilitate visualization. The percentage of non-recombining regions retained in the blocks of strata are indicated next to the mating type, on each side of the circos plots. Blue and orange lines link alleles, the latter corresponding to inversions. The link width is proportional to the corresponding gene length.

**Supplementary Figure S17:**
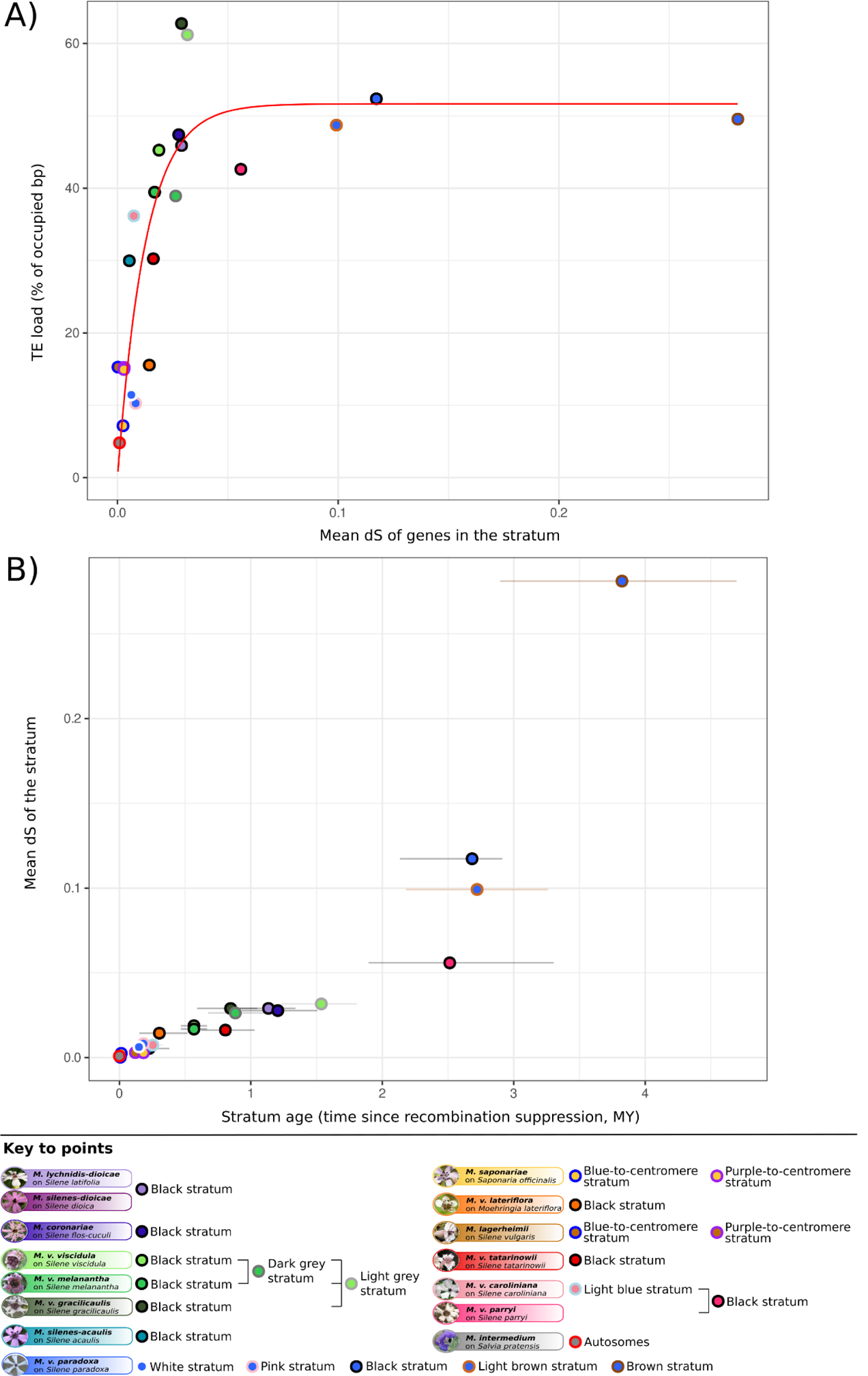
Relationship between synonymous divergence (d_S_), transposable element (TE) load and stratum age in *Microbotryum* mating-type chromosomes. A) Percentage of base pairs occupied by TEs in non-recombining regions (means in a_1_ and a_2_ genomes and means across species sharing the same strata) as a function of the mean d_S_ of the genes present in the stratum. The red curve shows the prediction of the best model corresponding to a negative exponential model. B) Relationship between the stratum age in million years (MY) and the mean d_S_ of the genes contained in this stratum. Each dot corresponds to an independent evolutionary stratum in A) and B).

**Supplementary Figure S18:**
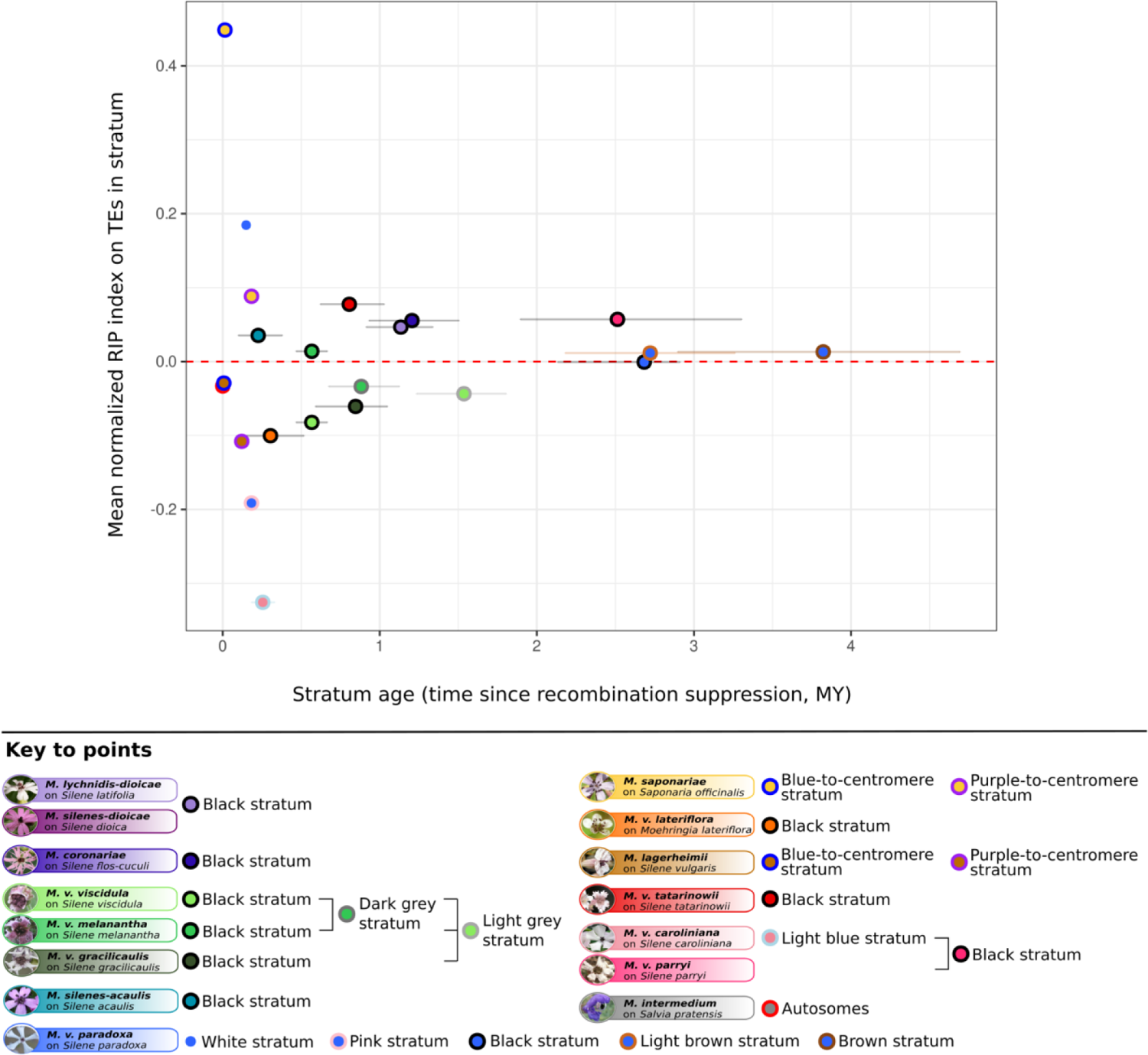
Relationship between the abundance of transposable element (TE) inactivation footprints by repeat-induced point mutation (RIP) in the non-recombining regions of the *Microbotryum* mating-type chromosomes and time since recombination suppression in million years (MY). Mean normalized RIP index on TEs in non-recombining regions as a function of their age with confidence intervals. The red dashed line separates the RIP index values suggesting RIP mutations (> 0) or no RIP activity (≤ 0). The two-sided Pearson correlation test was not significant (*p-value* = 0.9).

**Supplementary Figure S19:**
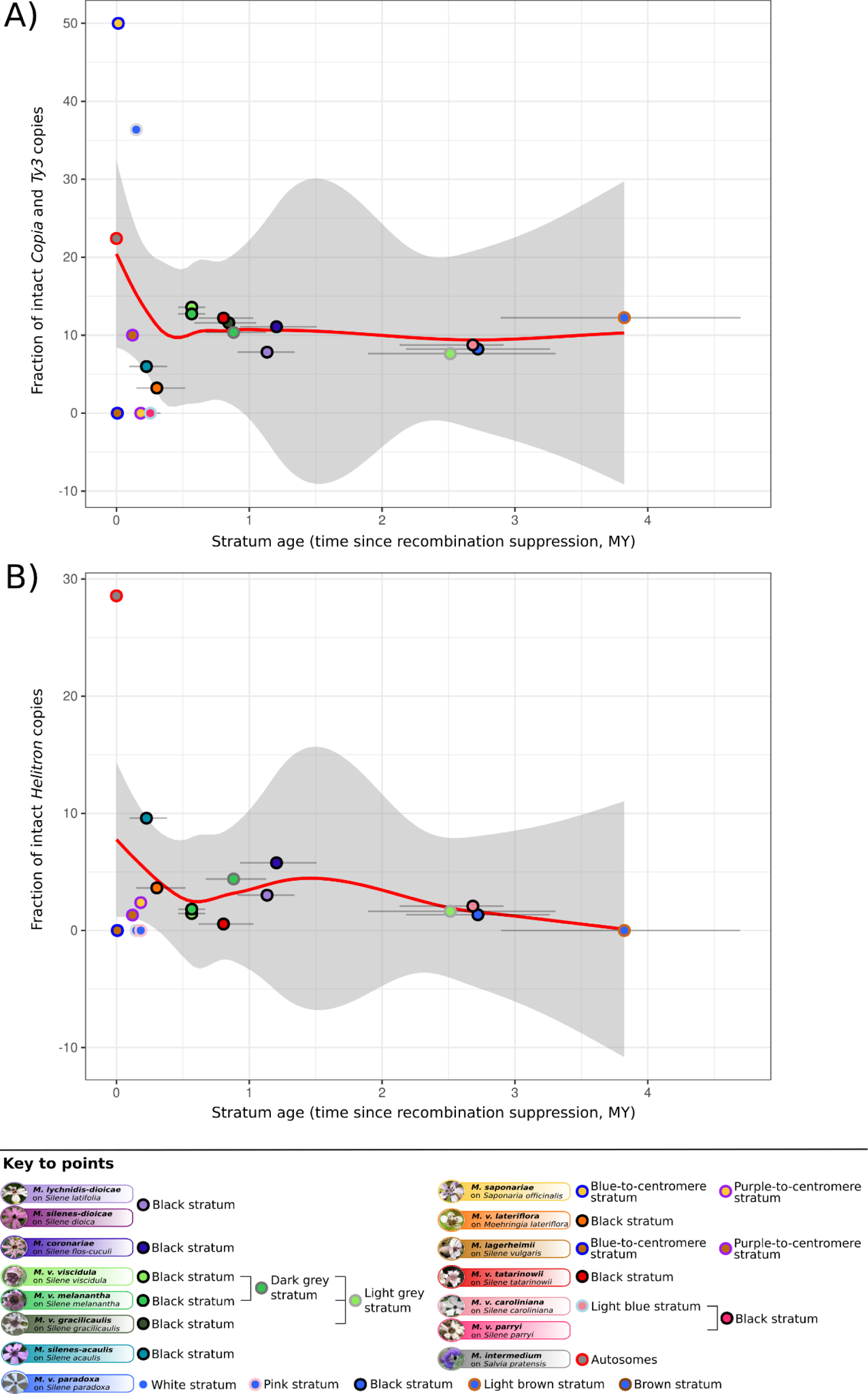
Relationship between the fraction of intact *Copia, Ty3* and *Helitron* copies and stratum age in *Microbotryum* mating-type chromosomes. Fraction of intact *Copia* and *Ty3* copies (5-7.5 kb for intact copies) (A) and *Helitron* copies (5-11 kb for intact copies) (B) in non-recombining regions (means in a_1_ and a_2_ genomes and means across species sharing the same strata) as a function of the time since recombination suppression in million years (MY). Each dot corresponds to an independent evolutionary stratum in A) and B). The red curve shows the smooth curve obtained by local regression.

## Notes

### Competing Interest Statement

The authors have declared no competing interest.

### Summary of Updates

Test for conversion events in the TE genealogies added; Figure 5, S11 and S13 added; Figures 3, 4, 6, S6, S8, S9, S10, S12, S14, S15 revised; manuscript revised.

